# Simultaneous Isolation and Characterization of Lipoprotein Classes in Plasma, Including HDL Subclasses and the Uncharacterized Dense HDL

**DOI:** 10.1101/2025.09.29.679308

**Authors:** Joanne Agus, Olesia Gololobova, Jack Zheng, Armin Oloumi, Xinyu Tang, Brian Hong, Susan Lei, Andrey Turchinovich, Wyatt Vreeland, Carlito Lebrilla, Juan Pablo Tosar, Kenneth Witwer, Angela M. Zivkovic

**Affiliations:** Department of Nutrition, University of California Davis, California, United States of America; Molecular and Comparative Pathobiology, John Hopkins University School of Medicine, Maryland, United States of America; Department of Chemistry, University of California Davis, California, United States of America; Heidelberg Biolabs, Heidelberg, Germany; Bioprocess Measurement Group, National Institute of Standards and Technology, Maryland, United states of America; Functional Genomics Unit, Institut Pasteur de Montevideo, Montevideo, Uruguay; Faculty of Science, Nuclear Research Center, Analytical Biochemistry Unit, Universidad de la República, Montevideo, Uruguay

**Keywords:** Lipid nanoparticles, extracellular vesicles, liquid biopsy, high density lipoprotein, dense high density lipoprotein

## Abstract

Lipoproteins (LPP) and extracellular vesicles are carriers of extracellular small RNA, with potential applications both in the areas of diagnostics and therapeutics. Lipid nanoparticles overlap across a range of densities and sizes in plasma, making them difficult to isolate intact and without contamination from other plasma components. Accurate characterization of their cargo through efficient isolation from other plasma components is required to understand their function. Here we describe the simultaneous separation of LPP classes using sequential flotation ultracentrifugation followed by size exclusion chromatography from 500µL of starting plasma. Using western blot, denaturing and non-denaturing gel electrophoresis, nuclear magnetic resonance, and electron microscopy, we demonstrate separation of the LPP classes with minimal contamination. We also show unique lipidomic, proteomic and small RNA signatures for each LPP class, including, for the first time, very high density high density lipoprotein particles in the density range of 1.21–1.25 g/mL.

**Motivation:** In plasma, lipid nanoparticles overlap in size and density, making it difficult to separate and isolate intact particles. Additionally, currently available methods do not allow for simultaneous characterization and isolation of these particles. This method resolves both challenges. The results presented also analyze and describe the presence of known lipoprotein particles, as well as those that have been previously under-characterized.

## Introduction

Lipoproteins (LPP) and extracellular vesicles (EVs) are the major vesicular transporters of extracellular ribonucleic acid (RNA) in plasma^1^. Interestingly, LPP outnumber EVs by six to seven orders of magnitude in plasma^2,3^, and yet they are understudied for their role in RNA transport, and their potential utility for diagnostics and therapeutics is underexplored. This is particularly true for high-density lipoproteins (HDL), which have been found to transport small RNA^4^ but which remain understudied due to the high heterogeneity of HDL subclasses, and their extensive overlap in terms of both density and size with other LPP, EVs, and other plasma components. LPP, including the triglyceride-rich lipoprotein particles (TRLP) (intestinally derived chylomicrons, liver derived very low-density lipoproteins (VLDL), and their remnants), low-density lipoproteins (LDL), and HDL, are widely known for their roles in the transport of lipids such as cholesterol and phospholipids, as well as other lipophilic molecules, yet they also play a prominent role in regulating inflammation and immune function^5,6^. The roles of HDL particles in immunoregulation^6^ and their ability to deliver small RNA to and from a wide range of tissues^4^ have been described. However, HDL are a highly heterogeneous class of LPP with particles ranging in size from 7 nm to 15 nm in diameter, ranging in density from 1.063 g/mL to 1.21 g/mL, and with as many as 16 possible subclasses based on their protein composition^7^. The RNA content of the HDL subclasses has not been fully described. Moreover, the existence of very high-density HDL (VHDL) particles, in the density range of 1.21 g/mL–1.25 g/mL, was first identified in 1966^8^ though these particles have remained largely uncharacterized, despite the fact that they may represent as much as 15%-20% of total HDL in circulation^8^. The RNA content of these VHDL particles, as well as their comprehensive proteomic and lipidomic composition have never been previously described.

For the most part, methods to isolate and study LPP and EVs are specialized for the isolation of a single target nanoparticle class, including, for example, antibody-based methods to extract only those nanoparticles that carry a specific protein or set of proteins^1,9^. Current approaches to simultaneously isolate all LPP classes, including the HDL subclasses, have significant limitations. For example, traditional density gradient-based ultracentrifugation (UC) methods require long centrifugation times (24-48 hours) and large sample volumes (1 mL-5 mL), and the density gradients can be difficult to generate and maintain^10^. The protocols for many published LPP isolation methods also require steps that are not amenable to throughput, for example, protocols that require piercing the centrifuge tube with a syringe to extract the target fraction from the bottom of the tube^11^. These, among other complications, highlight the need for a method to simultaneously isolate the LPP fractions, including HDL subclasses, that requires a small sample volume (500µL), and can comprehensively capture the heterogeneity of these particles while conserving their functionality.

We have previously shown that a method for isolating HDL using density-based sequential UC followed by size exclusion chromatography separation (UC-SEC) from a starting volume of 500µL of plasma^12^ yields adequate amounts of isolated HDL particles to run multiple functional assays, which was capable of distinguishing clinical diagnoses and lifestyle differences^13,14^. The addition of optical detectors, including multi-angle light scattering (MALS), dynamic light scattering (DLS), and differential refractive index (dRI), further makes it possible to simultaneously characterize the particle diameter and molecular weight of the particles as they are being isolated.

In this paper, we demonstrate the utility of the UC-SEC-UV/MALS/DLS/dRI method for the isolation of all LPP classes in plasma, and the simultaneous characterization of HDL particle diameter and molecular weight. We also characterize the major differences in composition of all LPP fractions, including the HDL subfractions, by measuring the lipidome, proteome, and RNA content of these particles and whole particle sizing by non-denaturing gel electrophoresis (NDGE). We also characterize, for the first time, the composition of VHDL, which were first identified in 1966^8^, but whose basic characteristics have not yet been described. This includes particles between the densities of 1.21 g/mL and 1.25 g/mL, which includes dense low-density lipoprotein (dLDL), and dense HDL large, medium, and small (dHDL-L, dHDL-M, dHDL-S). Additionally, we also characterize the non-dense LPP particles, which includes TRLP, intermediate density lipoprotein (IDL), LDL, HDL large, medium, and small (HDL-L, HDL-M, HDL-S), and albumin (Alb) – all particles between the densities of 1.006 g/mL and 1.21 g/mL.

## Method

### 1. Blood Processing

Plasma samples were from 20 healthy participants, 10 males and 10 pregnant females. Samples were obtained from the University of California San Diego^15^. Whole blood processing protocols were reported previously^15^. Participants were consented and samples were obtained under an approved Institutional Review Board protocol by the Human Research Protection Programs at the University of California, San Diego following the ethical standards set by the Helsinki declaration. Equivalent volumes of plasma from male and pregnant female participants were thawed on ice, then pooled, and centrifuging to further remove platelets at 10, 000 RPM for 30 minutes using the Sorvall Refrigerated Benchtop Centrifuge, Legend X1R (Thermo Fisher Scientific, USA).

Quality control (QC) plasma was obtained from 3 healthy male participants after an overnight fast in EDTA tubes, and centrifuged at 1500 x *g* for 10 minutes using Sorvall Legend RT Refrigerated Benchtop Centrifuge (Thermo Fisher Scientific, USA). The participants were recruited from the University of California, Davis and blood draws were performed at the Ragle Human Nutrition Research Center, in the Department of Nutrition. Samples were obtained under a protocol approved by the University of California Davis, Institutional Review Board and followed all ethical standards set by the Helsinki declaration. Plasma was extracted, aliquoted and stored at −80°C. The QC plasma pool was generated by thawing aliquoted plasma from each participant on ice, then pooling, and centrifuging to further remove platelets at 10, 000 RPM for 30 minutes using the Sorvall Refrigerated Benchtop Centrifuge, Legend X1R (Thermo Fisher Scientific, USA). Pooled QC plasma was then re-aliquoted into 500µL aliquots and stored at - 80°C until further analysis. Plasma pool sample characteristics can be found in Supplemental Table S1.

### 2. UC-SEC-UV/MALS/DLS/dRI

A previously published method^12^ for isolating HDL particles was modified to enable the isolation of additional LPP classes and additional HDL subclasses (Figure 1). First, 500µL of plasma was underlaid in a 4.7 mL OptiSeal tube (Beckman-Coulter, IN, USA) with 1.0035 g/mL density solution of potassium bromide (KBr, Sigma Aldrich, ACS Reagent ≥ 99%) to achieve a final density of 1.006 g/mL. The sample was subjected to UC using an Optima Max-TL (Beckman-Coutler, IN, USA) and fixed angle TLA-110 rotor (28°, k factor 13, Beckman-Coulter, IN, USA) for 4 hours at 110,000 RPM (656241.5 x *g*). Following the first UC step, the top 4 mL was collected as the TRLP fraction, and the bottom 700µL was collected and balanced with 1.1 mL of 1.34 g/mL KBr to form 1.8 mL of solution at a final density of 1.21 g/mL. The solution was then further underlaid in another 4.7 mL Optiseal tube with 2.9 mL of KBr solution with a density of 1.21 g/mL and subjected to UC for 12 hours at 58,000 RPM (182446x *g*). The top 2 mL of the density-separated solution in the Optiseal was collected and concentrated using a 50 kDa Amicon filter (Ultracell-50, Sigma Aldrich, IN, USA) to 250µL before injection into a high performance liquid chromatography instrument (HPLC, 1260 Infinity, Agilent) with a 24 mL sepharose, size exclusion column (SEC, Superdex 200 Increase 10/300, Cytiva) at a flow rate of 0.4 mL per minute. The eluting fractions included IDL, LDL, HDL-L, HDL-M and HDL-S. The HDL fractions were further analyzed for particle molecular weight (Mw) and hydrodynamic radius (Rh) using the MALS (miniDAWN, Waters-Wyatt Technologies, CA, USA) and DLS (WyattQELS, Waters-Wyatt Technologies, CA, USA) detectors respectively, and the dRI detector (rEX, Waters-Wyatt Technologies, CA, USA) to measure the total mass of the particles.

**Figure 1.**
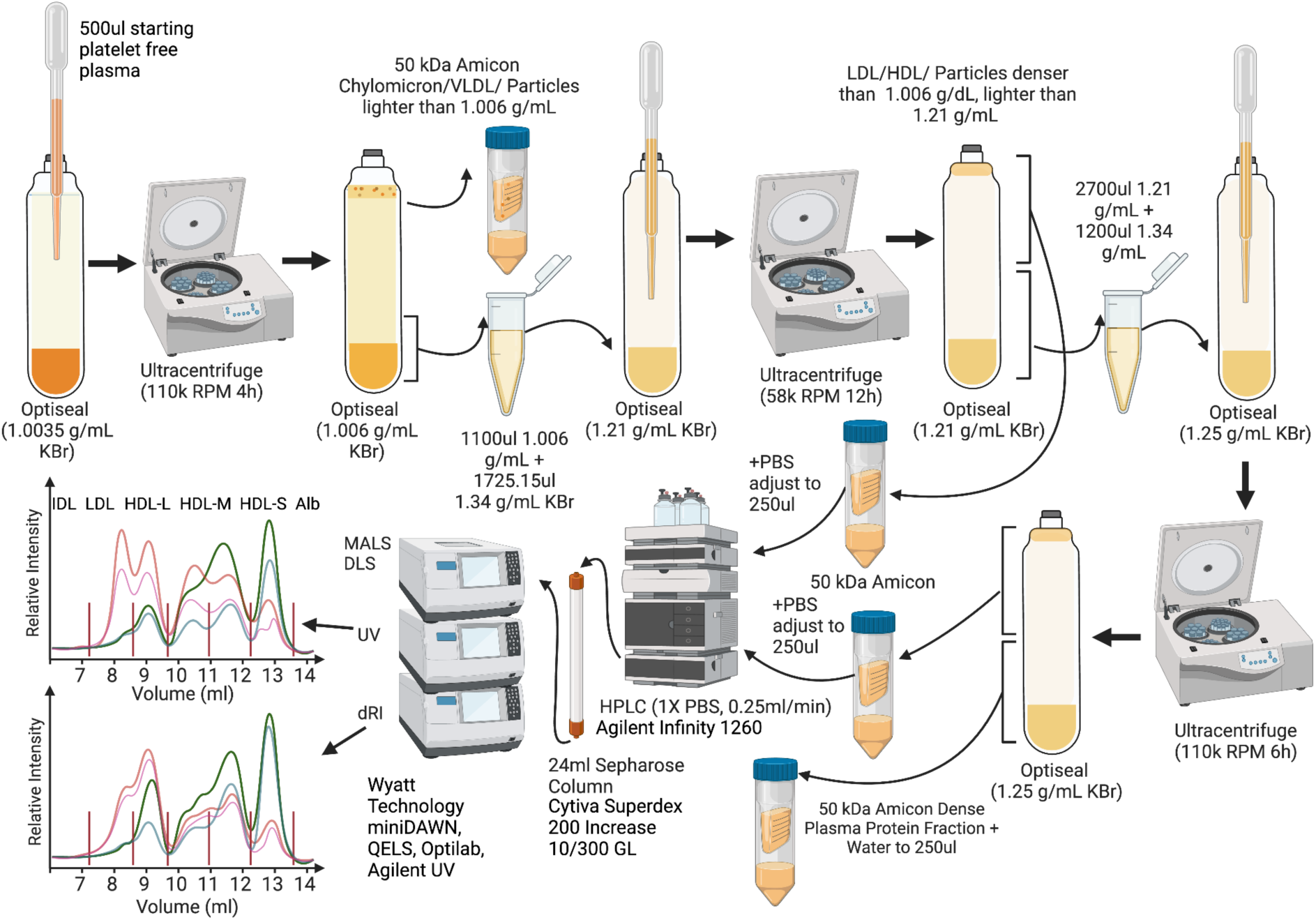
Diagram of lipid nanoparticle isolation work flow using ultracentrifugation (UC) followed by size exclusion chromatography (SEC) from a starting plasma sample of 500ul. The process includes three UC steps: First, to float particles lighter than 1.006 g/mL in density. Second, to float particles between 1.006 g/mL and 1.21 g/mL in density. Third, to float particles between 1.21 g/mL and 1.25 g/mL in density. The method also includes two high performance liquid chromatography (HPLC) injections to separate particles using an SEC column after the second (1.006 g/mL and 1.21 g/mL) and third (1.21 g/mL and 1.25 g/mL) UC steps. SEC separation after the second UC step yields seven fractions, which includes intermediate density lipoprotein (IDL), low density lipoprotein (LDL), high density lipoprotein large (HDL-L), HDL-medium (HDL-M), HDL-small (HDL-S) and Albumin. SEC separation after the third UC step yields 4 fractions, which includes dense-LDL (dLDL), dense-HDL-L (dHDL-L), dense-HDL-M (dHDL-M), and dense-HDL-S (dHDL-S). During the SEC isolation, samples were analyzed for size (molecular weight and hydrodynamic radius), protein content, and total mass, using in-line multi-angle light scattering (MALS), dynamic light scattering (DLS), ultraviolet (UV) at 280 nm, and differential refractive index (RI) detectors.

Following the second UC step, the bottom 2.7 mL of the 1.21 g/mL solution was added to 1.2 mL of 1.34 g/mL KBr solution to form a 3.9 mL solution with a final density of 1.25 g/mL. The 3.9 mL of 1.25 g/mL solution was then underlaid in a 4.7 mL Optiseal tube with 0.8 mL of KBr solution with a density of 1.25 g/mL. A third UC step was conducted for 6 hours at 110,000 RPM. The top 2 mL of the solution in the OptiSeal was collected and concentrated using a 50 kDa Amicon filter to 250µL. The solution was then injected into the SEC-UV/MALS/DLS/dRI system for separation by size and particle analysis to obtain the following fractions using the same size cutoffs as for the 1.21 g/mL density fraction: dLDL, dHDL-L, dHDL-M, and dHDL-S.

Samples were fractionated based on previously determined analyses, which includes peak-to-through differentiations, reducing gel-electrophoresis, western blot (WB), and nuclear magnetic resonance (NMR) LipoProfile during the method development period. Fractions were taken off from the temperature controlled (10°C) fraction collector (Infinity 1260, Agilent) and concentrated using a 50 kDA Amicon filter to 100µL and stored in 2% sucrose. Quality control samples were analyzed using 500µL of another plasma pool using the same isolation method every 8th sample. The HPLC and optical detectors were operated using HPLC Connect 2.0.1.35 (Agilent Technologies, CA, USA) and Astra 8.1.2 (Waters-Wyatt Technologies, CA, USA) respectively.

#### Molecular Weight and Hydrodynamic Radius Determination

The Mw of each lipoprotein fraction was determined using the protein conjugate method in Wyatt Astra 8.1.2 software. The weighted average protein extinction coefficient was calculated from amino acid sequence-derived values using ProtParam and the formula from Gill and von Hippel (1989) ^16^ (Supplemental Table S2 and S3). These values were then weighted based on proteomic composition (Supplemental Table S4) to obtain the weighted average extinction coefficient. Protein concentration was determined from UV absorbance at 280 nm. The weight-average Mw of each lipoprotein fraction, as well as individual protein and lipid component masses, was determined using MALS and dRI. A miniDAWN (Wyatt Technology, Goleta, CA) MALS detector (λ = 662.5 nm ± 4 nm) was used for light scattering measurements, while an Optilab (Wyatt Technology, Goleta, CA) dRI detector was used for concentration measurements. The refractive index increment (dn/dc) values used were 0.1899 mL/g for proteins^17^ and 0.1605 mL/g for lipids^18^, which were applied in refractive index-based concentration calculations to determine the mass contribution of each component. Mw was calculated from light scattering intensity measured at three detection angles (45°, 90°, and 135°) using Rayleigh-Gans-Debye (RGD) scattering theory. Smaller HDL particles (∼3.5 nm–5 nm Rh) exhibit isoscattering behavior (<10 nm Rh), meaning they scatter light nearly isotropically with minimal angular dependence, allowing Mw determination from lower-angle scattering data. Medium-sized HDL particles (∼5 nm–6 nm Rh) remain nearly isoscattering but may exhibit slight angular dependence. Larger HDL particles (∼6 nm–7.5 nm Rh) are closer to the isoscattering threshold and may begin to show more noticeable angular dependence, though they still behave approximately isotropically. LDL particles (∼9 nm–12.5 nm Rh) approach or exceed the isoscattering threshold (∼10 nm Rh); their larger size results in greater angular dependence, requiring multi-angle measurements to improve Mw accuracy.

The Rh was determined using DLS with a 658 nm laser. Rh was calculated from the diffusion coefficient using the Stokes-Einstein equation, assuming a spherical particle structure. Further details on UC-SEC and detector settings are available in Table 1.

**Table 1.** Key resource table for lipoprotein and EV isolation method using UC-SEC-MALS-DLS-UV

| REAGENT or RESOURCE | SOURCE | IDENTIFIER |
| --- | --- | --- |
| Biological samples |  |  |
| Human plasma pool<br>Male Pool<br>Pregnant Female Pool<br>Quality control | University of California, San Diego (UCSD)<br>University of California, Davis (UCD) |  |
| Chemicals, peptides, and recombinant proteins |  |  |
| Phosphate Buffered Saline 10X Molecular Biology Grade | Corning | MT-46013CM |
| Potassium Bromide (KBr) | Sigma Aldrich | CAT #243418 |
| Software and algorithms |  |  |
| ASTRA 8.1.2 | Waters-Wyatt Technology | https://wyatttechnology.zendesk.com/hc/en-us/articles/360052705793-ASTRA-8 |
| HPLC Connect 2.0.1.35 | Waters-Wyatt Technology | https://wyatttechnology.zendesk.com/hc/en-us/articles/360052705833-HPLC-CONNECT-2 |
| Other |  |  |
| 4.7 mL, OptiSeal Polypropylene Tube | Beckman Coulter | CAT #361621 |
| Amicon Ultra-4 Ultracell-50 4ml Tube | EMD Millipore | UFC805096 |
| Optima Max-TL Ultracentrifuge | Beckman Coulter | CAT #A95761 |
| TLA-110 Fixed-Angle rotor (k-factor 13) | Beckman Coulter | CAT #366735 |
| Agilent 1260 Infinity II | Agilent Technologies | https://www.agilent.com/en/product/liquid-chromatography/hplc-systems/analytical-hplc-systems/1260-infinity-ii-lc-system |
| miniDAWN | Waters-Wyatt Technology | https://www.wyatt.com/products/instruments/minidawn-multi-angle-light-scattering-detector-sec-mals.html |
| WyattQELS | Waters-Wyatt Technology | https://store.wyatt.com/shop/udawn/udawn3/wiq/ |
| Optilab | Waters-Wyatt Technology | https://www.wyatt.com/products/instruments/optilab-refractive-index-detector.html#optilab-1 |

#### Chromatogram Analysis and Quality Control

Analysis of chromatograms was performed to evaluate replicability of QC samples by assessing variability in the volume of elution of UV peaks (min) and UV and LS peak heights (1/cm and AU). To evaluate differences in particle abundances between fractions, the total area of the peak of the chromatogram was calculated by multiplying the signal from dRI in refractive index unit (RIU) with the elution time (min). The yield of particle abundances was also adjusted to the starting plasma sample to allow comparison of yield from the same starting plasma. Fractions obtained from non-dense LPP (1.21 g/mL) separation was compared against fractions from dense LPP (1.25 g/mL) separation to obtain a yield ratio. This method modification enables further separation of 6 total HDL subfractions that are differentiated by size and density from previously published methods^12^. Each LPP fraction was dialyzed then aliquoted. Identical aliquots were used for further analysis.

### 3. Western Blot

Samples were lysed in 1x radioimmunoprecipitation assay buffer (RIPA, Cell Signaling Technology, Cat. #9806) for 30 minutes at room temperature. Lysates were heated at 95°C for 5 minutes together with Laemmli sample buffer (Bio-Rad, Cat. #1610747, Non-reducing condition, all markers.

Lysates were resolved using a 4% to 15% Criterion TGX Stain-Free Precast gel (Bio-Rad, Cat.# 5678084), with Spectra Multicolor Broad Range protein ladder (Thermo Scientific, Cat.# 26634), then transferred onto a PVDF membrane (Invitrogen, Cat.# IB24001) using iBlot 2 semi-dry transfer system (Invitrogen) with p0 protocol. Blots were probed using primary antibodies in PBST (PBS with 0.05% Tween-20 (BioXtra, Cat. #P7949) and 5% Blotting Grade Blocker (Bio-Rad, Cat. #1706404) (PBST-Milk) overnight at 4°C, then washed 4 times in PBST-Milk, and incubated with corresponding primary antibody for 1 hour at room temperature, washed 2 times in PBST-Milk, and 2 times in PBST. SuperSignal West Pico PLUS Chemiluminescent Substrate (Thermo Scientific, Cat. # 34580) was used for detection, and blots were visualized with an iBright 1500FL Imager (Thermo Fisher, Waltham, MA). Further method details are available in Table 2.

**Table 2.** List of antibodies used for western blot analysis, including their manufacturer, catalog number, and dilution factor at which the products were used.

| REAGENT or RESOURCE | SOURCE | IDENTIFIER | DILUTION |
| --- | --- | --- | --- |
| Antibodies |  |  |  |
| mouse anti-human Serum Albumin | Abcam, MA, USA (now discontinued)/ | ab10241 | 1:1000 |
| mouse anti-human Serum Albumin | Invitrogen | MA1-19174 | 1:1000 |
| goat anti-human Apo A1 | Academy Bio-Medical, TX, USA | 11A-G2b | 1:1000 |
| goat anti-human Apo B-100 | Academy Bio-Medical, TX, USA | 20A-G1b | 1:1000 |
| goat anti-human Apo E | Academy Bio-Medical, TX, USA | 50A-G1b | 1:1000 |
| m-IgGk BP-HRP | Santa Cruz Biotechnology, CA, USA | sc-516102 | 1:10000 |
| mouse anti-goat IgG-HRP | Santa Cruz Biotechnology, CA, USA | sc-2354 | 1:10000 |

### 4. Non-Denaturing Gel Electrophoresis

Sample was prepared at 0.4 mg/mL protein concentration and mixed with diluted Native Sample Buffer (Bio-rad, U.S.A., Cat. #1610738) at 1:2 (v:v) ratio. The mixed samples were loaded into individual wells on a gradient precast protein gel (Bio-rad, U.S.A., Cat. #4561091) at a volume of 25 𝜇L per well. A protein molecular weight marker (Invitrogen, U.S.A., Cat. #LC5688) was loaded on the same precast gel to indicate the Mw of sample bands. The gel was then run with tris-glycine buffer (Genesee Scientific, Cat #: 18-238B) at 25 mA current. Bands were visualized using a commercial silver stain kit (Thermo Scientific, Cat. #24612) by following the manufacturer’s instructions. Further method details are available in Table 3.

**Table 3.** List of reagents and commercial kits in non-denaturing gel electrophoresis and silver stain analysis, including their manufacturer and catalog number.

| REAGENT or RESOURCE | SOURCE | IDENTIFIER |
| --- | --- | --- |
| Critical commercial assays |  |  |
| Pierce Silver Stain Kit | Thermo Scientific, USA | 24612 |
| Chemicals, peptides, and recombinant proteins |  |  |
| Native Sample Buffer | Bio-Rad, USA. | 1610738 |
| 4–20% Mini-PROTEAN® TGX™ Precast Protein Gels | Bio-Rad, USA. | 4561091 |
| Tris-Glycine, 10X | Genesee Scientific, USA. | 18-238B |
| HiMark™ Unstained Protein Standard | Invitrogen, USA | LC5688 |

### 5. Electron Microscopy Imaging and Size Analysis

Particles in each isolated fraction were viewed using negative-stain transmission electron microscopy (NS-TEM) using a previously described protocol^12^. Briefly, samples from each fraction were adjusted for dilution before being loaded to carbon-coated grids (TedPella Inc., CA, USA). After 5 minutes, excess sample was removed and stained with uranyl acetate solution, then blotted and left to dry at room temperature. Samples were then transferred to a specimen holder, then inserted into the TEM (JEOL USA 1230 Transmission Electron Microscope, JEOL, USA Inc., MA, USA). For each sample, five images were taken from different grid regions. The size of particles found in the micrographs was analyzed using a previously described method^12^.

### 6. NMR LipoProfile

LPP size distribution and subclass concentrations were also measured by NMR spectroscopy at Labcorp (USA), and the data were processed using the optimized version LipoProfile (LP4). NMR sizing characterization of fractions from SEC has been demonstrated to be comparable to gel filtration sizing^19^.

### 7. RNA Sequencing and Data Analysis

RNAs were extracted by Trizol LS (Thermo Fisher, USA) and purified by miRNeasy Mini Kit solutions (Qiagen, Netherlands) and Zymo-Spin I Columns (Zymo Research, USA) according to the manufacturer’s instructions. 8 µL of RNA were used for small RNA library construction by BioLiqX Small RNA-seq kit (Heidelberg Biolabs GmbH, Germany) and purified twice with AMPure XP reagent (Beckman Coulter, USA). The yield and size distribution of the small RNA libraries were assessed using the Fragment Bioanalyzer™ system with DNA 1000 chip (Agilent, USA). Multiplexed libraries were equally pooled to 1 nM and sequenced on the NextSeq 500 High 75 (Illumina, USA). Obtained FASTQ files were assessed for sequence read quality. Identified contamination from vector or adapter sequences and low-quality bases were removed as part of the trimming process. Further method details are available on Table 4 and Figure S1. PolyA enriched tails and Illumina adapter sequences were removed by cutadapt (version 3.4) as suggested by the laboratory manual of the BioLiqX small RNA-seq kit (Heidelberg Biolabs GmbH). All sequences shorter than 15 nucelotide (nt) were discarded. The resulting reads were aligned to manually curated hg38 reference transcriptome in a sequential manner using bowtie (version 1.2.2) enabling either 0 or 1 mismatch tolerance. Briefly, all reads were first mapped to human mitochondrial chromosome and discarded. The unaligned reads were next mapped to various small noncoding RNA biotypes with low sequence complexity (including rRNA, tRNA, RN7S, snRNA, sno/scaRNA, vault RNA, and RNY). The remaining reads were aligned sequentially to human mature miRNA, pre-miRNA, protein-coding mRNA transcripts (mRNA), as well as long noncoding RNAs. Finally, any remaining reads were mapped to the human reference genome to capture sequences originating from intronic and intergenic regions exclusively. All non-hg38 reads were subsequently extracted and mapped to small (16S/18S, SSU) and large subunit (23S/28S, LSU) ribosomal RNA (rRNA) references across all three domains of life (Bacteria, Archaea, and Eukarya) from the SILVA database (release 138.2, https://www.arb-silva.de/). Raw transcript counts were obtained using the eXpress package (version 1.5.1), converted to log-counts-per-million (lcpm), and visualized through multidimensional scaling (MDS) plots using the glimma package from R/Bioconductor. Differentially abundant LSU and SSU rRNA reads across analyzed fractions were visualized using the ggplot2 R package. The tables with mapping statistics to different RNA biotypes as well as raw count tables for all samples are available in Supplementary Materials.

**Table 4.**
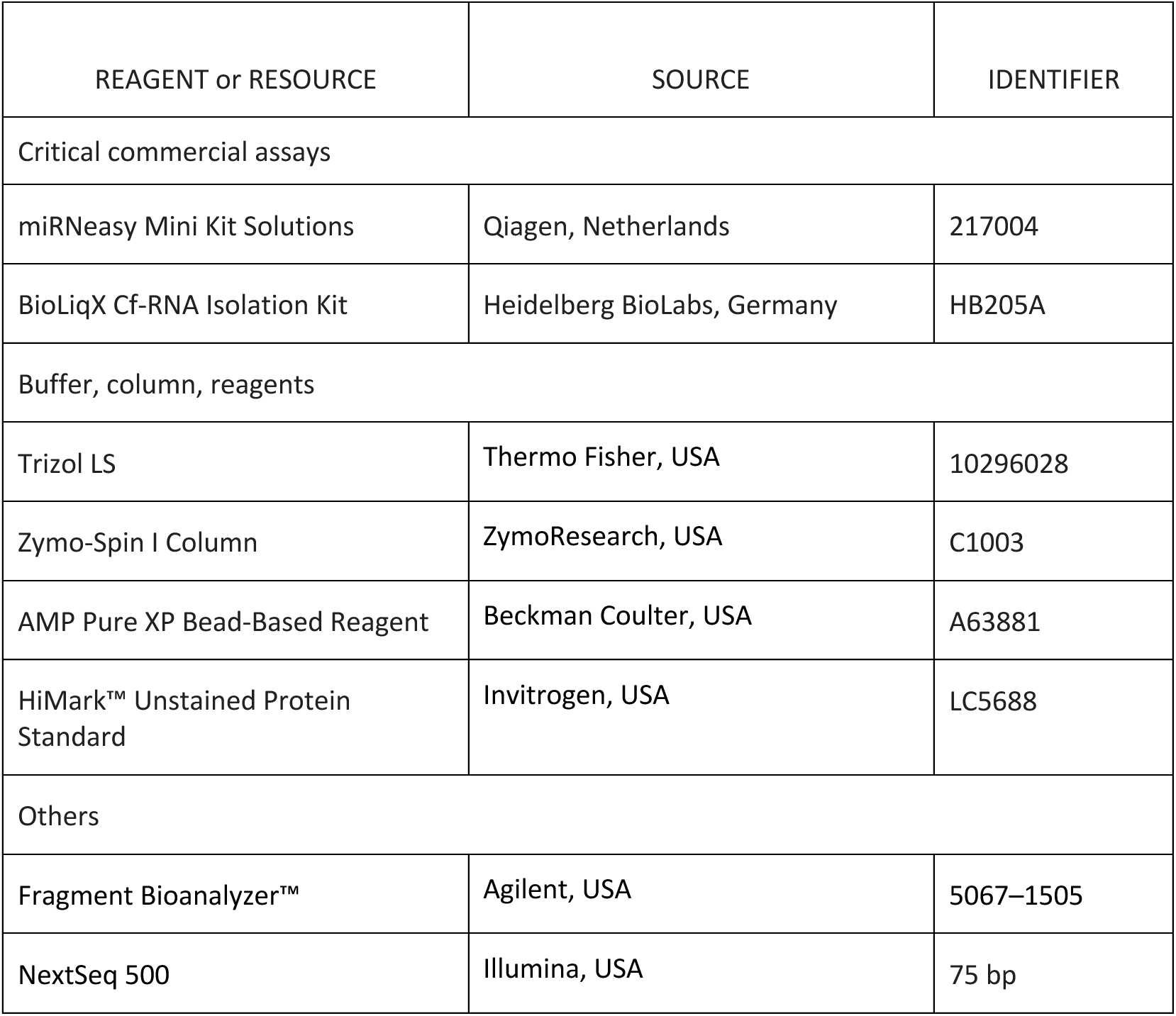
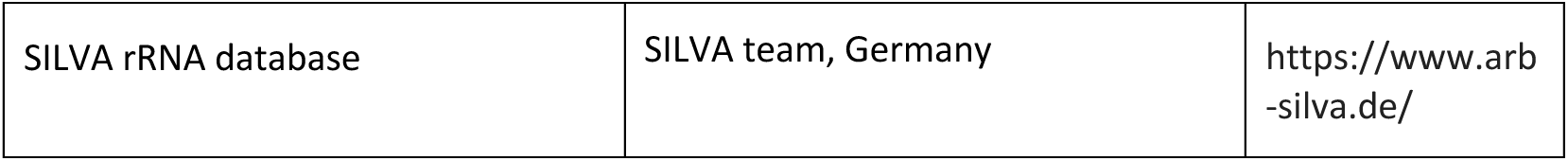
List of reagents and commercial kits in RNA sequencing analysis, including their manufacturer and catalog number.

| REAGENT or RESOURCE | SOURCE | IDENTIFIER |
| --- | --- | --- |
| Critical commercial assays |  |  |
| miRNeasy Mini Kit Solutions | Qiagen, Netherlands | 217004 |
| BioLiqX Cf-RNA Isolation Kit | Heidelberg BioLabs, Germany | HB205A |
| Buffer, column, reagents |  |  |
| Trizol LS | Thermo Fisher, USA | 10296028 |
| Zymo-Spin I Column | ZymoResearch, USA | C1003 |
| AMP Pure XP Bead-Based Reagent | Beckman Coulter, USA | A63881 |
| HiMark™ Unstained Protein Standard | Invitrogen, USA | LC5688 |
| Others |  |  |
| Fragment Bioanalyzer™ | Agilent, USA | 5067–1505 |
| NextSeq 500 | Illumina, USA | 75 bp |
| SILVA rRNA database | SILVA team, Germany | https://www.arb-silva.de/ |

### 8. Lipidomics

Lipidomic analysis was performed at the UC Davis West Coast Metabolomics Center using untargeted liquid chromatography followed by mass spectrometry (LC-MS) with 23 internal standards [carnitine (CAR) (18:0), cholesterol ester (CE) (20:3), Cer(d18:1/24:1), diglyceride (DG) (17:0/20:3), lysophosphatidyl choline (LPC) (17:0), lysophosphatidyl ethanolamine (LPE) (17:0), lysophosphatidyl glycerol (LPG) (17:0), lysophosphatidyl inositol (LPI) (17:0), LPS (17:0), phosphatidyl choline (PC) (17:0/18:1), phosphatidyl enthanolamine (PE) (17:0/22:4), phosphatidyl glycerol (PG) (17:0/18:1), phosphatidyl inositol (PI) (17:0/16:1), PS (17:0/16:1), sphingomyelin (SM) (d18:1/16:1), Sphingosine (d17:1), triglyceride (TG) (16:0/17:1/16:0), ceramide (Cer) (d18:1/16:1), fatty acid (FA) (20:4), PE (17:0/18:1), PG (17:0/20:3), PI (17:0/18:1), SM (d18:1/16:1)] using a previously published protocol^20^. Briefly, samples were extracted using the Matyash extraction procedure which includes MTBE, MeOH, and H2O^21^. The organic (upper) phase was dried down and resuspended with 110µL of a solution of 9:1 methanol: toluene and 50 ng/mL 12-[[(cyclohexylamino)carbonyl]amino]-dodecanoic acid (CUDA). This solution was shaken for 20 seconds, sonicated for 5 minutes at room temperature, and then centrifuged for 2 minutes at 16100 relative centrifugal force. The samples were then aliquoted into three parts: 33µL were aliquoted into a vial with a 50µL glass insert for positive and negative mode lipidomics, and the remainder was aliquoted into an eppendorf tube to be used as a pool. The samples were then loaded onto an Agilent 1290 Infinity LC stack then followed by Agilent 6546 QTOF mass spectrometer (Agilent Technologies Inc., CA, USA). Both positive and negative modes were run on an Agilent 6546 with a scan range of m/z 120-1200 Da with an acquisition speed of 2 spectra/s. The gradient used was 0 min 15% (B), 0.75 min 30% (B), 0.98 min 48% (B), 4.00 min 82% (B), 4.13-4.50 min 99% (B), 4.58-5.50 min 15% (B) with a flow rate of 0.8 mL/min. The mass resolution for the Agilent 6546 was 10,000 for electrospray ionization (ESI) (+) and 30,000 for ESI (-). Lipidomics analysis using this method on isolated HDL particles has also been performed previously^22^. The concentration of lipid species was quantified by comparing the peak heights of each lipid species to those of the internal standard for that lipid class and ESI mode. The concentration was presented in ng/mL. The total amount of lipid species within each class was summed to calculate the fractional composition of each lipid class.

### 9. Proteomic Analysis Using LC-MS/MS

The method for untargeted proteomics analysis was adapted from a previous publication^23^. Briefly, the peptide samples were reconstituted with nanopure water and directly characterized using UltiMate™ WPS-3000RS nanoLC 980 system coupled to the Nanospray Flex ion source of an Orbitrap Fusion Lumos Tribrid Mass Spectrometer system (Thermo Fisher Scientific, MA, United States). The analytes were separated on an Acclaim™ PepMap™ 100 C18 LC Column (3 μm, 0.075 mm × 150 mm, ThermoFisher Scientific, USA). A binary gradient was applied using 0.1% (v/v) formic acid in (A) water and (B) 80% acetonitrile: 0–5 min, 4–4% (B); 5–133 min, 4–32% (B); 133–152 min, 32%–48% (B); 152–155 min, 48–100% (B);155–170 min, 100–100% (B); 170–171 min, 100–4% (B); 171–180 min, 4–4% (B). The instrument was run in data-dependent mode with 1.8 kV spray voltage, 275°C ion transfer capillary temperature, and the acquisition was performed with the full MS scanned from 700 to 2000 m/z in positive ionization mode. Stepped higher-energy C-trap dissociation (HCD) at 30 ± 10% was applied to obtain tandem MS/MS spectra with m/z values starting from 120.

Peptide fragmentation spectra were annotated using Byonic software (Protein Metrics, CA, United States) against the reviewed human proteome sequences from Uniprot. Common modifications, including cysteine carbamidomethyl, methionine oxidation, asparagine deamidation and glutamine deamidation were assigned. The most abundant proteins, along with those closely associated with HDL function, were selected as target proteins. The fractional composition of these target proteins was then calculated within the different LPP fractions. Percent abundance was analyzed using intensities of peptide ion counts over total ion counts to obtain a relative protein quantitation.

## Method Limitations

There are multiple approaches for isolating HDL particles, each with inherent limitations. Our method builds upon the UC-SEC workflow previously described (Zheng et al., 2021), where concerns such as protein loss and structural perturbation are particularly relevant for ultracentrifugation-based protocols. These effects are more pronounced under higher g-forces and prolonged spin times, and some disruption may also occur due to ionic strength of salt buffers used during density separations (Davidson et al., 2022). Light scattering techniques such as MALS and DLS offer non-destructive means of characterizing particle size and composition (Matson, 2023), but depend on accurate input values—especially the refractive index increment (dn/dc), which remains poorly defined for lipids in aqueous buffers like PBS. MALS-derived Rg is limited for smaller spherical particles like HDL (<10 nm), necessitating DLS pairing to determine Rh. However, DLS assumes uniform solvent interactions across particle subtypes, which may not hold true for all lipoproteins. SEC-based separations also require careful attention to column cleanliness and sample-resin interactions, as poor maintenance can compromise resolution. Finally, increased tubing volume between separation and collection points can introduce peak broadening. Nonetheless, our particle validation results confirm accurate **(not sure if there is a better word than accurate)** retention of expected lipoprotein classes, underscoring the robustness of the optimized method.

Davidson, 2022: https://doi.org/10.1016/j.bbalip.2021.159072

Matson, 2023: DOI: 10.1039/D3PY01181J

## Results

Figure 2 shows overlaid chromatograms of the UV and LS signals of the QC plasma pool of the injected solution after the second UC step, which included particles between 1.006–1.21 g/mL across 5 isolation runs. Repeatability of sample processing can be found on Table 5, Figure 3. Additionally, particle size distribution, in Rh (Figure 4) and Mw, were also analyzed using MALS and DLS (Table 6). Injections of samples from additional UC times at the same densities were performed to evaluate whether the current UC times and rotation speeds were effective in separating particles through the UC steps (Supplemental figure S1).

**Figure 2.**
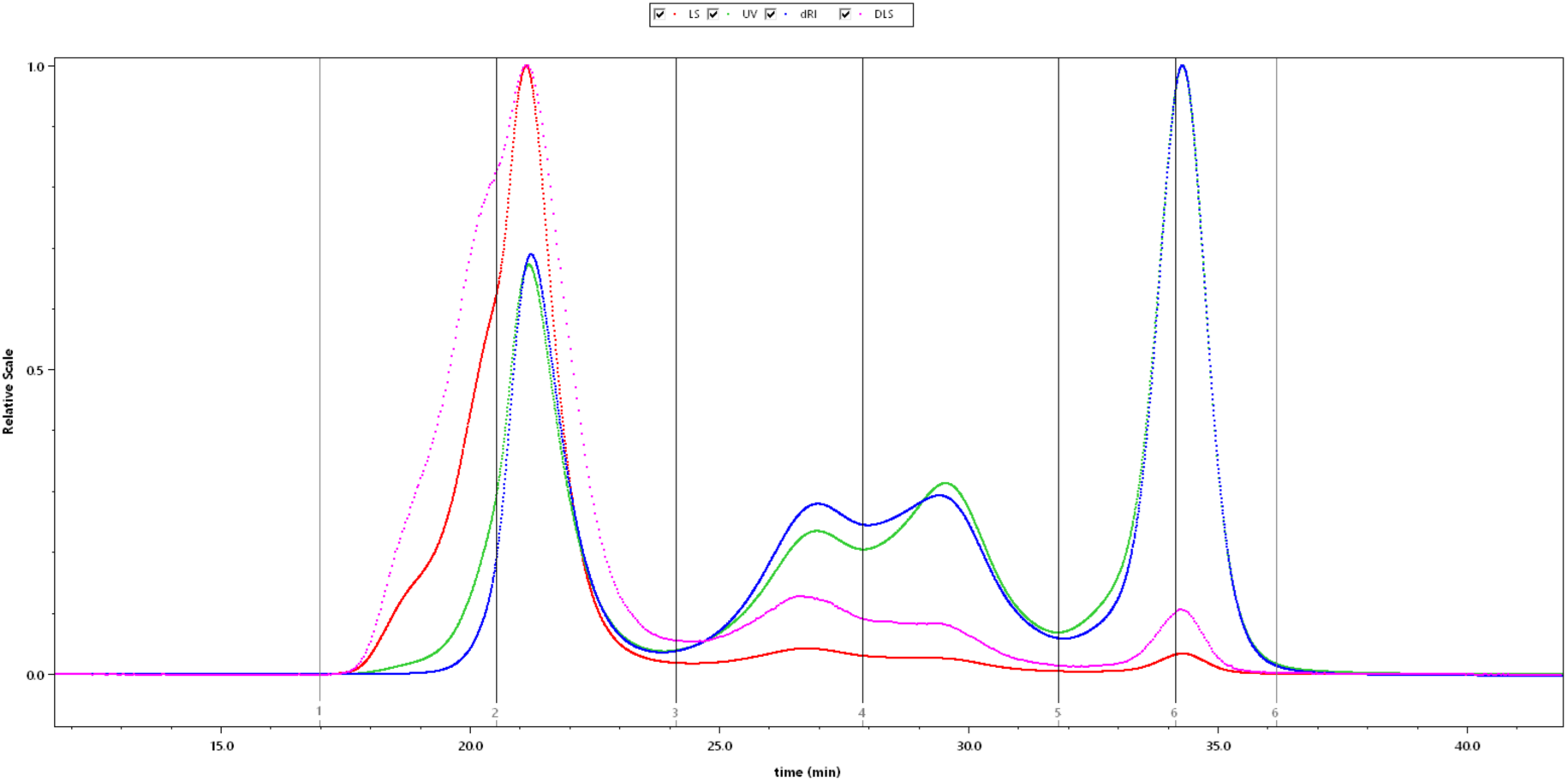
Overlaid chromatograms of lipoprotein particles and albumin between 1.006-1.21 g/mL density. Four detectors - multi angle light scattering (MALS; red), ultraviolet (UV; green), differential refractive index (dRI; blue), and dynamic light scattering (DLS; pink) - were used to analyze particles eluting from the size exclusion chromatography column. The X-axis shows the elution time and the Y-axis shows the signal quantity in relative scale. Each chromatogram is scaled to its highest peak (valued at 1.0 relative scale). From left to right, fractions include intermediate density lipoprotein (IDL, eluting between lines labeled 1-2), low density lipoprotein (LDL, eluting between lines labeled 2-3), high density lipoprotein large (HDL-L, eluting between lines labeled 3-4), high density lipoprotein medium (HDL-M, eluting between lines labeled 4-5), high density lipoprotein small (HDL-S, eluting between lines labeled 5-6), and albumin (Alb, eluting between lines labeled 6-6).

**Figure 3.**
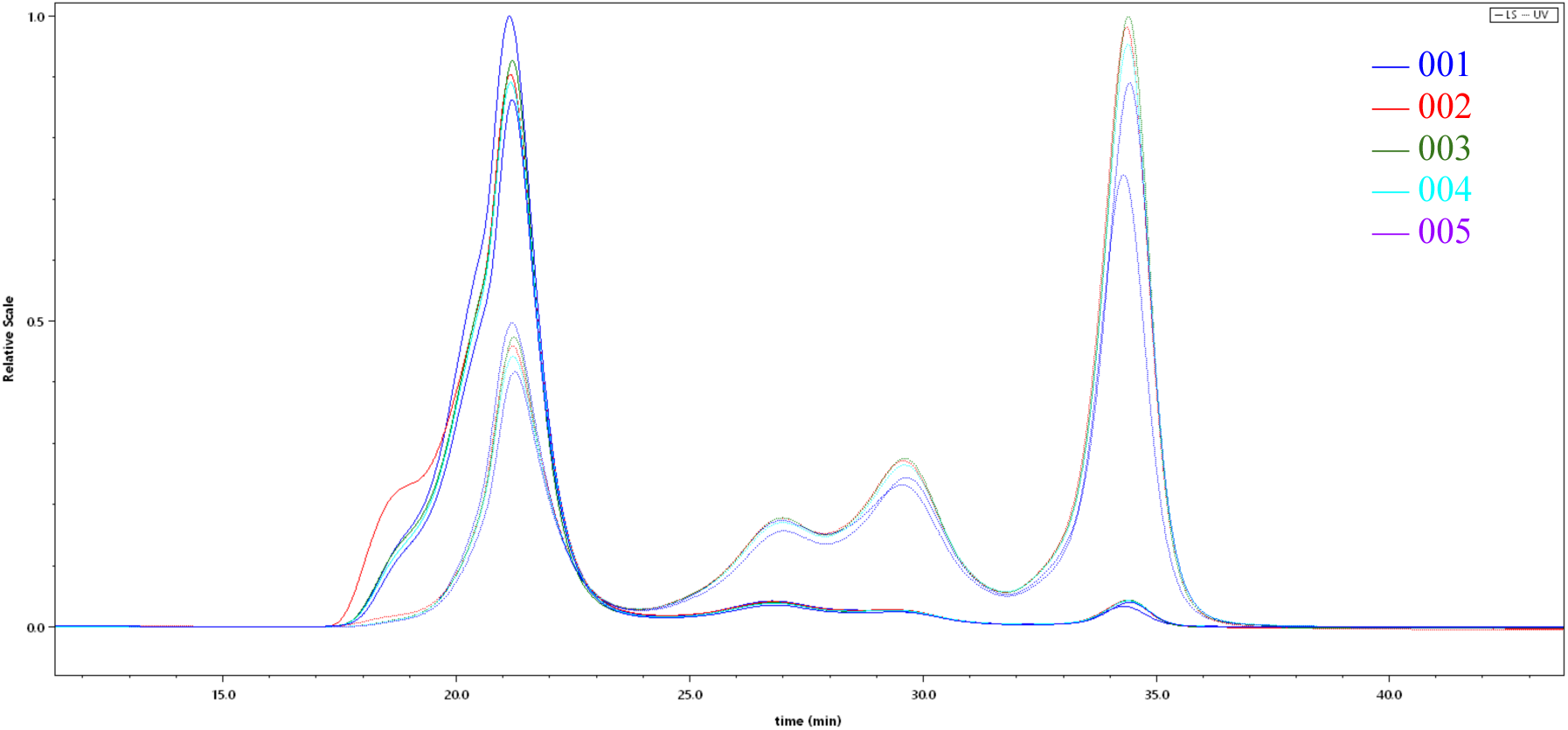
Chromatogram of overlaid lipoprotein particles (density 1.006-1.21 g/mL) from five plasma pool sample isolations and injections (001, 002, 003, 004, 005) demonstrating technical reproducibility. Ultraviolet (UV) signal at 280 nm is represented by solid lines, while light scattering (LS) signal is shown by dotted lines. The chromatogram is plotted with elution time (minutes) on the X-axis and relative scale on the Y-axis, where each chromatogram is normalized to its respective highest peak (1 relative scale).

**Figure 4.**
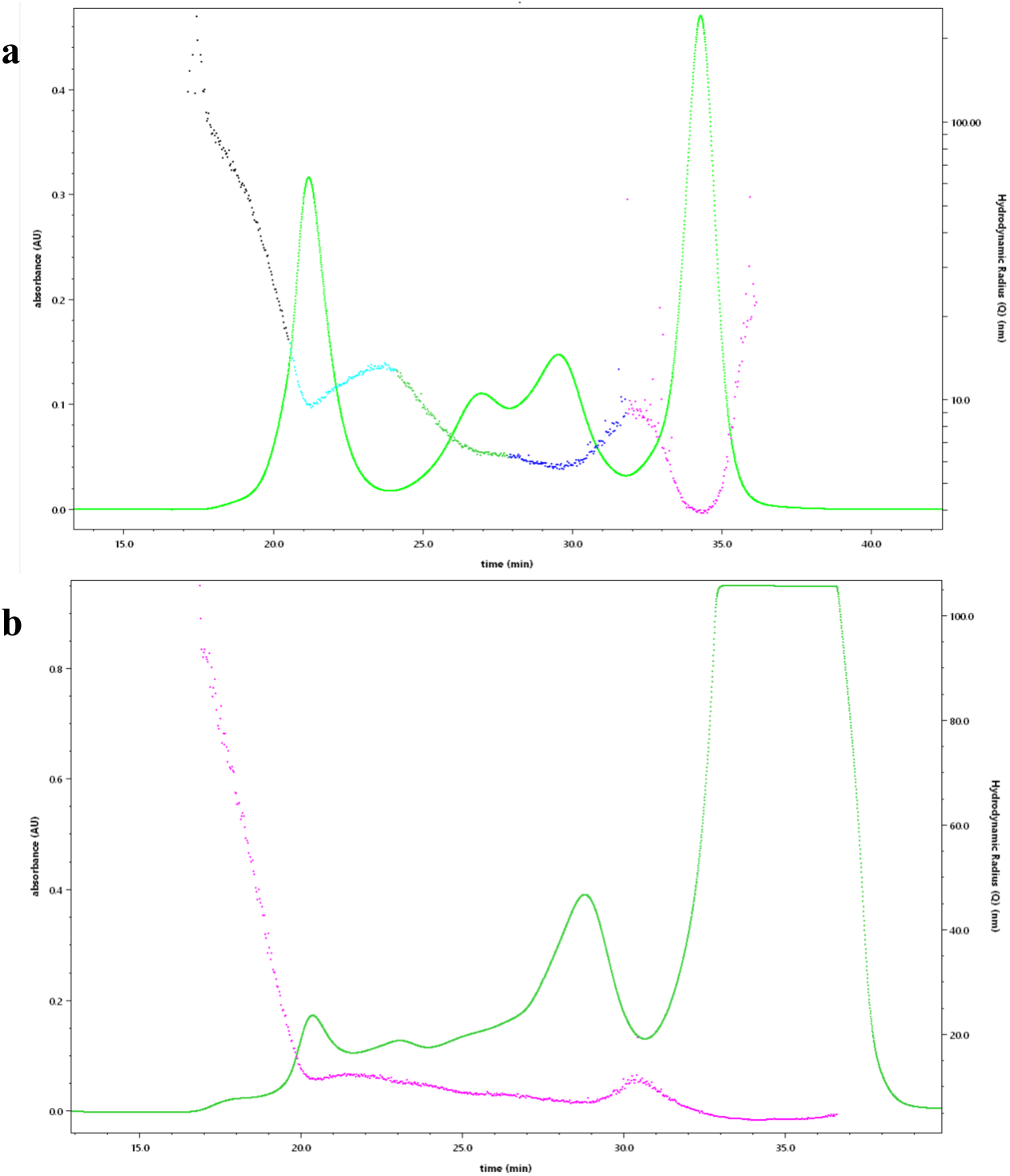
Distribution of hydrodynamic radii (Rh) of particles measured by dynamic light scattering (DLS; dotted lines) along with the ultraviolet chromatogram (UV) at 280nm (solid lime green) of a) particles between the 1.006-1.21 g/mL density range, which includes, intermediate density lipoprotein (IDL, black dots), low density lipoprotein (LDL, cyan dots), high density lipoprotein large (HDL-L, green dots), high density lipoprotein medium (HDL-M, blue dots), and high density lipoprotein small (HDL-S) and albumin (pink dots) and b) all fractions across the 1.21-1.25 g/mL density range. X axis represent time of elution (min) and Y axis represent UV absorbance (AU, left) and hydrodynamic radii (nm, right).

**Table 5.** Variability of light scattering (LS) peak heights, ultraviolet (UV) peak height, and UV elution time (min) between sample isolation run 001 to 005 of 1.006-1.21 g/mL LPP. The mean ± standard deviation and % coefficient variation of UV peak elution for peak 1, 2, 3, and 4 are 21.2 min ± 0.0277 (0.131%), 26.8 min ± 0.01818 (0.0679%), 29.6 min ± 0.04549 (0.117%), and 34.4 min ± 0.04908 (0.143%).

|  | Peak 1 |  |  | Peak 2 |  |  | Peak 3 |  |  | Peak 4 |  |  |
| --- | --- | --- | --- | --- | --- | --- | --- | --- | --- | --- | --- | --- |
| Sample | LS peak height (1/cm) | UV peak height (AU) | UV Elution time (min) | LS peak height (1/cm) | UV peak height (AU) | UV Elution time (min) | LS peak height (1/cm) | UV peak height (AU) | UV Elution time (min) | LS peak height (1/cm) | UV peak height (AU) | UV Elution time (min) |
| 1 | 11.1000 | 0.3170 | 21.1000 | 0.4670 | 0.1110 | 26.8000 | 0.2980 | 0.1470 | 29.5000 | 0.3720 | 0.4700 | 34.3000 |
| 2 | 10.0000 | 0.2920 | 21.2000 | 0.4520 | 0.1130 | 26.8000 | 0.3080 | 0.1730 | 29.6000 | 0.4780 | 0.6240 | 34.4000 |
| 3 | 10.3000 | 0.3020 | 21.2000 | 0.4350 | 0.1130 | 26.8000 | 0.2960 | 0.1750 | 29.6000 | 0.4810 | 0.6360 | 34.4000 |
| 4 | 9.8700 | 0.2820 | 21.2000 | 0.4260 | 0.1090 | 26.8000 | 0.2940 | 0.1680 | 29.6000 | 0.4710 | 0.6060 | 34.4000 |
| 5 | 9.5600 | 0.2650 | 21.2000 | 0.3920 | 0.0997 | 26.8000 | 0.2700 | 0.1550 | 29.6000 | 0.4420 | 0.5660 | 34.4000 |
| Average | 10.2000 | 0.2910 | 21.2000 | 0.4350 | 0.1090 | 26.8000 | 0.2930 | 0.1640 | 29.6000 | 0.4490 | 0.5800 | 34.4000 |
| Coefficient of Variation (%) | 5.6600 | 6.6900 | 0.1310 | 6.5400 | 5.1100 | 0.0679 | 4.8200 | 7.3000 | 0.1170 | 10.1000 | 11.5000 | 0.1430 |
| Standard Deviation | 0.5750 | 0.0195 | 0.0277 | 0.0284 | 0.0056 | 0.0182 | 0.0142 | 0.0120 | 0.0347 | 0.0455 | 0.0669 | 0.0491 |

**Table 6.**
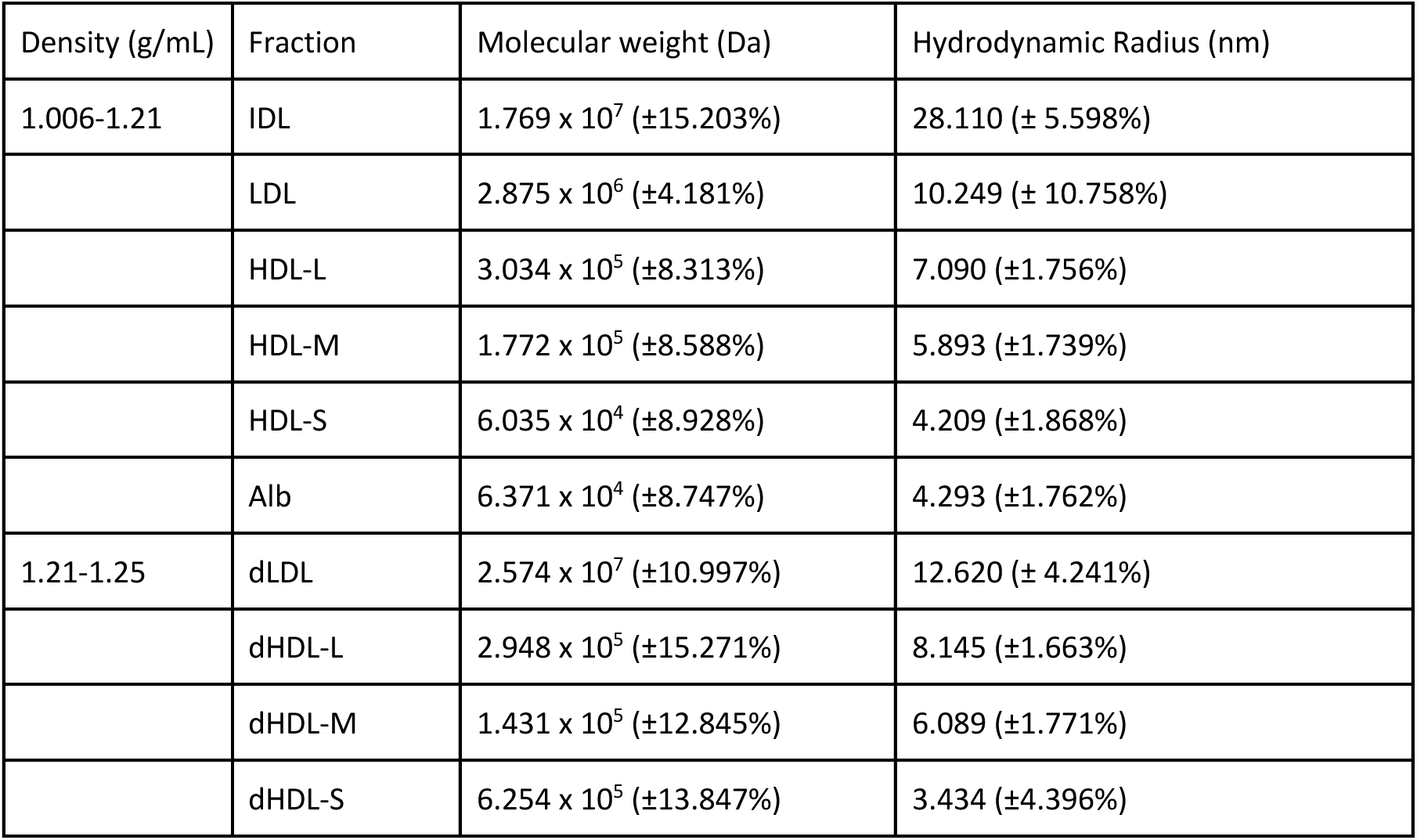
Molecular weights in kilodaltons (kDa) and hydrodynamic radii (nm) of LPP particles analyzed by Multi Angle Light Scattering (MALS) and dynamic light scattering (DLS) instruments. Fractions analyzed includes triglyceride rich lipoprotein (TRLP); dense plasma protein (DPP) as all particles denser than the density of 1.25 g/mL; intermediate density lipoprotein (IDL); low density lipoprotein (LDL); high density lipoprotein large (HDL-L); high density lipoprotein medium (HDL-M); high density lipoprotein small (HDL-S); albumin (Alb); dense LDL (dLDL); dense HDL-L (dHDL-L); dense HDL-M (dHDL-M); dense HDL-S (dHDL-S).

Both the measurements of Mw and Rh (Table 6) in the dense and non-dense LPP decreased over the elution times and across the peaks from left to right, except for measurements at the peak troughs where there is minimal signal and some noise from carryover of larger particles. These values include average Mw and Rh of 2.865 x 10^3^ kDa and 10.249nm Rh for LDL, 303.4 kDa and 7.090 nm Rh for HDL-L, 177.2 kDa and 5.893nm Rh for HDL-M, and 63.7 kDa and 4.209nm Rh for HDL-S. This demonstrates that the size of the particles is decreasing over the elution range of the column, as expected for SEC. In Table 5 we demonstrated the repeatability of the UC-SEC method by assessing the elution volume of peaks in the chromatogram from several sample processing runs and injections. Additionally, data from the dRI detector showed total yield (mass or RIU min) of particles obtained in each fraction and was used to compare differences in total mass of particles in the dense vs. non-dense LPP. Sample captured in the fraction HDL-L was about 4 times more concentrated than the fraction in dHDL-L. Sample captured in the fraction HDL-M was about 10,000 times more concentrated than the fraction in dHDL-M (Table 7).

**Table 7.**
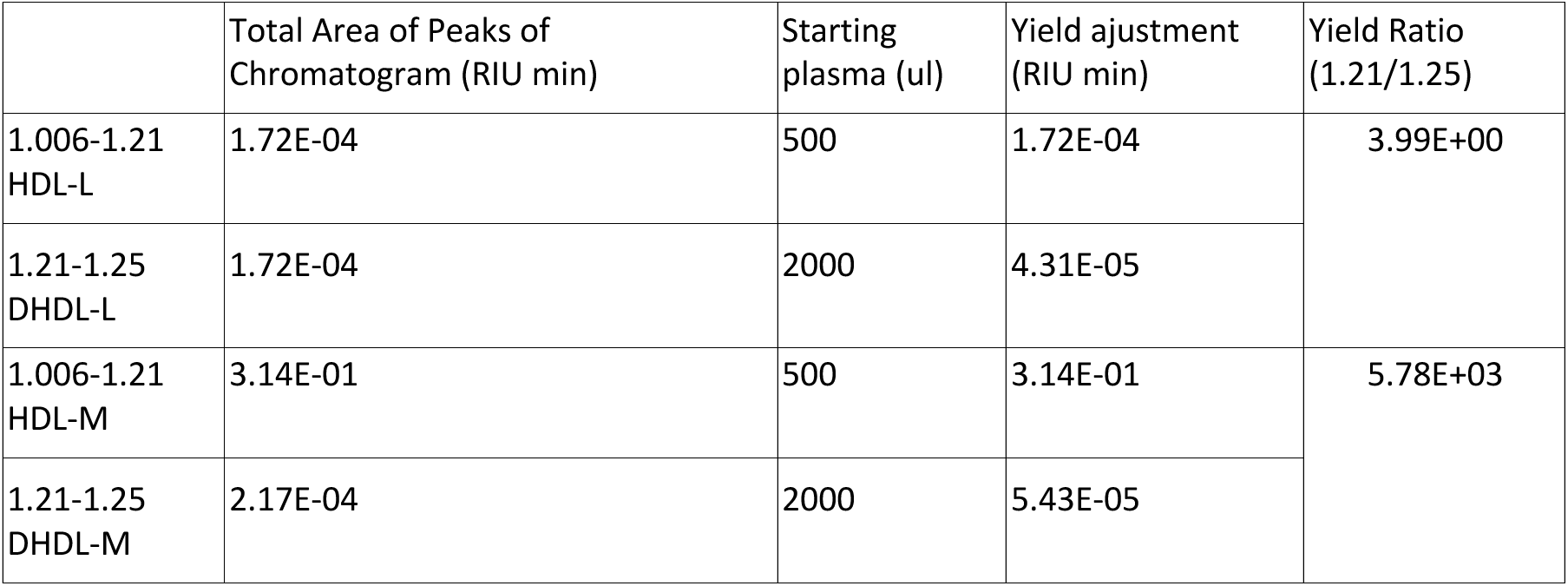
Yield differences between high density lipoprotein large (HDL-L) and high density lipoprotein medium (HDL-M) between 1.006-1.21 g/mL density and dense HDL-L (DHDL-L) and dense HDL-M (DHDL-M) between 1.21-1.25 g/mL as measured in, and compared by, refractive index unit (RIU) multiplied by elution time (min).

WB analysis of denatured LPP and EV samples from the UC-SEC isolation shows that Apolipoprotein B-100 (ApoB-100, 516 kDa, P04114) was most abundant in the TRLP, LDL, HDL-L, and dHDL-L fractions (Figure 5). ApoB-100 bands were also found to be distributed across the size spectrum in TRLP, LDL, and HDL-L fractions. Apolipoprotein A-1 (ApoA-1, 31 kDa, P02647) was most abundant in the HDL-L, HDL-M, HDL-S, dHDL-L, and dHDL-M fractions. Apolipoprotein E (ApoE, 36 kDa, P02649) was most abundant in LDL, HDL-L, HDL-M, dHDL-L, and dHDL-M. ApoE bands were also heterogeneously distributed across multiple size ranges including 40 kDa, 50 kDa, and 70 kDa in the LDL, HDL-L, HDL-M, dLDL, and dHDL-L fractions. Human serum albumin (Alb, 69,367 Da, P02768) was most abundant in the Alb, and dHDL-S fractions. Bands were heterogeneously distributed across 40-140 kDa in both the Albumin and dHDL-S fractions, and a band of around 100 kDa was present in the Alb fraction and all dense LPP fractions.

**Figure 5.**
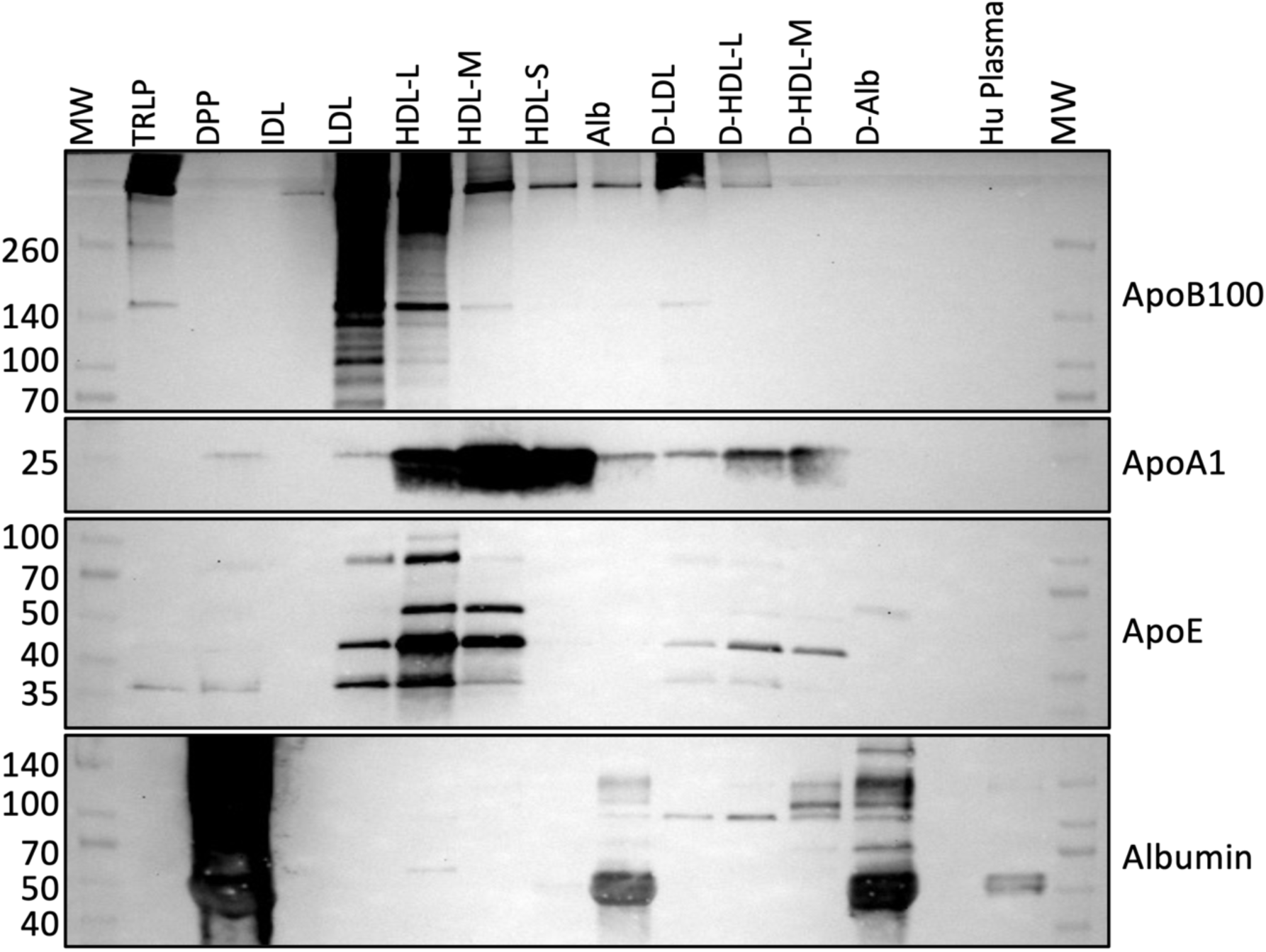
Western blot of proteins across the isolation fractions, which includes apolipoprotein B-100 (ApoB-100), apolipoprotein A-I (ApoA-I), apolipoprotein E (ApoE), and human serum albumin (Albumin). Fractions analyzed include triglyceride rich lipoprotein (TRLP); dense plasma protein (DPP) as all particles with a density >1.25 g/mL; intermediate density lipoprotein (IDL); low density lipoprotein (LDL); high density lipoprotein large (HDL-L); high density lipoprotein medium (HDL-M); high density lipoprotein small (HDL-S); albumin (Alb); dense LDL (dLDL); dense HDL-L (dHDL-L); dense HDL-M (dHDL-M); dense HDL-S (dHDL-S). Samples applied were normalized by volume. Lefthand markers are for molecular weight ladder, measured in kDa.

NDGE revealed the Mw distribution of particles across all LPP fractions (Figure 6). Most fractions showed particles within distinct Mw ranges, though only a few clear bands were visible. No bands appeared in the TRLP fraction within the Mw ladder. In the DPP fraction, distinct bands were seen at ∼500 kDa, 160 kDa, and at the bottom of the gel. Light bands in the IDL fraction were observed just below 160 kDa and at the bottom of the gel. A prominent band was noted in the LDL fraction above 500 kDa. In HDL-L, two regions of staining, at 160–290 kDa and above 500 kDa, were evident. The HDL-M fraction displayed a dense staining pattern from

**Figure 6.**
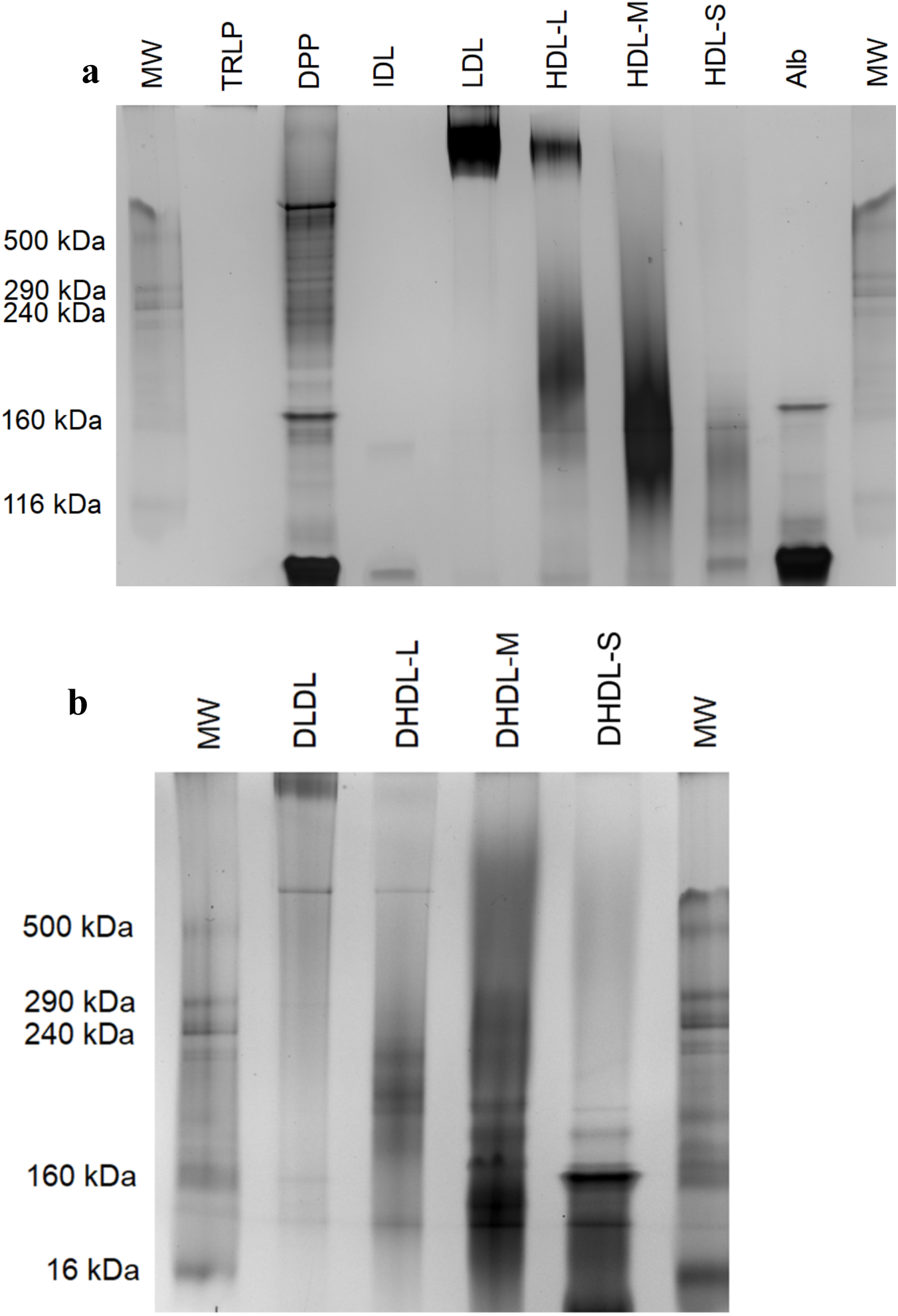
Silver-stained non-denaturing gel of samples across isolation fractions, which includes a) gel scan of fractions: triglyceride rich lipoprotein (TRLP); dense plasma protein (DPP) as all particles with a density >1.25 g/mL; intermediate density lipoprotein (IDL); low density lipoprotein (LDL); high density lipoprotein large (HDL-L); high density lipoprotein medium (HDL-M); high density lipoprotein small (HDL-S); albumin (Alb), b) gel scan of dense LDL (dLDL); dense HDL-L (dHDL-L); dense HDL-M (dHDL-M); dense HDL-S (dHDL-S). Molecular weight ladders are present in the furthest left and right of each gel. Samples applied were normalized by volume.

116–240 kDa, while faint bands were seen in HDL-S at 116–160 kDa and below 116 kDa. In the Alb fraction, two strong stains were observed at 160 kDa and below 116 kDa. Furthermore, Figure 5b show a single band above 500 kDa in the dLDL fraction, and a dense stain at the top of the gel. dHDL-L exhibited a band above 500 kDa and staining between 160–240 kDa. dHDL-M contained particles distributed across the Mw ladder, with the most intense staining between 16 kDa and just above 160 kDa. Finally, dHDL-S showed a strong band at 160 kDa and a distribution of smaller particles.

EM images show presence of round-lucent structures of particles (Figure 7) (previously described for LPP) with trace amounts of particles in the DPP and IDL fractions. Of those particles present in the DPP fraction, most particles were detected as being < 10nm. The largest particles are those in the TRLP fraction, followed by the LDL and dLDL fractions. Particles captured in the dLDL fraction were observed to be smaller and more monodispersed than particles in the LDL fraction. Particles captured in the HDL-L fraction were observed to be more monodispersed than particles in the dHDL-L fraction. Additionally, particles captured in the dHDL-S fraction more closely resemble particles captured in Alb. Previous EM imaging of the column void volume and the IDL fraction captured large, round, unilamelar, polydispersed particles with diameter spanning from 110 nm to 190 nm (Supplemental Figure S2).

**Figure 7.**
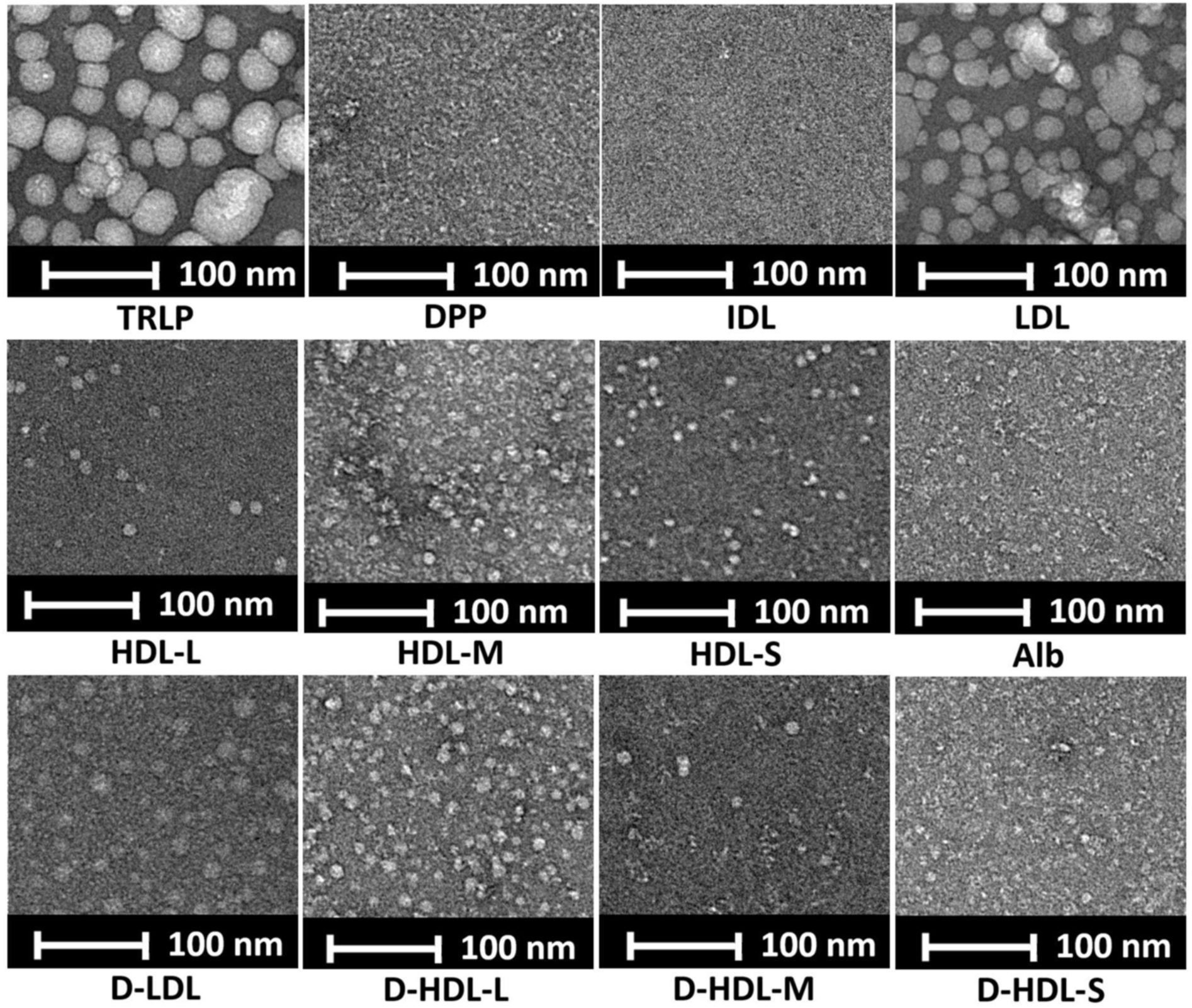
Representative negative-stain transmission electron microscopy (TEM) micrographs of samples across isolation fractions. Fractions analyzed include triglyceride rich lipoprotein (TRLP); dense plasma protein (DPP) as all particles with a density >1.25 g/mL; intermediate density lipoprotein (IDL); low density lipoprotein (LDL); high density lipoprotein large (HDL-L); high density lipoprotein medium (HDL-M); high density lipoprotein small (HDL-S); albumin (Alb); dense LDL (dLDL); dense HDL-L (dHDL-L); dense HDL-M (dHDL-M); dense HDL-S (dHDL-S). A 100nm bar is shown for each fraction.

NMR LipoProfile analysis showed enrichment of the target particles in each LPP fraction (Figure 8). TRLP were most abundant in the TRLP fraction, which is the fraction of particles with density < 1.006 g/mL from the first UC spin. No particles were detected in the IDL fraction by NMR LipoProfile. The LDL fraction contained the majority of detected LDL particles across the size range (large, medium, and small), and some TRLP particles. The HDL-L fraction mostly contained large HDL particles with some small HDL particles and small LDL particles. The HDL-M fraction contained mostly medium HDL particles followed by lower amounts of small and large HDL, with minimal presence of LDL. The HDL-S fraction contained mostly small HDL particles, followed by small amounts of medium and large HDL. Any detected LPP particles in the Alb fraction were mostly small HDL particles.

**Figure 8.**
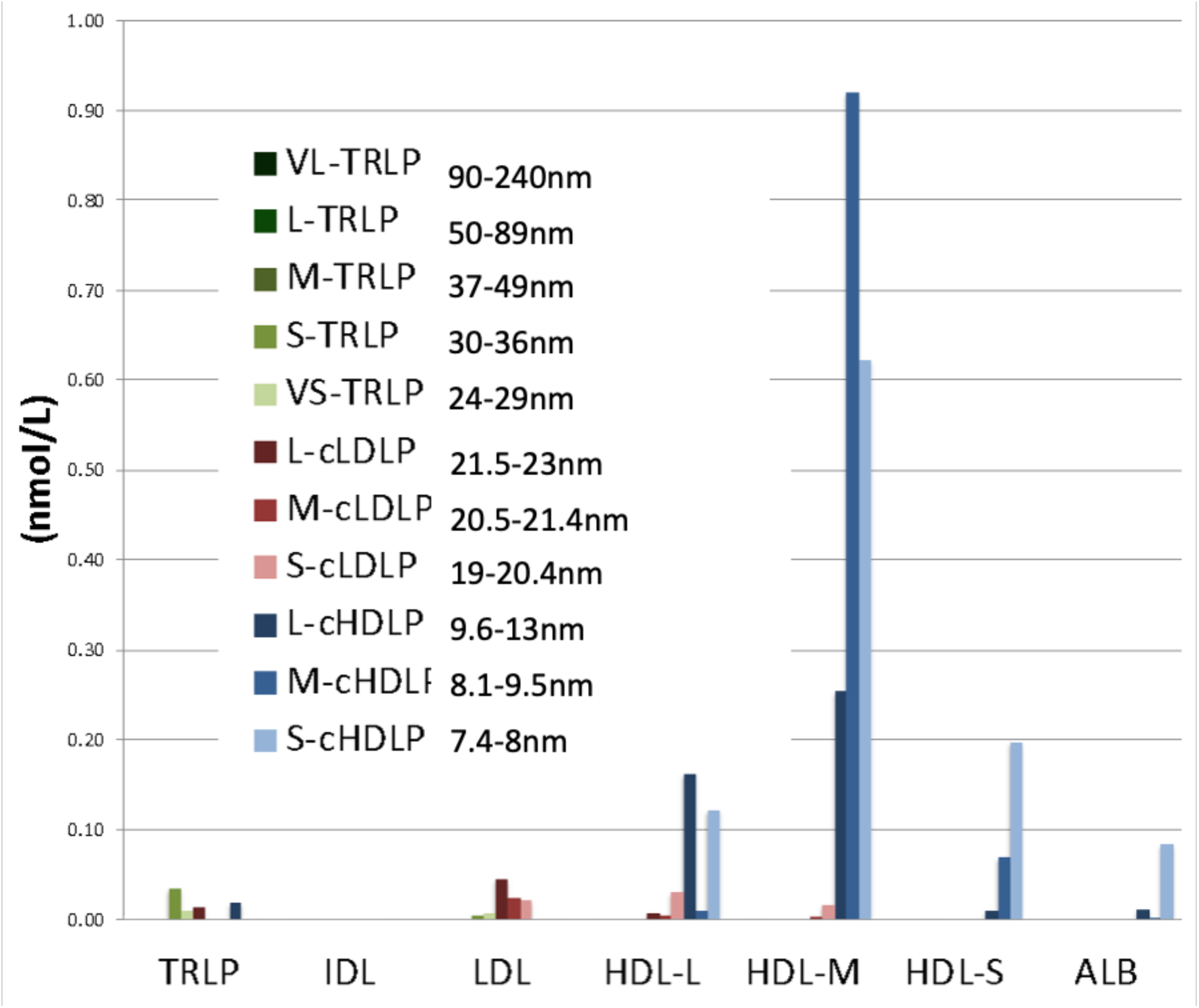
Nuclear magnetic resonance (NMR) LipoProfile data of lipoprotein distribution across isolation fractions. Green bars represent concentrations of triglyceride rich lipoprotein particles (TRLP), red bars represent concentrations of calibrated low density lipoprotein particles (cLDLP), and blue bars represent concentrations of calibrated high density lipoprotein particles (cHDLP). Darker colored bars represent very large (VL), and large particles (L) while lighter colored bars represent medium particles (M) and the lightest colored bars represent smaller particles (S) and very small (VS) of each lipoprotein category. Size ranges of large, medium and small can be found in supplemental table (Table S2).

Lipidomic analysis of isolated LPP fractions showed that PC was the predominant lipid component in both the LDL and HDL fractions (Figure 9). Free fatty acids (FFA) was the predominant lipid component in the IDL and DPP fractions, and TG was the predominant lipid component in the TRLP fraction. CE proportion decreased in particles through the elution time - CE proportion was highest in the LDL fraction, followed by HDL-L, then HDL-M, and HDL-S. We observed a similar trend in the CE proportion of dense particles. CE proportion was highest in dLDL, followed by dHDL-L, dHDL-M, and dHDL-S. SM was also present in both lower-density and dense particles, though at lower abundance than in the non-dense particles. The lipidome of classic density HDL was characteristic of what has been previously described, with 45-55 mol% phospholipid, 20-25 mol% CE, 5-10 mol% TG, and 2-10 mol% unesterified cholesterol. Conversely, the dense HDL particles were enriched in unesterified cholesterol (10-25 mol%) and depleted in CE (10-15 mol%) and TG (3-6 mol%), characteristic of particles with a smaller lipid core and a phospholipid monolayer enriched in lipid rafts. LPC made up the highest proportion in the DPP fraction, and less than 5 mol% of lipid content in the LPP fractions. Proteomic analysis (Figure 10) revealed that ApoB was the dominant protein in the TRLP and LDL fractions, while albumin was most abundant in the DPP, IDL, and Alb fractions.

**Figure 9.**
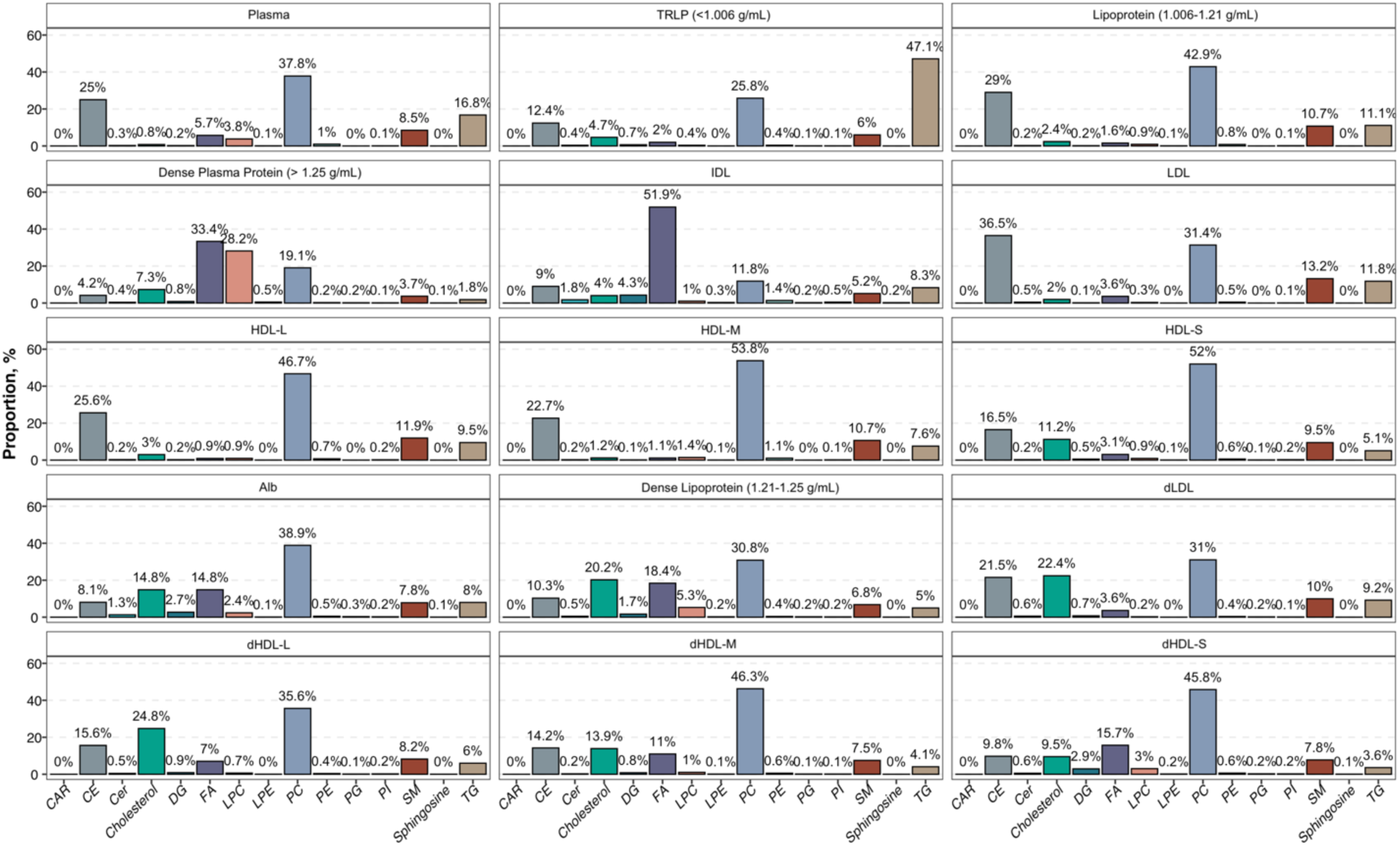
Lipidomic analysis showing relative proportions of lipid classes across isolated fractions in percentages (%). Fractions analyzed include Plasma, as whole plasma; triglyceride rich lipoprotein (TRLP); lipoprotein as all particles with a density of 1.006-1.21 g/mL; dense plasma protein (DPP) as all particles with a density > 1.25 g/mL; intermediate density lipoprotein (IDL); low density lipoprotein (LDL); high density lipoprotein large (HDL-L); high density lipoprotein medium (HDL-M); high density lipoprotein small (HDL-S); albumin (Alb); Dense lipoprotein as all particles with a density of 1.21-1.25 g/mL; dense LDL (dLDL); dense HDL-L (dHDL-L); dense HDL-M (dHDL-M); dense HDL-S (dHDL-S). Bars represent relative mol percentages. Identified lipid species were grouped into the lipid classes of (left to right) acyl carnitine (CAR), cholesterol ester (CE), ceramide (Cer), unesterified cholesterol, diglyceride (DG), fatty acid (FA), lyso-phosphatidyl choline (LPC), lyso-phosphatidyl ethanolamine (LPE), phosphatidyl choline (PC), phosphatidyl ethanolamine (PE), phosphatidyl glycerol (PG), phosphatidyl inositol (PI), sphingomyelin (SM), sphingosine, and triglyceride (TG).

**Figure 10.**
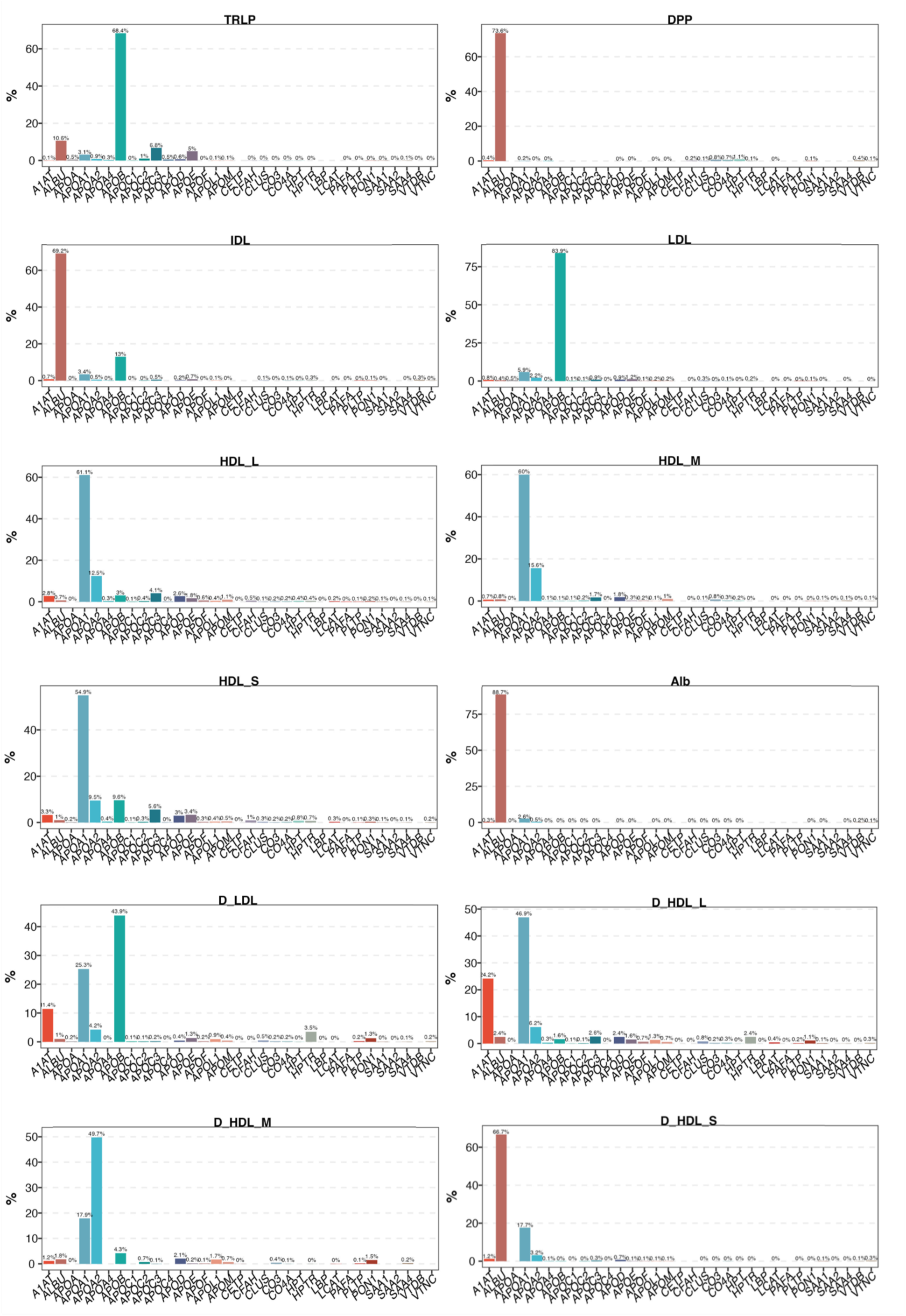
Shotgun proteomics of isolated fractions from the plasma pool, demonstrating that the most abundant proteins of each fraction are associated with the respective particle of interest. Fractions analyzed include triglyceride rich lipoprotein (TRLP); dense plasma protein (DPP) as all particles with a density > 1.25 g/mL; intermediate density lipoprotein (IDL); low density lipoprotein (LDL); high density lipoprotein large (HDL-L); high density lipoprotein medium (HDL-M); high density lipoprotein small (HDL-S); albumin (Alb); dense LDL (dLDL); dense HDL-L (dHDL-L); dense HDL-M (dHDL-M); dense HDL-S (dHDL-S). Protein abundance is represented as percentage (%) of total ion counts for each sample.

Apolipoprotein A-I (ApoA-I) was the predominant protein across the HDL fractions, however, it was lower in relative abundance in the VHDL compared to classic density HDL (60-70% vs. 40%). Apolipoprotein A-II (ApoA-II) was highly concentrated in the dHDL-M fraction, comprising 49.7% of total protein, compared to just 15.6% in HDL-M. Additionally, the relative abundance of alpha-1-antitrypsin (A1AT) in the VHDL fractions was higher than in classic density HDL. Though in low abundance, lipopolysaccharide-binding protein (LBP), cholesteryl ester transfer protein (CETP), and phospholipid transfer protein (PLTP), were enriched in all VHDL fractions as shown in Supplemental Table 5.

RNA sequencing results indicate that overall fraction sRNAs associated with enriched LPP fractions mapped to the utilized reference transcripts with 50% or higher efficiency with a zero mismatch tolerance, and approximately with 70% efficiency or higher when 1 mismatch was allowed. (Figure 11A, Table S5 and S6). The sRNA reads mapped to the human genome (host) were most abundant in the DPP fraction, followed by the TLRP, dHDL-S, Alb, and HDL-M fractions (Figure 11A, Table S5 and S6). Conversely, the largest proportions of sRNAs in the IDL, LDL, HDL-L, dLDL, dHDL-L, and dHDL-M fractions mapped to non-host small and large rRNA subunits, from SILVA rRNA reference database (release 138.2), with large sub-unit (LSU) (23S/28S) being the most prevalent, followed by small sub-unit (SSU) (16S/18S). The length distribution of the aforementioned rRNA reads was drastically different between the fractions indicating that those reads does not derive from contamination during RNA isolation or library preparation for RNA-seq (Figure 11B). In addition, correlation analysis using log-count-per million (lcpm) of foreign rRNA reads indicated strong dissimilarity in the content of rRNA between fractions (Figure 11C, Figure 11D). Finally, MDS method was used to assess similarities and differences in the non-host sRNA populations across different LPP fractions (Figures 12).

**Figure 11.**
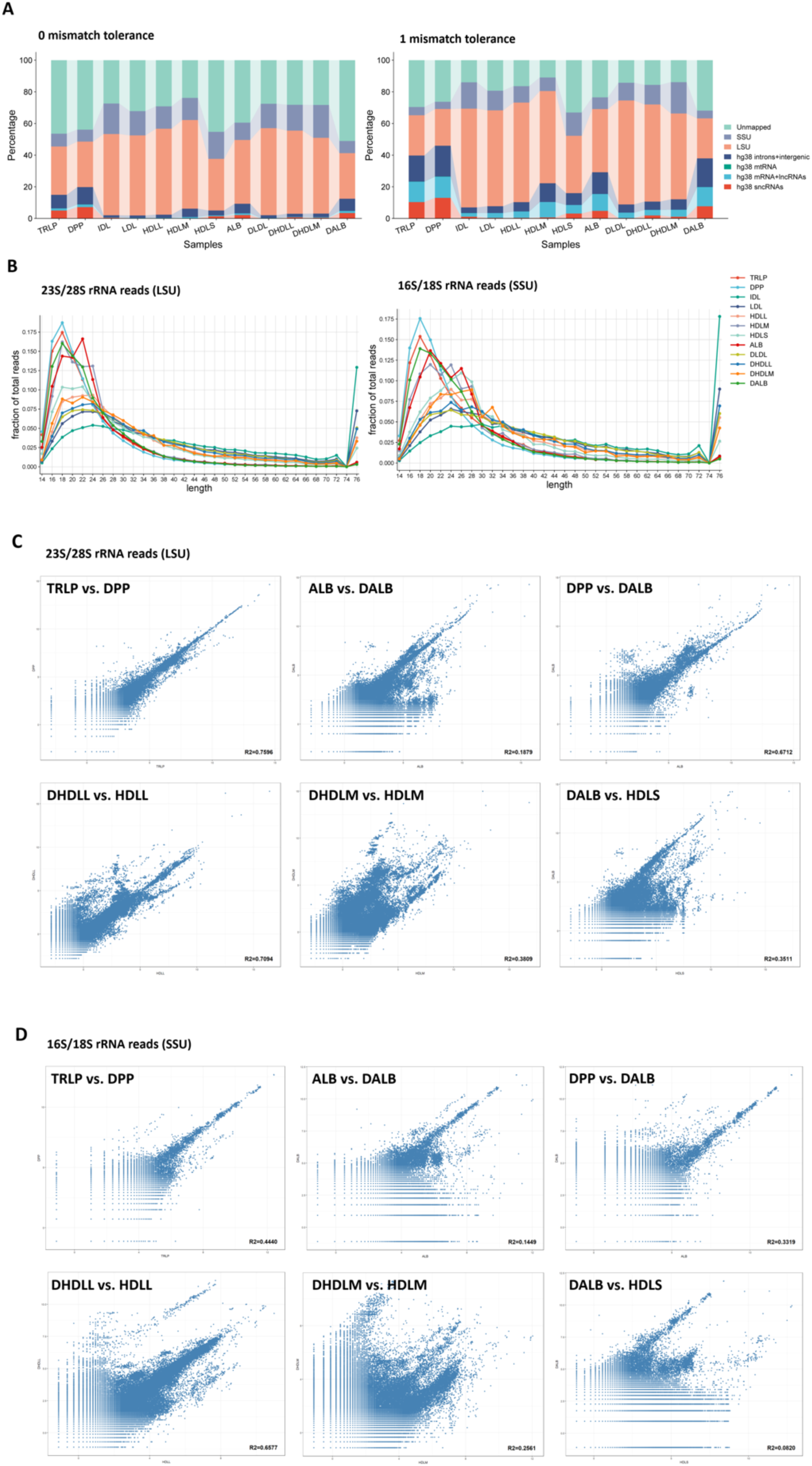
RNA sequencing analysis of isolated fractions. a) Percentages of reads that were unmapped, and mapped to host (hg38) as well as non-host small rRNA subunit (SSU) and large rRNA subunit (LSU); b) Length distribution of reads mapped to non-host small rRNA subunit (SSU) and large rRNA subunit (LSU) references across all samples; c) and d) Correlation plots of log-count-per million (lcmp) values of reads mapped to large and small ribosomal RNA subunits from various species, respectively. Each dot represents a distinct organism (specie). Correlation coefficient (R^2^) values are reported at the bottom right corner of each correlation plot in b) and c). Fractions analyzed include triglyceride rich lipoprotein (TRLP); dense plasma protein (DPP) as all particles with a density > 1.25 g/mL; intermediate density lipoprotein (IDL); low density lipoprotein (LDL); high density lipoprotein large (HDLL); high density lipoprotein medium (HDLM); high density lipoprotein small (HDLS); albumin (ALB); dense LDL (DLDL); dense HDL-L (DHDLL); dense HDL-M (DHDLM); dense HDL-S (DHDLS).

**Figure 12.**
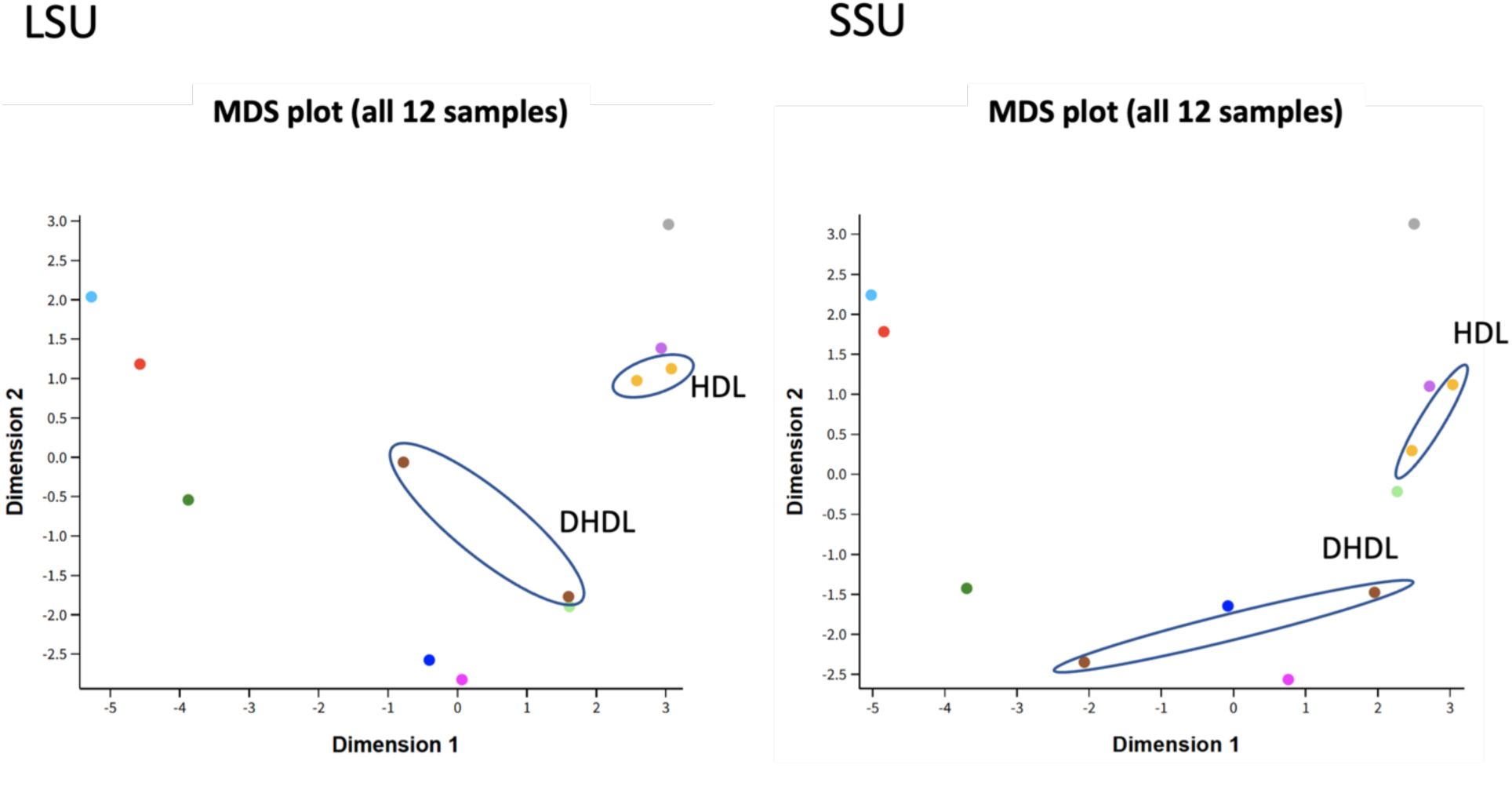
Differential gene expression analysis shown in multiple dimension scaling (MDS) plots of RNA sequenced from isolated fractions. SSU, small 16S/18S subunit of ribosomal RNA; LSU, large 23S/28S subunit of ribosomal RNA. Ellipses are drawn around the non-dense high-density lipoprotein (HDL) and dense HDL (DHL) fractions to highlight their compositional differences in non-human ribosomal RNA.

Interestingly, correlations of LSU sRNAs between HDL-L, HDL-M, and HDL-S and their corresponding denser fractions (dHDL-L, dHDL-M, and dHDL-S) were stronger for HDL-L vs. dHDL-L (R² = 0.7094) and weaker for HDL-M vs. dHDL-M (R² = 0.3809) and HDL-S vs. dHDL-S (R² = 0.3511) (Figure 11b). Correlations of SSU sRNAs between these fractions were weaker, with R² values of 0.6577 (HDL-L vs. dHDL-L), 0.2561 (HDL-M vs. dHDL-M), and 0.0820 (HDL-S vs. dHDL-S) (Figure 11c). Notably, the dHDL-S fraction exhibited a stronger correlation with the DPP fraction compared to the Alb fraction. Multidimensional scaling (MDS) plots in Figure 12 reveal distinct clustering of RNA-seq data, with HDL fractions positioned further from dHDL fractions.

Additionally, greater distances were observed between dHDL samples compared to HDL samples.

## Discussion

Results from NDGE, WB, EM and NMR analyses show that our method successfully isolates and enriches the target LPP particles in each fraction. Differences in the methods used for size measurement of LPP have been thoroughly discussed^24,25^: whereas NMR measures the resonance of the methyl group in the LPP core, negative stain EM measures the edge to edge diameter of the LPP structure. Fractions with the highest abundance of contaminating particles were the IDL fraction, which had very few particles in it, HDL-L, which had some LDL contamination, Alb, which had some small HDL left over from the HDL-S fraction, and conversely, HDL-S, which had albumin contamination, and dHDL-S, which was predominantly comprised of albumin. The coelution of albumin and HDL-S particles, which share a similar Rh, affects the accuracy of MALS-derived Mw calculations. Due to the high abundance of albumin, its signal disproportionately influences the Mw calculation. As a result, alternative analysis methods provide a more precise characterization of the particles and protein composition in these fractions (HDL-S and Alb). This highlights the importance of using orthogonal analytical methods to capture different aspects of the LPP isolated with our approach. EM confirmed the physical characteristics of LPP across fractions, while WB and proteomics provided sensitive measures of potential contamination and particle identity. NMR offered comparative insights into the size distribution of LPP subspecies, and NDGE resolved particle size based on Mw. By integrating these perspectives, we gained both detailed and broader views of the LPP in each fraction, which facilitated comparisons with existing literature ^24–27^.

Results from NDGE of the LPP fractions up to 1.21 g/mL in density show particle size distributions within the expected ranges (Figure 5)^28,29^, with the exception of the IDL fraction. Figure 6 shows the NDGE of the IDL fraction containing particles of ∼ 160 kDa and <116 kDa, though the bands are faint. Upon further investigation, we found that samples remaining in the fraction collector tube post Alb fraction collection were dispensed to the following sample.

Thus, the IDL fraction was contaminated with trace amounts of the previous sample’s albumin. All other fractions that were collected followed the expected decrease in particle size over the SEC elution time. HDL-L showed minor co-elution of particles from the LDL fraction, and HDL-S also showed co-elution of particles <116 kDa, similar to particles in the Alb fraction (Figure 6). In the HDL-S fraction, particles between 116-160 kDa could signify particles that are known in the literature as HDL3^28^ while particles below 116 kDa, could signify the presence of the common protein contaminant, human serum albumin^12^ (69 kDa) (Uniprot).

The dense LPP fractions were different from the non-dense fractions, as shown by NDGE (Figure 6). Both dLDL and dHDL-L contained particles slightly larger than 500 kDa, while the LDL and HDL-L fractions had significantly larger particles than the highest Mw band, suggesting that particles like LDL and other TRLP were present^30^. Bands between 16–160 kDa were detected in all dense LPP fractions but were detected only in the HDL and Alb fractions in the non-dense fractions. The particle size distribution in dHDL-M resembled that of HDL-M, but with more staining between 16–160 kDa compared to HDL-M, which contained mostly particles in the >160 kDa range. dHDL-S showed dense staining at 160 kDa, along with smaller particles and a faint band above 160 kDa. EM revealed similar results, showing that dLDL and dHDL-L particles were smaller in diameter than LDL and HDL-L, suggesting that particles in the dLDL and dHDL-L fractions are generally smaller in diameter and have a lower Mw (Figure 7). Additionally, dHDL-M and dHDL-S, like HDL-M and Alb, had particles under 10 nm but exhibited greater size variability. NMR results of dense LPP fractions were not available due to limitation in sample yield post fractionation.

WB results suggest that the expected protein was most abundant in each fraction^31^ (Figure 5). Both the TRLP and LDL fractions were primarily composed of ApoB-100. HDL-L, and to a lesser extent HDL-M, also contained ApoB-100, which was likely LDL particle co-elution in the fraction, as explained by the presence of particles > 500 kDa in the NDGE (Figure 6). However, the content of ApoB-100 was only 3% of the total protein content in HDL-L and less than 1% of total protein content in HDL-M as shown by proteomics data. These data show that LDL particles co-eluted into the HDL-L fraction using the cutoffs currently selected. Depending on the interest of the investigator in characterizing the LDL particles using the light scattering instruments vs. improving the purity of the HDL fractions, method adjustments can be made to resolve this issue. If the goal is to increase the purity of the HDL-L fraction but still be able to characterize LDL by LS, the simplest approach is to adjust the fractionation cut-off farther away from the LDL trough and farther toward the HDL-L peak to minimize the amount of carryover of LDL. If the goal is to increase the HDL-L purity and LDL are of no interest for LS characterization purposes, another approach is to adjust the density to exclude LDL particles all together from the SEC by bringing up the first UC density cut off to 1.063 g/mL during the UC preparatory step^30^.

While the IDL fraction would typically contain ApoB, the coelution of particles from the preceding Alb fraction explains why albumin was the predominant protein there (Figure 10 - though it is important to note that WB of all protein bands in the IDL lane were very faint, suggesting that there was not much protein present in that fraction to begin with (Figure 5). This was as expected given that the plasma was from healthy individuals, rather than those with dyslipidemia, who would not be expected to have remnant LPP circulating in the fasted state.

Proteomics results show that HDL-L and HDL-M had the expected HDL protein composition, with ApoA-I accounting for about 60-70% and ApoA-II for 12-16% of total protein (Figure 10) ^7,8,12^. A broader range of apoproteins, including ApoC-III, ApoE, and ApoD, constituted a higher percentage in HDL-L compared to HDL-M, consistent with existing literature on larger versus smaller HDL particles^12,32^. The dLDL fraction contained about 44% ApoB, 25% ApoA1, and 11% A1AT, corresponded to LDL size (20-25nm in diameter) as measured by EM and an Rh of 12.62nm as measured by DLS (Figure 7, Table 6). The dLDL migrated at around 500 kDa and larger by NDGE and had a Mw of 2,574 kDa by MALS (Figure 6, Table 6). Further investigation is needed to explore the role of these particles and whether they are a specific subclass of LPP as these particles have a higher density than the densest LDL, whose density is 1.044–1.063 g/mL^33^ as well as a different protein composition compared to classic LDL particles (Figure 7). dHDL-M contained predominantly ApoA-II at 50%, followed by ApoA-I at 18%, a unique composition unlike non-dense HDL-M. The dHDL-L fraction contained about 47% ApoA-I and 24% A1AT, which is very different from the proteomic composition of non-dense HDL-L. The VHDL fractions were particularly enriched in A1AT, apolipoprotein L-1 (ApoL-1), PON1, PLTP, and LBP compared to non-dense HDL, suggesting that these are “specialty” HDL particles which likely have distinct functions in addition to cholesterol efflux, such as functions involved in innate immunity^34^ (Supplemental Table S8).

Lipidomics of the non-dense HDL fractions show values that generally concur with the published literature^22,35^. Literature values for non-dense HDL particles include 32-45 mol% PC, 30-40 mol% CE, 3-12 mol% TG, 5-10 mol% SM, 0.5-8% LPC, and 5-10 mol% unesterified cholesterol^35,36^. Compared to those values, the lipid composition in the HDL-L, HDL-M, and HDL-S fractions (Figure 9) are higher in PC (46-53 mol%) and SM (10-12 mol%), lower in CE (17-26 mol%), and comparable in TG (5-10 mol%), SM (10-12 mol%), and LPC (0.9-1.4 mol%). Unesterified cholesterol values were lower than literature values in HDL-L and HDL-M (3 mol% and 1.2 mol%) but comparable for HDL-S (11 mol%)^37^. Trends for changes in relative proportions of lipid classes over the HDL-L, HDL-M, and HDL-S fractions agree with current literature including decreasing CE and TG relative abundance from large to small particles^36^. These changes reflect the decrease in the size of the neutral lipid core from large to small particles. Lipidomic analysis of the VHDL fractions (Figure 9) showed a higher relative abundance of unesterified cholesterol with values double what has been previously reported for non-dense HDL (10-25 mol% vs. literature 5-10 mol%). VHDL particles also had about five times higher relative proportions of FA (7-16 mol% vs literature 0.5-8 mol%) compared to the literature values and values reported here for HDL particles in the classic density range^35,36^. VHDL additionally contained lower relative proportions of CE (14-16 mol%), and about the same relative abundances of TG (4-6 mol%), and SM (8 mol%) compared to literature values of HDL particles in the classic density range (literature CE 30-40 mol%, TG 3-12 mol%, and SM 5-10 mol%) (Figure 9)^35,36^. The relative abundances of PC in VHDL, though comparable to literature values, were about 10% lower in VHDL than non-dense HDL reported here (Figure 9). The enrichment of FC and decrease in PC is indicative of the lipid composition of lipid rafts^38^ which may lead to higher association of raft-associated proteins in VHDL particles.

To decrease ambiguous mapping between different sRNA biotypes, RNA sequencing data of the LPP fractions were analyzed using sequential mapping strategy described previously by several authors^39,40^ in combination with custom curated human transcriptome references. The additional step in RNA-seq analysis included mapping of all non-hg38 reads to SILVA database of small and large rRNA subunits for all three domains of life (Bacteria, Archaea and Eukarya)^41^. Our findings showed the presence of mostly non-host RNA in the fractions, which aligns with previous reports^4,42^. Correlation analysis of sRNAs between non-dense LPP fractions and dense LPP fractions revealed low R² values, indicating differential sRNA associations between these groups (Figures 11). This finding is further supported by the MDS plot from the DGE analysis (Figure 12), which demonstrates clear separation between HDL and dHDL fractions, highlighting distinct sRNA profiles. Moreover, the dHDL-L and dHDL-M fractions exhibited greater variability in RNA expression patterns compared to HDL-L and HDL-M fractions, suggesting that dense LPP fractions transport a unique set of RNAs and are more heterogeneous in their sRNA composition.

Intestinally-derived particles, which account for as much as 30% of the circulating HDL pool^43^, are smaller^44^, denser and have a higher protein content than liver-derived HDL^43,45^, and have been shown to protect the liver from inflammation by binding lipopolysaccharide (LPS) derived from the gut via LBP^44,46^ However, intestinally-derived HDL have only been studied in humans in small studies involving stable isotope tracers^45^. There are currently no available methods to study these particles directly in human samples. It is very likely that VHDL, which are denser than “classic density HDL” due to a higher protein content, are enriched in intestinally-derived HDL. Our finding that dHDL-M particles are more highly enriched in gut microbiome-derived sRNA than all other plasma and LPP fractions and that their sRNA profile is dissimilar to that of classic-density HDL provide further evidence that VHDL are likely enriched in intestinally-derived HDL.

The LS and dRI detectors allow for on-line physicochemical characterization of particles as they elute through the instrument. The LS detectors are inordinately more sensitive to larger particles compared to smaller particles^26^. The intensity of light scattering is dependent on the diameter of the particle to the 6^th^ power, such that an average sized LDL particle scatters approximately 1,000 times more light than an average sized HDL particle. Thus, particles measured between peaks show larger size measurements due to the elution of trace amounts of ApoB containing particles still eluting from the void volume and LDL peaks. The elution of trace amounts of ApoB containing particles throughout the HDL elution size range could be due to particle-particle charge or biological interactions, and hydrophobic column adsorption ^47^.

However, when enough particles are provided in each LS instrument cell (when chromatograms show peak elution of particles) signals from the particles provide ideal correlation and can provide measurements with lower variability, comparable to both NDGE (Mw), EM (Radius), and NMR. As mentioned previously, one way to increase the resolution of the measurements and decrease contamination from ApoB containing particles would be to remove the ApoB particles completely from the injected fraction by increasing the density of the solution in the UC preparatory step, or by using filtration to eliminate larger particles.

We demonstrated the ability of the LS, UV, and dRI detectors to measure low concentration samples in the dense LPP fractions (1.21-1.25 g/mL), when other post processing analysis steps could not provide analysis results due to low sample concentrations. In these samples, dHDL-L had larger average Rh than non-dense HDL-L (7.090nm in HDL-L vs 8.145nm in DHDL-L). Additionally, though dHDL-M and HDL-M have Rh that were close to one another (6.089 vs 5.983nm), HDL-M had a higher Mw than dHDL-M (177.2 kDa vs 143.1 kDa).

## Conclusion

Here we demonstrate that the UC-SEC-UV/MALS/DLS/dRI method described is a useful, effective, and versatile approach for isolating all LPP fractions in plasma, including the until recently ignored VHDL particles. We demonstrate that each fraction is enriched in the target LPP particle, with little contamination from other particle classes or plasma components. The method is versatile in that fractionation can be adjusted to either further refine the isolation of particles by size (e.g. even more HDL size-based fractions), or to further exclude any remaining contaminating particles with adjustments in fractionation cutoffs or density cutoffs during the UC preparative steps. The addition of light scattering instruments further enhances the characterization of the particles as they are fractionated, providing particle size and Mw measurements, and also the potential to calculate particle concentration. The use of this method paired with proteomics, lipidomics, and RNA sequencing of the isolated fractions revealed unique structural and compositional characteristics of the particles in the 1.21–1.25 g/mL density range. We describe for the first time the presence of dLDL particles which share the size of classic density LDL particles, but which have a unique proteome enriched in ApoA-I and A1AT. We also describe for the first time the unique compositional characteristics of VHDL particles, showing a unique lipidome with an enrichment in unesterified cholesterol and a depletion in CE, a unique proteome with depletion of ApoA-I and an enrichment in A1AT, transfer proteins like LBP, PLTP, and CETP, and a unique sRNA profile enriched in gut microbiome-derived RNA.

## Author contributions

Designed the study: A.M.Z.; supervised the study: A.M.Z., C.B.L., W.V., J.P.T, K.W.; performed experiments: J.K.A., O.G., J.Z., A.O., B.H., S.L.; analyzed data: X.T., B.H., A.T.; writing, review, and editing: J.K.A., O.G., J.Z., B.H., A.T., A.M.Z. All authors agree with the submission and have no conflicts of interest to disclose.

## Declaration of generative AI and I-assisted technologies in the writing process

During the preparation of this work the authors used ChatGPT (OpenAI) in order to improve the readability and language of the manuscript. After using this tool/service, the authors reviewed and edited the content as needed and take full responsibility for the content of the published article.

## Acknowledgements

This work was supported in part by National Institute on Aging (RO1AG062240); the National Institute of General Medical Sciences (R01 GM147545); the Common Fund’s Extracellular RNA Communications Consortium (UG3/UH3 CA241694) of the National Institutes of Health; and the USDA National Institute of Food and Agriculture, Hatch project (grant number CA-D-NUT-2242-H). We acknowledge the support from the UC Davis West Coast Metabolomics Center.

