## Supplemental Tables and Figures for "Simultaneous Isolation and Characterization of Lipoprotein Classes in Plasma, Including HDL Subclasses and the Uncharacterized Dense HDL"

**Supplementary Materials**

**Supplemental Table S1.** Plasma pool sample characteristics, including mean, standard deviation and percent coefficient of variation of concentrations of platelet, red blood cell (RBC), hemoglobin (HGB), white blood cell (WBC), cholesterol, triglyceride, low-density lipoprotein (LDL)-cholesterol (LDL-C), and high-density lipoprotein (HDL)-cholesterol (HDL-C).

|  | Platelet [K/uL] | RBC [M/uL] | HGB [g/dL] | WBC [g/dL] | Cholesterol (mg/dL) | Triglycerides (mg/dL) | LDL-C (mg/dL) | HDL-C (mg/dL) |
| --- | --- | --- | --- | --- | --- | --- | --- | --- |
| Mean | 9.500 | 0.003 | 0.011 | 1.205 | 187.778 | 110.833 | 101.591 | 64.020 |
| Standard Deviation | 16.446 | 0.005 | 0.040 | 1.108 | 39.573 | 65.793 | 31.332 | 17.046 |

**Supplemental Table S2**. Experimental constant values used for data processing through Astra 8.1.2. Fractions analyzed on Astra include intermediate density lipoprotein (IDL); low density lipoprotein (LDL); high density lipoprotein large (HDL-L); high density lipoprotein medium (HDL-M); high density lipoprotein small (HDL-S); albumin (Alb); dense LDL (dLDL); dense HDL-L (dHDL-L); dense HDL-M (dHDL-M); dense HDL-S (dHDL-S).

| Variable | Sub-variable | Value | Source |
| --- | --- | --- | --- |
| dn/dc of Protein (mL/g) | All fractions analyzed on Astra | 0.1899 | Zhao, et al., 2011 (doi: 10.1016/j.bpj.2011.03.004) |
| dn/dc of Lipid (or modifier) (mL/g) | All fractions analyzed on Astra | 0.1605 | Parkilla, et al., 2018 (doi: 10.1021/acs.langmuir.8b01259 |
| Light Scattering Model | All fractions analyzed on Astra | Zimm* | Astra |
| Fit Degree | All fractions analyzed on Astra | 1 | Astra |
| Refractive Index of Solvent | 1 X PBS | 1.331 | Astra |
| Extinction Coefficient at 280nm (mL/(mgcm)) | IDL  LDL  HDL-L  HDL-M  HDL-S  Alb  dLDL  dHDL-L  dHDL-M  dHDL-S | 0.7449  0.8396  1.0419  1.1484  0.9508  0.6074  0.8512  0.9456  0.8674  0.6719 | Extinction coefficient from UniProt sequence (Supplemental Table S3)  Proteomic protein composition of top 28 most abundant protein, including Albumin and ApoB100 (Supplemental Table S4)  Gill and Von Hippel, 1989 (doi: https://doi.org/10.1016/0003-2697(89)90602-7) |

**Supplemental Table S3.** The 28 most abundant HDL associated proteins along with Apolipoprotein B-100 and Albumin and their UniProt identifier for extinction coefficient analysis.

| **Protein Record Name** | **Protein** | **UniProt ID** | **Number of Amino Acids** | **Molecular Weight (Da)** | **Extinction Coefficient (M-1 cm-1)** | **notes** |
| --- | --- | --- | --- | --- | --- | --- |
| ApoA-I | APOA1 | P02647 | 267 | 30777.83 | 37930 | NA |
| ApoA-II | APOA2 | P02652 | 100 | 11175.02 | 6085 | assuming all pairs of Cys residues form cystines |
| ApoD | APOD | P05090 | 189 | 21275.55 | 32680 | assuming all pairs of Cys residues form cystines |
| ApoC-III | APOC3 | P02656 | 99 | 10852.31 | 19480 | NA |
| ApoM | APOM | O95445 | 188 | 21253.29 | 27305 | assuming all pairs of Cys residues form cystines |
| PON1 | PON1 | P27169 | 355 | 39731.31 | 47455 | assuming all pairs of Cys residues form cystines |
| ApoE | APOE | P02649 | 317 | 36154.08 | 50085 | assuming all pairs of Cys residues form cystines |
| Vitronectin | VTNC | P04004 | 478 | 54305.59 | 90145 | assuming all pairs of Cys residues form cystines |
| Serum paraoxonase/lactonase 3 | PON3 | Q15166 | 354 | 39607.49 | 32110 | assuming all pairs of Cys residues form cystines |
| Apolipoprotein F | APOF | Q13790 | 326 | 35399.49 | 34755 | assuming all pairs of Cys residues form cystines |
| Apolipoprotein C-II | APOC2 | P02655 | 101 | 11283.87 | 12950 | NA |
| Apolipoprotein L1 | APOL1 | O14791 | 398 | 43974.2 | 48930 | assuming all pairs of Cys residues form cystines |
| A1AT | A1AT | P01009 | 418 | 46736.55 | 25565 | assuming all pairs of Cys residues form cystines |
| Serum amyloid A-4 protein | SAA4 | P35542 | 130 | 14746.65 | 32555 | assuming all pairs of Cys residues form cystines |
| Complement C3 | CO3 | P01024 | 1663 | 187148.06 | 180055 | assuming all pairs of Cys residues form cystines |
| Apolipoprotein C-I | APOC1 | P02654 | 83 | 9331.93 | 5500 | NA |
| LCAT | LCAT | P04180 | 440 | 49577.92 | 107175 | assuming all pairs of Cys residues form cystines |
| Complement C4-A | CO4A | P0C0L4 | 1744 | 192785.48 | 189230 | assuming all pairs of Cys residues form cystines |
| ApoJ | CLUS | P10909 | 449 | 52494.58 | 44515 | assuming all pairs of Cys residues form cystines |
| Haptoglobin-related protein | HPTR | P00739 | 348 | 39029.57 | 71320 | assuming all pairs of Cys residues form cystines |
| ApoA-IV | APOA4 | P06727 | 396 | 45372.04 | 15930 | NA |
| SAA1 | SAA1 | P0DJI8 | 122 | 13532.02 | 23950 | assuming all pairs of Cys residues form cystines |
| PLTP | PLTP | P55058 | 493 | 54739.44 | 34630 | assuming all pairs of Cys residues form cystines |
| Haptoglobin | HPT | P00738 | 406 | 45205.31 | 76040 | assuming all pairs of Cys residues form cystines |
| Antithrombin-III | ANT3 | P01008 | 464 | 52602.44 | 45880 | assuming all pairs of Cys residues form cystines |
| Apolipoprotein C-IV | APOC4 | P55056 | 127 | 14553.05 | 30855 | assuming all pairs of Cys residues form cystines |
| Ceruloplasmin | CERU | P00450 | 1065 | 122205.19 | 200735 | assuming all pairs of Cys residues form cystines |
| CETP | CETP | P11597 | 493 | 54756.21 | 42775 | assuming all pairs of Cys residues form cystines |
| Albumin | ALB | P02768 | 609 | 69366.68 | 41435 | assuming all pairs of Cys residues form cystines |
| Apolipoprotein B | APOB | P04114 | 4563 | 515545 | 431480 | assuming all pairs of Cys residues form cystines |

**Supplemental Table S4.** Composition of proteins identified by proteomics analysis of the top 28 most abundant HDL-associated proteins along with Apolipoprotein B-100 and Albumin. Fractions analyzed include intermediate density lipoprotein (IDL); low density lipoprotein (LDL); high density lipoprotein large (HDL-L); high density lipoprotein medium (HDL-M); high density lipoprotein small (HDL-S); albumin (Alb); dense LDL (dLDL); dense HDL-L (dHDL-L); dense HDL-M (dHDL-M); dense HDL-S (dHDL-S).

| **Protein** | **IDL** | **LDL** | **HDL-L** | **HDL-M** | **HDL-S** | **Alb** | **dLDL** | **dHDL-L** | **dHDL-M** | **dHDL-S** |
| --- | --- | --- | --- | --- | --- | --- | --- | --- | --- | --- |
| A1AT | 0.80% | 0.79% | 3.00% | 0.83% | 3.41% | 0.33% | 11.81% | 24.91% | 1.39% | 1.32% |
| ALB | 77.57% | 0.37% | 0.75% | 0.93% | 1.06% | 95.90% | 0.99% | 2.50% | 2.21% | 73.35% |
| ANT3 | 0.09% | 0.00% | 0.00% | 0.04% | 0.00% | 0.13% | 0.15% | 0.10% | 0.00% | 0.01% |
| APOA1 | 3.79% | 6.06% | 65.21% | 70.75% | 57.42% | 2.83% | 26.20% | 48.30% | 21.58% | 19.44% |
| APOA2 | 0.57% | 2.24% | 13.32% | 18.41% | 9.94% | 0.55% | 4.38% | 6.34% | 59.87% | 3.52% |
| APOA4 | 0.04% | 0.05% | 0.31% | 0.14% | 0.37% | 0.04% | 0.03% | 0.32% | 0.00% | 0.07% |
| APOB | 14.58% | 85.79% | 3.21% | 0.09% | 10.07% | 0.00% | 45.45% | 1.65% | 5.12% | 0.04% |
| APOC1 | 0.02% | 0.15% | 0.07% | 0.12% | 0.12% | 0.01% | 0.12% | 0.06% | 0.00% | 0.02% |
| APOC2 | 0.03% | 0.09% | 0.39% | 0.26% | 0.35% | 0.01% | 0.12% | 0.12% | 0.90% | 0.05% |
| APOC3 | 0.54% | 0.93% | 4.40% | 2.04% | 5.83% | 0.00% | 0.24% | 2.65% | 0.10% | 0.38% |
| APOC4 | 0.00% | 0.01% | 0.04% | 0.01% | 0.03% | 0.00% | 0.01% | 0.02% | 0.00% | 0.00% |
| APOD | 0.28% | 0.97% | 2.83% | 2.09% | 3.10% | 0.01% | 0.45% | 2.50% | 2.52% | 0.81% |
| APOE | 0.81% | 1.24% | 1.88% | 0.30% | 3.55% | 0.00% | 1.40% | 1.60% | 0.25% | 0.11% |
| APOF | 0.02% | 0.07% | 0.69% | 0.19% | 0.31% | 0.00% | 0.22% | 0.73% | 0.16% | 0.11% |
| APOL1 | 0.08% | 0.20% | 0.42% | 0.14% | 0.46% | 0.00% | 0.96% | 1.35% | 2.09% | 0.07% |
| APOM | 0.01% | 0.17% | 1.13% | 1.15% | 0.55% | 0.00% | 0.40% | 0.67% | 0.85% | 0.12% |
| CERU | 0.00% | 0.00% | 0.01% | 0.29% | 0.00% | 0.03% | 0.01% | 0.00% | 0.00% | 0.00% |
| CETP | 0.00% | 0.00% | 0.00% | 0.00% | 0.01% | 0.00% | 0.00% | 0.00% | 0.05% | 0.00% |
| CLUS | 0.16% | 0.26% | 0.15% | 0.11% | 0.34% | 0.02% | 0.48% | 0.84% | 0.00% | 0.00% |
| CO3 | 0.02% | 0.04% | 0.23% | 0.93% | 0.18% | 0.01% | 0.21% | 0.23% | 0.51% | 0.02% |
| CO4A | 0.06% | 0.07% | 0.26% | 0.40% | 0.26% | 0.02% | 0.22% | 0.33% | 0.06% | 0.03% |
| HPT | 0.05% | 0.03% | 0.42% | 0.22% | 0.80% | 0.00% | 0.03% | 0.03% | 0.00% | 0.00% |
| HPTR | 0.30% | 0.24% | 0.43% | 0.01% | 0.75% | 0.00% | 3.61% | 2.47% | 0.06% | 0.01% |
| LCAT | 0.00% | 0.00% | 0.25% | 0.03% | 0.35% | 0.00% | 0.01% | 0.43% | 0.03% | 0.02% |
| PLTP | 0.06% | 0.05% | 0.07% | 0.00% | 0.09% | 0.00% | 0.25% | 0.20% | 0.16% | 0.00% |
| PON1 | 0.09% | 0.11% | 0.20% | 0.04% | 0.26% | 0.01% | 1.34% | 1.14% | 1.83% | 0.03% |
| PON3 | 0.00% | 0.00% | 0.02% | 0.00% | 0.02% | 0.00% | 0.54% | 0.10% | 0.00% | 0.00% |
| SAA1 | 0.00% | 0.03% | 0.14% | 0.16% | 0.09% | 0.00% | 0.05% | 0.13% | 0.00% | 0.06% |
| SAA4 | 0.00% | 0.02% | 0.06% | 0.18% | 0.07% | 0.00% | 0.10% | 0.03% | 0.27% | 0.01% |
| VTNC | 0.03% | 0.03% | 0.12% | 0.12% | 0.18% | 0.09% | 0.20% | 0.26% | 0.00% | 0.38% |

**Supplemental Table S5**. RNA sequencing mapping statistics for both “No mismatches allowed” or 0 mismatch tolerance and “Maximum 1 mismatch allowed” or 1 mismatch tolerance. Fractions analyzed include intermediate density lipoprotein (IDL); low density lipoprotein (LDL); high density lipoprotein large (HDL-L); high density lipoprotein medium (HDL-M); high density lipoprotein small (HDL-S); albumin (Alb); dense LDL (dLDL); dense HDL-L (dHDL-L); dense HDL-M (dHDL-M); dense HDL-S (dHDL-S).

| **No mismatches allowed** | **TRLP** | **DPP** | **IDL** | **LDL** | **HDL-L** | **HDL-M** | **HDL-S** | **Alb** |
| --- | --- | --- | --- | --- | --- | --- | --- | --- |
| Total_reads | 14140603 | 20707320 | 44623463 | 51756564 | 42766481 | 39426839 | 35415435 | 29326208 |
| Reads ≥ 15 nt | 8472677 | 12364783 | 41184472 | 48387315 | 39824515 | 32929899 | 30146755 | 19563008 |
| % Reads ≥ 15 nt | 59.92% | 59.71% | 92.29% | 93.49% | 93.12% | 83.52% | 85.12% | 66.71% |
| mtRNA | 0.02% | 0.07% | 0.00% | 0.00% | 0.00% | 0.00% | 0.00% | 0.01% |
| tRNA | 0.08% | 0.14% | 0.01% | 0.01% | 0.00% | 0.00% | 0.02% | 0.03% |
| RN7S | 0.01% | 0.02% | 0.00% | 0.00% | 0.00% | 0.00% | 0.00% | 0.00% |
| RNU | 0.13% | 0.49% | 0.01% | 0.00% | 0.00% | 0.00% | 0.01% | 0.02% |
| snoRNA& scaRNA | 0.01% | 0.02% | 0.00% | 0.00% | 0.00% | 0.00% | 0.00% | 0.00% |
| vault RNA | 0.00% | 0.00% | 0.00% | 0.00% | 0.00% | 0.00% | 0.00% | 0.00% |
| RNY | 0.02% | 0.36% | 0.00% | 0.00% | 0.00% | 0.00% | 0.00% | 0.01% |
| rRNA | 4.77% | 6.06% | 0.46% | 0.11% | 0.12% | 0.18% | 1.37% | 2.10% |
| Combined_tRNA_to_rRNA | 5.01% | 7.09% | 0.48% | 0.12% | 0.13% | 0.19% | 1.39% | 2.16% |
| miRNA | 0.01% | 0.10% | 0.00% | 0.00% | 0.00% | 0.00% | 0.00% | 0.00% |
| premiRNA_only | 0.00% | 0.00% | 0.00% | 0.00% | 0.00% | 0.00% | 0.00% | 0.00% |
| protein coding (PC) mRNA | 0.69% | 0.86% | 0.10% | 0.13% | 0.15% | 0.41% | 0.28% | 0.60% |
| lncRNAs | 0.12% | 0.14% | 0.02% | 0.03% | 0.03% | 0.08% | 0.06% | 0.11% |
| other ncRNA (rest transcriptome) | 0.53% | 0.60% | 0.06% | 0.08% | 0.08% | 0.23% | 0.18% | 0.34% |
| rest hg38 (introns & intergenic) | 8.55% | 10.97% | 1.34% | 1.54% | 2.00% | 5.28% | 3.12% | 6.07% |
| hg38 total | 14.93% | 19.84% | 2.01% | 1.91% | 2.39% | 6.20% | 5.04% | 9.29% |
| no hg38 | 85.07% | 80.16% | 97.99% | 98.09% | 97.61% | 93.80% | 94.96% | 90.71% |
| LSU | 30.53% | 28.74% | 51.35% | 50.56% | 54.33% | 56.01% | 32.64% | 40.32% |
| SSU | 8.12% | 7.55% | 19.26% | 15.36% | 14.12% | 13.97% | 17.09% | 10.96% |
| **Maximum 1 mismatch allowed** | **1_S1** | **2_S2** | **3_S3** | **4_S4** | **5_S5** | **6_S6** | **7_S7** | **8_S8** |
| Total_reads | 14140603 | 20707320 | 44623463 | 51756564 | 42766481 | 39426839 | 35415435 | 29326208 |
| Reads ≥ 15 nt | 8472677 | 12364783 | 41184472 | 48387315 | 39824515 | 32929899 | 30146755 | 19563008 |
| % Reads ≥ 15 nt | 59.92% | 59.71% | 92.29% | 93.49% | 93.12% | 83.52% | 85.12% | 66.71% |
| mtRNA | 0.07% | 0.13% | 0.01% | 0.01% | 0.01% | 0.01% | 0.01% | 0.03% |
| tRNA | 0.28% | 0.35% | 0.02% | 0.04% | 0.02% | 0.04% | 0.07% | 0.12% |
| RN7S | 0.13% | 0.18% | 0.02% | 0.02% | 0.03% | 0.06% | 0.04% | 0.10% |
| RNU | 0.20% | 0.63% | 0.02% | 0.01% | 0.02% | 0.03% | 0.03% | 0.06% |
| snoRNA& scaRNA | 0.05% | 0.07% | 0.01% | 0.01% | 0.01% | 0.02% | 0.01% | 0.03% |
| vault RNA | 0.00% | 0.00% | 0.00% | 0.00% | 0.00% | 0.00% | 0.00% | 0.00% |
| RNY | 0.03% | 0.43% | 0.00% | 0.00% | 0.00% | 0.00% | 0.00% | 0.01% |
| rRNA | 9.65% | 11.19% | 1.03% | 0.26% | 0.32% | 0.51% | 2.94% | 4.40% |
| Combined_tRNA_to_rRNA | 10.32% | 12.81% | 1.09% | 0.34% | 0.39% | 0.64% | 3.08% | 4.68% |
| miRNA | 0.04% | 0.16% | 0.01% | 0.01% | 0.01% | 0.02% | 0.01% | 0.06% |
| premiRNA_only | 0.02% | 0.03% | 0.01% | 0.01% | 0.01% | 0.02% | 0.02% | 0.05% |
| protein coding (PC) mRNA | 9.63% | 9.96% | 1.68% | 2.25% | 2.94% | 7.48% | 4.06% | 8.13% |
| lncRNAs | 0.92% | 0.99% | 0.18% | 0.22% | 0.27% | 0.60% | 0.43% | 0.79% |
| other ncRNA (rest transcriptome) | 2.32% | 2.47% | 0.45% | 0.60% | 0.82% | 1.64% | 0.92% | 1.73% |
| rest hg38 (introns & intergenic) | 16.48% | 19.43% | 3.57% | 4.31% | 5.89% | 11.84% | 7.45% | 13.75% |
| hg38 total | 39.79% | 45.96% | 6.99% | 7.74% | 10.32% | 22.24% | 16.00% | 29.22% |
| no hg38 | 60.21% | 54.04% | 93.01% | 92.26% | 89.68% | 77.76% | 84.00% | 70.78% |
| LSU | 25.43% | 23.21% | 62.43% | 60.58% | 62.92% | 58.29% | 36.22% | 39.92% |
| SSU | 5.20% | 4.57% | 16.61% | 12.44% | 10.37% | 8.56% | 14.74% | 7.40% |

**Supplemental Table S6**. RNA sequencing mapping statistics for both “No mismatches allowed” or 0 mismatch tolerance and “Maximum 1 mismatch allowed” or 1 mismatch tolerance. Fractions analyzed include dense low density lipoprotein (dLDL); dense high density lipoprotein large (dHDL-L); dense HDL medium (dHDL-M); dense HDL small (dHDL-S).

| **No mismatches allowed** | **dLDL** | **dHDL-L** | **dHDL-M** | **dHDL-S** | **dLDL** | **dHDL-L** | **dHDL-M** | **dHDL-S** |
| --- | --- | --- | --- | --- | --- | --- | --- | --- |
| Total_reads | 46852616 | 44077870 | 33132431 | 17010258 | 46852616 | 44077870 | 33132431 | 17010258 |
| Reads ≥ 15 nt | 43531434 | 41202261 | 30070646 | 9744602 | 43531434 | 41202261 | 30070646 | 9744602 |
| % Reads ≥ 15 nt | 92.91% | 93.48% | 90.76% | 57.29% | 92.91% | 93.48% | 90.76% | 57.29% |
| mtRNA | 0.00% | 0.00% | 0.00% | 0.01% | 0.00% | 0.00% | 0.00% | 0.01% |
| tRNA | 0.00% | 0.00% | 0.01% | 0.04% | 0.00% | 0.00% | 0.01% | 0.04% |
| RN7S | 0.00% | 0.00% | 0.00% | 0.00% | 0.00% | 0.00% | 0.00% | 0.00% |
| RNU | 0.00% | 0.00% | 0.00% | 0.06% | 0.00% | 0.00% | 0.00% | 0.06% |
| snoRNA& scaRNA | 0.00% | 0.00% | 0.00% | 0.01% | 0.00% | 0.00% | 0.00% | 0.01% |
| vault RNA | 0.00% | 0.00% | 0.00% | 0.00% | 0.00% | 0.00% | 0.00% | 0.00% |
| RNY | 0.00% | 0.00% | 0.00% | 0.01% | 0.00% | 0.00% | 0.00% | 0.01% |
| rRNA | 0.04% | 0.75% | 0.47% | 3.32% | 0.04% | 0.75% | 0.47% | 3.32% |
| Combined_tRNA_to_rRNA | 0.05% | 0.75% | 0.48% | 3.42% | 0.05% | 0.75% | 0.48% | 3.42% |
| miRNA | 0.00% | 0.00% | 0.00% | 0.00% | 0.00% | 0.00% | 0.00% | 0.00% |
| premiRNA_only | 0.00% | 0.00% | 0.00% | 0.00% | 0.00% | 0.00% | 0.00% | 0.00% |
| protein coding (PC) mRNA | 0.14% | 0.14% | 0.25% | 0.81% | 0.14% | 0.14% | 0.25% | 0.81% |
| lncRNAs | 0.03% | 0.03% | 0.03% | 0.09% | 0.03% | 0.03% | 0.03% | 0.09% |
| other ncRNA (rest transcriptome) | 0.08% | 0.08% | 0.09% | 0.41% | 0.08% | 0.08% | 0.09% | 0.41% |
| rest hg38 (introns & intergenic) | 1.75% | 1.99% | 2.22% | 7.72% | 1.75% | 1.99% | 2.22% | 7.72% |
| hg38 total | 2.05% | 2.99% | 3.07% | 12.46% | 2.05% | 2.99% | 3.07% | 12.46% |
| no hg38 | 97.95% | 97.01% | 96.93% | 87.54% | 97.95% | 97.01% | 96.93% | 87.54% |
| LSU | 55.01% | 52.51% | 47.88% | 28.79% | 55.01% | 52.51% | 47.88% | 28.79% |
| SSU | 15.43% | 16.32% | 20.78% | 7.71% | 15.43% | 16.32% | 20.78% | 7.71% |
| **Maximum 1 mismatch allowed** | **dLDL** | **dHDL-L** | **dHDL-M** | **dHDL-S** | **dLDL** | **dHDL-L** | **dHDL-M** | **dHDL-S** |
| Total_reads | 46852616 | 44077870 | 33132431 | 17010258 | 46852616 | 44077870 | 33132431 | 17010258 |
| Reads ≥ 15 nt | 43531434 | 41202261 | 30070646 | 9744602 | 43531434 | 41202261 | 30070646 | 9744602 |
| % Reads ≥ 15 nt | 92.91% | 93.48% | 90.76% | 57.29% | 92.91% | 93.48% | 90.76% | 57.29% |
| mtRNA | 0.01% | 0.01% | 0.01% | 0.04% | 0.01% | 0.01% | 0.01% | 0.04% |
| tRNA | 0.04% | 0.03% | 0.03% | 0.15% | 0.04% | 0.03% | 0.03% | 0.15% |
| RN7S | 0.02% | 0.02% | 0.03% | 0.11% | 0.02% | 0.02% | 0.03% | 0.11% |
| RNU | 0.01% | 0.01% | 0.01% | 0.09% | 0.01% | 0.01% | 0.01% | 0.09% |
| snoRNA& scaRNA | 0.01% | 0.01% | 0.02% | 0.03% | 0.01% | 0.01% | 0.02% | 0.03% |
| vault RNA | 0.00% | 0.00% | 0.00% | 0.00% | 0.00% | 0.00% | 0.00% | 0.00% |
| RNY | 0.00% | 0.00% | 0.00% | 0.01% | 0.00% | 0.00% | 0.00% | 0.01% |
| rRNA | 0.18% | 1.83% | 1.01% | 7.20% | 0.18% | 1.83% | 1.01% | 7.20% |
| Combined_tRNA_to_rRNA | 0.25% | 1.89% | 1.10% | 7.57% | 0.25% | 1.89% | 1.10% | 7.57% |
| miRNA | 0.01% | 0.01% | 0.01% | 0.11% | 0.01% | 0.01% | 0.01% | 0.11% |
| premiRNA_only | 0.01% | 0.01% | 0.01% | 0.04% | 0.01% | 0.01% | 0.01% | 0.04% |
| protein coding (PC) mRNA | 2.44% | 2.53% | 3.31% | 9.15% | 2.44% | 2.53% | 3.31% | 9.15% |
| lncRNAs | 0.27% | 0.27% | 0.32% | 0.95% | 0.27% | 0.27% | 0.32% | 0.95% |
| other ncRNA (rest transcriptome) | 0.70% | 0.72% | 0.79% | 2.01% | 0.70% | 0.72% | 0.79% | 2.01% |
| rest hg38 (introns & intergenic) | 5.14% | 5.25% | 6.56% | 18.12% | 5.14% | 5.25% | 6.56% | 18.12% |
| hg38 total | 8.83% | 10.68% | 12.10% | 37.99% | 8.83% | 10.68% | 12.10% | 37.99% |
| no hg38 | 91.17% | 89.32% | 87.90% | 62.01% | 91.17% | 89.32% | 87.90% | 62.01% |
| LSU | 65.73% | 61.33% | 54.23% | 25.28% | 65.73% | 61.33% | 54.23% | 25.28% |
| SSU | 11.17% | 12.41% | 19.81% | 4.89% | 11.17% | 12.41% | 19.81% | 4.89% |

**Supplemental Table S7.** Proteomic result of isolated lipoprotein fractions. Units are presented in ion counts from LC-MS run. Fractions analyzed include intermediate density lipoprotein (IDL); low density lipoprotein (LDL); high density lipoprotein large (HDL-L); high density lipoprotein medium (HDL-M); high density lipoprotein small (HDL-S); albumin (Alb); dense LDL (dLDL); dense HDL-L (dHDL-L); dense HDL-M (dHDL-M); dense HDL-S (dHDL-S); and two quality control HDL samples (QC HDL)

| Protein Record Name | QC HDL | QC HDL | TRLP | DPP | IDL | LDL | HDL-L | HDL-M | HDL-S | Alb | dLDL | dHDL-L | dHDL-M | dHDL-S |
| --- | --- | --- | --- | --- | --- | --- | --- | --- | --- | --- | --- | --- | --- | --- |
| ApoA-I | 71.4594347 | 72.2472703 | 3.07817521 | 0.16986838 | 2.61893599 | 5.92697335 | 61.0591738 | 60.0371084 | 54.9340702 | 3.38092213 | 17.67136 | 46.9133224 | 25.2921792 | 17.9242289 |
| ApoA-II | 13.0523894 | 14.5001001 | 0.9035328 | 0.01787377 | 0.50853463 | 2.19650392 | 12.4738736 | 15.6267512 | 9.51250086 | 0.50559661 | 3.19968945 | 6.15629194 | 4.2319283 | 49.7423747 |
| ApoD | 2.88569482 | 2.02054494 | 0.61491034 | 0.00781197 | 0.00950954 | 0.94639785 | 2.64918422 | 1.77588416 | 2.96933803 | 0.245552 | 0.7324279 | 2.42367887 | 0.43284436 | 2.09636315 |
| ApoB-100 | 2.5115277 | 2.41102051 | 68.3632446 |  | 0.00047055 | 83.9464793 | 3.0067354 | 0.07899673 | 9.63857902 | 12.9971642 | 0.04033846 | 1.59881955 | 43.8652877 | 4.25278958 |
| ApoC-III | 1.83654001 | 1.84899272 | 6.84367271 |  | 4.5041E-05 | 0.90553012 | 4.11859611 | 1.73428363 | 5.58018534 | 0.48012554 | 0.34915905 | 2.57363125 | 0.23602612 | 0.08670169 |
| ApoM | 1.31413772 | 1.27242249 | 0.06484646 | 0.00039667 | 0.00174662 | 0.16728289 | 1.05948226 | 0.97428622 | 0.52169186 | 0.00897979 | 0.10941192 | 0.65375591 | 0.38903078 | 0.7049071 |
| PON1 | 0.8846962 | 0.77287345 | 0.03371408 | 0.05631116 | 0.00792265 | 0.11141057 | 0.18897628 | 0.03255269 | 0.25014921 | 0.07703632 | 0.02551112 | 1.11207878 | 1.28980273 | 1.51852244 |
| ApoE | 0.79370373 | 0.83829397 | 4.96804171 | 0.01703377 | 0.00284812 | 1.21357726 | 1.75648287 | 0.25302035 | 3.39778192 | 0.72587418 | 0.1037715 | 1.55734728 | 1.34960462 | 0.20872863 |
| Immunoglobulin gamma-1 heavy chain | 0.69406944 | 0.53607814 | 0.13607527 | 4.66230446 | 0.30418067 | 0.02170076 | 0.15427606 | 7.77621047 | 0.07860246 | 0.01188013 | 0.07108402 | 0.09527586 | 0.05229897 | 0.16678717 |
| Vitronectin | 0.56817645 | 0.01196711 | 0.00457897 | 0.09684524 | 0.08425732 | 0.03272404 | 0.11132925 | 0.10480702 | 0.17377683 | 0.02897515 | 0.34495533 | 0.25157313 | 0.19781986 |  |
| Serum paraoxonase/lactonase 3 | 0.53026429 | 0.01060923 |  | 1.9512E-05 |  | 8.9001E-05 | 0.02268402 | 0.00150722 | 0.02233172 | 0.00073656 | 0.00270647 | 0.0974054 | 0.51990683 |  |
| Immunoglobulin kappa constant | 0.42879261 | 0.34853375 | 0.16190515 | 2.55016232 | 0.178565 | 0.13784168 | 0.63297685 | 2.81545188 | 0.20885436 | 0.06223995 | 0.07871145 | 0.12928441 | 0.14406339 | 1.33395773 |
| Alb | 0.41903825 | 0.61475702 | 10.623264 | 73.5691204 | 88.7492855 | 0.36167672 | 0.70008397 | 0.78576686 | 1.01543047 | 69.1502842 | 66.6661653 | 2.42524519 | 0.95714717 | 1.83344675 |
| Apolipoprotein F | 0.3005492 | 0.28591246 | 0.01459145 |  |  | 0.06804495 | 0.64286034 | 0.16492403 | 0.30132473 | 0.02175012 | 0.09878254 | 0.70472629 | 0.21626961 | 0.13611397 |
| Apolipoprotein C-II | 0.26030861 | 0.23053781 | 0.99458094 |  | 0.00527014 | 0.09153395 | 0.36841819 | 0.21862643 | 0.33340182 | 0.02943166 | 0.04417792 | 0.11595526 | 0.11647747 | 0.74795433 |
| Apolipoprotein L1 | 0.20406129 | 0.20047566 | 0.07152701 | 0.00364217 | 0.00024767 | 0.19566073 | 0.39190628 | 0.12193016 | 0.43865147 | 0.0693011 | 0.06624635 | 1.3092829 | 0.92499318 | 1.73266771 |
| Immunoglobulin heavy constant gamma 2 | 0.19780074 | 0.08147911 | 0.00988708 | 1.11668089 | 0.06618633 | 0.00036504 | 0.0063063 | 1.02613398 | 0.00431683 |  | 0.00311289 | 0.00533531 | 0.01008365 | 0.02306612 |
| A1AT | 0.17943268 | 0.15741392 | 0.05797123 | 0.44624402 | 0.30361129 | 0.76919296 | 2.80690801 | 0.70395251 | 3.26677804 | 0.71340786 | 1.19936638 | 24.1925773 | 11.4006446 | 1.15179107 |
| Serum amyloid A-4 protein | 0.16720432 | 0.17902243 | 0.06480265 |  | 0.00170421 | 0.02125376 | 0.05463807 | 0.14880794 | 0.06671856 | 0.00345578 | 0.01060218 | 0.03176824 | 0.09420946 | 0.22289047 |
| Complement C3 | 0.16386176 | 0.17154397 | 0.02005765 | 0.81439162 | 0.00983804 | 0.0358603 | 0.2133454 | 0.79138915 | 0.17654775 | 0.01811188 | 0.01534148 | 0.21901058 | 0.20454621 | 0.4219121 |
| Apolipoprotein C-I | 0.12217327 | 0.1714951 | 0.03326562 |  | 0.00585514 | 0.1479586 | 0.06140295 | 0.10125966 | 0.11298723 | 0.01414708 | 0.0174327 | 0.05814835 | 0.11283964 |  |
| Immunoglobulin lambda-like polypeptide 5 | 0.10972038 | 0.11149614 | 0.01573428 | 0.77025078 | 0.01150705 | 0.08779946 | 0.46387941 | 1.02454545 | 0.06565378 | 0.01714417 | 0.00857999 | 0.02492832 | 0.08054697 | 0.07359152 |
| LCAT | 0.10782621 | 0.17989131 |  | 0.00013373 |  | 0.00245499 | 0.23371377 | 0.02276937 | 0.33695124 | 0.00204112 | 0.02233303 | 0.41891979 | 0.0083959 | 0.02293052 |
| Immunoglobulin heavy constant alpha 1 | 0.08534828 | 0.08419158 | 0.04783039 | 1.11057929 | 0.00029128 | 0.093706 | 1.99161417 | 0.42325958 | 0.40907463 | 0.0743856 | 0.0026401 | 0.15975809 | 0.33847587 |  |
| Complement C4-A | 0.0518888 | 0.037554 | 0.0395882 | 0.71578768 | 0.01748777 | 0.06566245 | 0.23915306 | 0.34353453 | 0.24752675 | 0.0515867 | 0.0283138 | 0.32451598 | 0.21115714 | 0.05110052 |
| ApoJ | 0.04800928 | 0.07813057 | 0.04544659 | 0.0944461 | 0.02298793 | 0.25271516 | 0.1408986 | 0.09026423 | 0.32781643 | 0.14192813 | 0.0025818 | 0.81410721 | 0.46267844 |  |
| Haptoglobin-related protein | 0.04670052 | 0.05253206 | 0.00235458 | 0.09890726 |  | 0.23841607 | 0.4033291 | 0.00885601 | 0.7157461 | 0.27076163 | 0.00610945 | 2.39871959 | 3.48239578 | 0.04595761 |
| ApoA-IV | 0.03833568 | 0.03520219 | 0.26856478 | 0.04612939 | 0.03347225 | 0.04498784 | 0.28711559 | 0.11772201 | 0.35665044 | 0.03733422 | 0.05927303 | 0.3134702 | 0.03162207 |  |
| SAA1 | 0.03688629 | 0.03001797 | 0.03474995 |  | 0.00126386 | 0.02857531 | 0.1272709 | 0.13414056 | 0.08241852 | 0.00342978 | 0.05567569 | 0.12789799 | 0.04670985 |  |
| Putative trypsin-6 | 0.03523209 | 0.04672543 | 0.45006884 | 0.02554032 |  | 0.08276619 | 0.04982966 | 0.09218334 | 0.05278231 | 0.03357934 | 0.03432968 | 0.00688988 | 0.2254801 | 13.617445 |
| Immunoglobulin lambda constant 3 | 0.03305908 | 0.02643893 |  | 0.23899128 | 0.00398283 | 0.00366104 | 0.09668379 | 0.26472347 | 0.02991945 | 0.00079155 |  | 0.01117938 | 0.01911329 |  |
| PLTP | 0.03288448 | 0.0310495 | 0.001521 |  |  | 0.04696302 | 0.0643883 | 0.00204913 | 0.08478904 | 0.04972272 | 0.00096208 | 0.19625691 | 0.24454172 | 0.1352816 |
| Prenylcysteine oxidase 1 | 0.03044963 | 0.02955319 | 0.06723443 |  |  | 0.03068691 | 0.07990773 | 0.00586083 | 0.06417899 | 0.00691299 | 0.00145737 | 0.07300047 | 0.04713928 | 0.10497775 |
| Haptoglobin | 0.02248345 | 0.02046141 | 0.03421843 | 1.13602065 | 0.00455528 | 0.02584536 | 0.39591775 | 0.19049478 | 0.76851514 | 0.0434591 | 0.00066055 | 0.03249206 | 0.03308167 |  |
| Antithrombin-III | 0.01805343 | 0.01569522 | 0.0015617 | 0.13707027 | 0.11983428 |  | 0.00054214 | 0.03749781 | 0.00105388 | 0.08019492 | 0.01293321 | 0.09360523 | 0.14946896 |  |
| Immunoglobulin heavy constant gamma 4 | 0.01804109 | 0.02008128 | 0.00369069 | 0.1177858 | 0.00493803 |  | 0.00034048 | 0.14116018 |  |  |  | 0.00114955 |  |  |
| Plasma protease C1 inhibitor | 0.01749645 | 0.02152998 |  | 0.06485211 |  |  | 0.20070677 | 0.25164317 | 0.01341883 |  | 7.8922E-05 | 0.00447407 | 0.03080502 |  |
| Adipocyte plasma membrane-associated protein | 0.01725567 |  |  |  |  |  |  |  |  |  |  |  | 0.00116159 |  |
| Inter-alpha-trypsin inhibitor heavy chain H4 | 0.0139516 | 0.00269219 | 0.04547535 | 0.04073545 | 0.04343156 |  | 0.02839574 | 0.07635574 | 0.02755244 | 5.1167E-05 | 0.00823702 | 0.02628855 | 0.00526595 | 0.00727559 |
| Inter-alpha-trypsin inhibitor heavy chain H2 | 0.01381299 | 0.01275967 | 0.00162739 | 0.1280781 | 0.00078598 |  | 0.38200757 | 0.09733519 | 0.27391894 | 0.0015967 | 0.00025033 | 0.0220895 | 0.00824137 |  |
| Platelet-activating factor acetylhydrolase | 0.01123126 | 0.01531196 | 0.00939981 |  |  | 0.01854649 | 0.0430514 | 0.00397191 | 0.01115464 | 0.00284842 |  | 0.01391487 |  |  |
| Indian hedgehog protein | 0.01049586 | 0.00187034 |  |  |  |  | 0.00160776 | 0.00153838 | 0.00056055 |  | 0.00045971 | 0.00261823 |  |  |
| Protein AMBP | 0.01019684 | 0.00861739 |  | 0.08047821 | 0.0133591 | 0.0014272 | 0.16459827 | 0.0641341 | 0.10690387 | 0.00108036 | 0.00091067 | 0.0107316 | 0.01713551 |  |
| Inter-alpha-trypsin inhibitor heavy chain H1 | 0.01019207 | 0.00937581 |  | 0.04304883 | 0.00130453 | 0.00524921 | 0.37313304 | 0.05272791 | 0.36812053 | 0.00130983 |  | 0.00992935 | 0.00188616 |  |
| Complement factor H | 0.00997016 | 0.01091449 | 0.03895801 | 0.18165198 | 4.8514E-05 | 0.00625285 | 0.54434485 | 0.01052161 | 0.95376679 |  |  | 0.00511167 | 0.00251096 |  |
| Complement C4-B | 0.00983979 | 0.05067988 |  | 0.01221408 | 0.00236867 | 0.00307969 | 0.01599241 | 0.0328077 | 0.01885801 | 0.00272178 | 0.02031111 | 0.06632094 | 0.00253644 |  |
| Immunoglobulin kappa variable 3-20 | 0.00966093 | 0.00722353 |  | 0.06959609 | 0.00264436 |  | 0.01154821 | 0.0803885 | 0.00574172 |  | 0.00072535 |  |  | 0.03104617 |
| Angiotensinogen | 0.00848227 | 0.00100916 | 0.00665133 | 0.06930904 | 0.10631303 | 0.02671601 | 0.0331615 | 0.00598671 | 0.04866948 | 0.07563445 | 0.07262437 | 0.21356426 | 0.2142732 |  |
| Complement C1s subcomponent | 0.00828135 | 0.00658834 |  | 0.0127523 | 0.00506861 | 0.00604932 | 0.00854628 | 0.0835773 | 0.02273587 | 0.00971143 |  | 0.01248478 | 0.02406385 |  |
| Apolipoprotein C-IV | 0.00723106 | 0.00856637 | 0.45432441 |  |  | 0.00793513 | 0.04127315 | 0.00755029 | 0.02857056 |  | 0.00065607 | 0.01541506 | 0.01148081 |  |
| Ceruloplasmin | 0.00713154 | 0.00474468 | 0.00842139 | 0.31727913 | 0.0311121 |  | 0.00701 | 0.24980434 | 0.00116028 |  | 0.00029846 | 0.00432911 | 0.00648567 |  |
| Beta-Ala-His dipeptidase | 0.00697967 | 0.00720841 |  | 0.00108654 | 0.00045141 | 0.00418489 | 0.01455662 | 0.00152788 | 0.01632878 | 0.00416925 | 0.0013546 | 0.17344783 | 0.20591671 | 0.0114877 |
| Alpha-1-acid glycoprotein 2 | 0.00687862 | 0.00668585 | 0.02026269 | 0.10460312 | 0.06266063 | 0.0377169 | 0.05600319 | 0.00320436 | 0.06805605 | 0.08300028 | 0.0101177 | 0.19884789 | 0.44145058 |  |
| Torsin-3A | 0.0065195 |  |  |  |  |  |  |  |  |  |  | 0.00687826 | 0.00798045 |  |
| Keratin, type II cytoskeletal 1 | 0.00525967 | 0.00382602 | 0.0245539 |  |  | 0.02570317 | 0.00359503 | 0.00562653 |  | 0.00981667 |  | 0.02018682 |  | 0.5337975 |
| CETP | 0.00474485 | 0.00261181 |  |  |  |  | 0.0012848 |  | 0.00495488 |  |  | 0.00324492 | 0.00168175 | 0.04191229 |
| Inter-alpha-trypsin inhibitor heavy chain H3 | 0.00462557 | 0.00087865 |  | 0.00611233 |  |  | 0.01021838 | 0.02014589 | 0.00187103 |  |  |  |  |  |
| Phosphatidylinositol-glycan-specific phospholipase D | 0.00450863 | 0.00336401 |  | 0.00076749 |  |  | 0.00428138 | 9.0264E-05 | 0.00468472 |  |  | 0.06315179 | 0.02334052 |  |
| Multimerin-2 | 0.00435984 | 0.00458035 |  |  |  |  | 0.00196696 | 0.00147806 | 0.00146688 |  |  | 0.00292125 | 0.00144294 |  |
| Transthyretin | 0.0042889 | 0.00399393 | 0.00485139 | 0.04468314 | 0.03319195 | 0.11540029 | 0.15777519 | 0.0328381 | 0.20897428 | 0.0989056 | 0.05753293 | 0.56255069 | 0.56347744 |  |
| Immunoglobulin kappa light chain | 0.00421507 | 0.001838 |  | 0.03730108 | 0.00137067 | 0.00119992 | 0.01616861 | 0.08100806 | 0.00272489 |  |  | 0.00031584 |  | 0.10201376 |
| Complement C1r subcomponent | 0.00401867 | 0.00096029 |  | 0.01425857 |  | 0.00394715 | 0.01058286 | 0.00131831 | 0.00963761 | 0.00243907 |  | 0.00313511 | 0.01655911 |  |
| Alpha-2-antiplasmin | 0.00401233 | 0.00387522 | 0.0002667 | 0.02758567 | 0.05071504 | 0.0030652 | 0.0169144 | 0.00177467 | 0.01955355 | 0.00749932 | 0.01485842 | 0.34652046 | 0.21779818 |  |
| Hemopexin | 0.00351775 | 0.00327664 | 0.01978939 | 1.19605002 | 0.97733549 | 0.00094767 | 0.00190766 | 0.0050291 | 0.01029515 | 0.07766859 | 0.02270764 | 0.00876463 | 0.01226504 |  |
| Prolyl endopeptidase FAP | 0.00332039 | 4.8028E-05 |  |  |  |  |  |  |  |  |  |  |  |  |
| Alpha-1B-glycoprotein | 0.00319958 | 0.00255881 | 0.00030524 | 0.13976187 | 0.31219311 |  | 0.00194161 | 0.00294818 | 0.00263483 | 0.02139485 | 0.05906245 | 0.00347202 | 0.00078668 |  |
| CD99 antigen | 0.00319364 | 0.00326566 |  |  |  |  | 0.00530438 | 8.2641E-05 | 0.00560355 |  | 0.00016103 | 0.00240572 |  | 0.00881013 |
| Lumican | 0.00317358 | 0.00291784 |  | 0.00472996 | 0.00631492 |  |  | 0.02576864 |  |  |  |  |  |  |
| Nucleophosmin | 0.00288196 |  |  |  |  |  |  |  |  |  |  |  |  |  |
| Beta-2-glycoprotein 1 | 0.00280783 | 0.00234686 | 0.02547607 | 0.21572507 | 0.30621313 | 0.00386913 | 0.06913533 | 0.0147542 | 0.02944709 | 0.04816855 | 0.03955632 | 0.03918854 | 0.00965727 |  |
| Secretoglobin family 3A member 1 | 0.00252054 | 0.00144334 |  |  |  |  | 0.00217449 | 0.00021512 | 0.00144528 |  |  | 0.00391084 |  |  |
| Insulin-like growth factor-binding protein complex acid labile subunit | 0.00212083 | 0.00157167 |  | 0.01413529 | 0.00804914 |  | 0.00582603 | 0.02805123 | 0.00185195 | 0.00081766 | 0.00379144 | 0.03384117 | 0.0245267 |  |
| Anthrax toxin receptor 1 | 0.00207679 | 0.00046617 |  |  |  |  | 0.00336366 |  | 0.00270448 | 0.00089909 |  | 0.00429119 | 0.00172374 |  |
| Serotransferrin | 0.00205942 | 0.00578098 | 0.32838348 | 4.37880869 | 2.9369743 | 0.0009372 | 0.01140119 | 0.00378328 | 0.00642883 | 0.4103522 | 0.08929905 | 0.01335457 | 0.00382089 |  |
| Cathelicidin antimicrobial peptide | 0.0018135 | 0.00116268 | 0.01334862 |  |  | 0.00033231 | 0.00899629 | 0.00039365 | 0.00041573 |  |  | 0.00101617 |  |  |
| Anthrax toxin receptor 2 | 0.0017989 | 0.00069896 |  |  |  |  | 0.00281437 |  | 0.00901254 | 0.00029077 |  | 0.00093383 |  |  |
| Kininogen-1 | 0.00178471 | 0.0015631 | 0.00241821 | 0.17879438 | 0.20519193 | 0.00814889 | 0.03321303 | 0.0829975 | 0.01514928 | 0.00071824 | 0.00962741 | 0.0159294 | 0.00905124 |  |
| Prosaposin | 0.00163489 | 0.00046398 |  |  |  |  | 0.00235517 | 0.00051625 |  |  |  | 0.0043674 |  |  |
| Fibrinogen beta chain | 0.00161532 | 0.0015036 | 0.03454984 | 0.5296258 |  | 0.3107332 | 0.07365025 | 0.0029011 | 0.11169676 | 0.27719496 |  | 0.00200537 | 0.02927899 |  |
| Keratin, type I cytoskeletal 9 | 0.001607 |  | 0.0061781 |  |  | 0.01488537 | 0.00164714 | 0.00116035 |  | 0.00912835 |  |  | 0.00029885 | 0.25615125 |
| L-selectin | 0.00158298 | 0.00120178 |  |  |  | 0.0017747 | 0.00121927 | 0.0012393 | 0.00796188 | 0.00329219 |  | 0.02966582 | 0.00560348 |  |
| Alpha-2-HS-glycoprotein | 0.00142105 | 0.00018067 | 0.09301246 | 0.15007543 | 0.50813568 | 0.00310063 | 0.00013791 | 0.00263367 | 0.00138265 | 0.08950875 | 0.37732978 | 0.00151303 | 0.0065757 |  |
| Immunoglobulin kappa variable 1-6 | 0.00137941 | 0.00110448 |  | 0.00245778 |  |  | 0.0020267 | 0.01759997 |  |  |  |  |  |  |
| Protein Z-dependent protease inhibitor | 0.00137752 | 0.0008367 |  |  | 0.00041876 | 0.00085044 | 0.00260608 |  | 0.00538699 |  |  | 0.01271332 | 0.01224408 |  |
| Fibrinogen gamma chain | 0.00137362 | 0.00136727 | 0.03159945 | 0.40561435 |  | 0.26866657 | 0.06160834 |  | 0.08277673 | 0.23156259 |  | 0.00074037 | 0.04017532 |  |
| Immunoglobulin heavy variable 5-51 | 0.00136472 | 0.0003374 |  | 0.00520348 |  |  | 0.00227784 | 0.01349086 |  |  |  |  |  |  |
| HLA class I histocompatibility antigen, A-2 alpha chain | 0.00134758 | 0.00052902 | 0.00069537 |  |  |  | 0.00432474 | 0.00033542 | 0.00686374 |  | 0.00056092 | 0.01914419 | 0.00736626 |  |
| Prothrombin | 0.00132428 | 0.00038674 |  | 0.12773112 | 0.22850041 |  |  | 0.0015394 |  | 0.00032568 | 0.0012828 | 0.00047568 |  |  |
| Complement component C6 | 0.00118732 | 0.0008213 |  | 0.0085109 |  |  |  | 0.01649527 |  |  |  |  |  |  |
| Tetranectin | 0.00114855 | 0.00019311 |  | 0.00273312 | 0.00379715 |  |  |  |  |  | 0.00120073 |  |  |  |
| Collagen alpha-1(XVIII) chain | 0.00111855 |  |  |  |  | 0.00360827 | 0.00077885 |  | 0.01472901 | 0.00208186 | 0.00073979 | 0.00618289 | 0.01194208 |  |
| Apo(a) | 0.0010629 | 0.00431017 | 0.5239811 |  | 0.00107216 | 0.46013531 | 0.01529863 | 0.00123681 | 0.15298229 | 0.03439899 |  | 0.00198822 | 0.1520067 | 0.01579798 |
| Glycophorin-A | 0.00105705 | 0.00015606 |  |  |  |  | 0.00072814 |  | 0.00184864 |  |  | 0.00131204 |  |  |
| Fibrinogen alpha chain | 0.00105648 | 0.00778033 | 0.02932915 | 0.27067628 | 0.00031373 | 0.20363593 | 0.03091367 | 0.00140019 | 0.0481012 | 0.10216691 |  | 0.00978763 | 0.02234072 |  |
| Alpha-1-antichymotrypsin | 0.00103056 | 0.0004861 | 0.04837891 | 0.12466692 | 0.1805278 |  | 0.00270987 | 0.00098891 | 0.00270288 | 0.01966738 | 0.02905681 | 0.01338327 | 0.01376069 |  |
| Tissue factor pathway inhibitor | 0.00102096 | 0.00097648 | 0.00093746 |  |  |  | 0.00340414 | 0.00028914 | 0.00322179 | 0.00091219 |  | 0.00378071 | 0.00729364 |  |
| Hemoglobin subunit beta | 0.00099002 | 0.0006255 |  | 0.01162571 |  |  | 0.0039811 | 0.00039456 | 0.00299412 |  |  | 0.000759 | 0.00047982 |  |
| Carbonic anhydrase 6 | 0.00095889 | 0.00121131 |  |  |  |  | 0.00211721 |  | 0.00362579 |  |  | 0.00066014 |  |  |
| Beta-2-microglobulin | 0.00094033 |  |  |  |  |  | 0.0040672 |  | 0.00300615 |  |  | 0.01095346 |  |  |
| A2MG | 0.00093212 | 0.00629054 | 0.09856997 | 0.71852039 |  | 0.00285938 | 0.00864366 |  | 0.01677755 |  |  | 0.01805551 | 0.02967939 |  |
| Immunoglobulin alpha-2 heavy chain | 0.00092981 | 0.00198309 |  | 0.03126204 |  |  | 0.00814408 | 0.01399337 | 0.00334368 |  |  |  | 0.00361851 |  |
| Fibroblast growth factor-binding protein 2 | 0.00090199 | 0.00050219 |  |  |  |  | 0.0006435 |  | 0.0020981 |  |  | 0.00179844 | 0.00030584 |  |
| Complement C2 | 0.0008953 |  |  | 0.00164742 | 0.00277764 |  |  | 0.00028559 |  |  |  |  |  |  |
| Complement C5 | 0.00088125 | 0.00015168 |  | 0.01307023 |  |  |  | 0.01132868 |  |  |  | 0.0090884 |  |  |
| Afamin | 0.00084828 | 0.00308941 | 0.00269208 | 0.04123832 | 0.03524892 |  | 0.00444997 | 0.00123526 |  |  | 0.00054827 | 0.00357606 |  |  |
| Immunoglobulin kappa variable 1-33 | 0.00082581 | 0.00116809 |  | 0.00815521 | 0.0001951 |  | 0.00267423 | 0.01000669 |  |  | 0.00309726 | 0.00023202 |  |  |
| Serum amyloid A-2 protein | 0.00081253 | 0.00280554 | 0.01085632 |  |  |  | 0.01659718 | 0.02160976 | 0.00768106 |  | 0.00641192 | 0.01363557 | 0.00070394 |  |
| Hemoglobin subunit alpha | 0.00080994 | 0.00056326 |  | 0.00838921 |  |  | 0.00182069 | 0.00089925 | 0.00157738 |  |  |  | 0.00010295 |  |
| Actin, cytoplasmic 2 | 0.00077996 | 1.8189E-05 |  |  |  |  |  |  |  |  |  |  |  | 0.22934925 |
| UPF0669 protein C6orf120 | 0.00076717 | 0.00052812 |  |  |  |  | 0.00033244 | 0.00058566 | 0.00088338 |  |  | 0.00134952 |  |  |
| SPARC-like protein 1 | 0.00076607 | 0.00019526 |  |  |  | 0.0244058 | 0.00382358 |  | 0.00283963 | 0.00282375 |  | 0.00792181 | 0.0330378 |  |
| BPI fold-containing family B member 1 | 0.00074079 | 0.00017205 |  |  |  |  | 0.00917175 |  | 0.00419108 |  |  | 0.01322938 | 0.0027562 |  |
| Immunoglobulin heavy constant gamma 3 | 0.00070943 | 0.00647023 |  | 0.06264525 |  |  | 0.06293704 | 0.07675488 | 0.00725755 |  |  | 0.00034169 |  |  |
| Immunoglobulin heavy variable 4-38-2 | 0.00067927 | 0.00054253 | 0.0016313 | 0.0038514 |  |  | 0.00171094 | 0.0181914 | 0.0005647 |  | 2.1502E-05 |  |  |  |
| Selenoprotein P | 0.0006683 | 0.00072439 |  | 0.00221327 |  |  | 0.00170022 | 0.00123541 | 0.00233491 |  |  | 0.00084007 | 0.00314683 |  |
| Extracellular matrix protein 1 | 0.00066819 |  |  | 0.00303625 | 0.00312723 |  |  | 0.0063454 |  |  |  |  |  |  |
| Histidine-rich glycoprotein | 0.00065677 |  |  | 0.06336788 | 0.0128577 |  |  |  |  | 0.00197438 | 0.00057242 | 0.0074694 | 0.00228636 |  |
| Alpha-enolase | 0.00059852 |  |  |  |  |  |  |  |  |  |  |  |  | 0.02219381 |
| Immunoglobulin delta heavy chain | 0.00054401 | 0.00033494 |  | 0.02158469 |  |  | 0.01634707 | 0.00747313 | 0.0302386 |  |  |  |  |  |
| Immunoglobulin kappa variable 1-27 | 0.00052016 | 0.00073206 |  | 0.00017542 |  |  | 0.00080168 | 0.0093837 |  |  |  | 0.0001424 |  |  |
| N-acetylmuramoyl-L-alanine amidase | 0.00051237 | 0.0004069 |  | 0.01087171 | 0.03321872 |  | 0.00196538 | 0.00043989 | 0.00369474 | 0.00084779 | 0.00560873 | 0.01043575 | 0.00275825 |  |
| Complement component C7 | 0.00049612 | 0.00318555 |  | 0.01121873 | 0.00124015 |  |  | 0.12081362 |  |  |  |  |  |  |
| Hepatocyte growth factor activator | 0.00036552 | 2.7403E-05 |  | 0.0001722 |  |  | 0.00129698 | 0.00098611 | 0.00129004 |  |  | 0.0086203 | 0.00392586 |  |
| Immunoglobulin kappa variable 3-11 | 0.00035996 |  |  | 0.0017325 |  |  | 0.00050225 | 0.00516037 |  |  |  |  |  |  |
| Integrin alpha-2 | 0.00033721 | 0.00015474 |  |  |  | 0.00666066 | 0.00020442 |  | 0.00187784 | 0.00189248 |  | 0.00102272 | 0.03031066 |  |
| Gelsolin | 0.00028465 |  | 0.00341355 | 0.08398524 | 0.0925536 |  | 0.00258254 | 0.00108021 | 0.00098532 | 0.00274897 | 0.01535326 | 0.00211573 | 0.00034564 |  |
| Heparin cofactor 2 | 0.00025449 |  | 0.00070037 | 0.01820739 | 0.03419065 |  |  |  |  | 0.00011805 | 0.00287838 |  |  |  |
| Apolipoprotein A-V | 0.00025369 |  |  |  |  |  | 0.00073623 | 0.00016392 | 0.00083503 |  |  | 8.7455E-05 |  |  |
| Complement factor B | 0.00025092 | 0.00065318 | 0.00292571 | 0.11232853 | 0.07444579 |  |  | 0.00066516 |  | 0.00127351 | 0.00264556 | 0.00010411 |  |  |
| Vitamin D-binding protein | 0.00023192 |  | 0.00764871 | 0.43294951 | 0.16077367 |  | 0.00113699 | 0.00220094 |  | 0.31685234 | 0.13112839 | 0.00052999 |  |  |
| Pulmonary surfactant-associated protein B | 0.00022626 | 0.00042171 |  |  |  |  | 0.00050638 |  |  |  | 0.00020313 | 0.00022376 |  |  |
| Vascular cell adhesion protein 1 | 0.00020651 | 1.3661E-05 |  |  |  | 0.00049016 |  |  | 0.00034334 | 0.000957 |  | 0.0049102 | 0.00116737 |  |
| Vitamin K-dependent protein S | 0.000196 | 1.4304E-05 |  | 0.00108304 |  |  |  | 0.00129636 | 0.00159813 |  |  | 0.00532288 | 0.00910075 |  |
| Carboxypeptidase N subunit 2 | 0.00016418 | 0.00013297 |  | 0.01193695 |  |  | 0.01125272 | 0.00183468 | 0.01958723 |  |  | 0.00021144 |  |  |
| Fibulin-1 | 0.00016081 | 1.1385E-05 |  | 0.00213762 |  |  | 0.00337855 | 0.00584523 | 2.4102E-05 |  |  |  |  |  |
| Integrin beta-1 | 0.00015329 |  |  |  |  | 0.02010178 |  |  | 0.00038726 | 0.10212643 |  | 0.00120204 | 0.00935338 |  |
| Adhesion G-protein coupled receptor G6 | 0.00014374 |  |  |  |  | 0.00050417 |  |  | 0.00030013 | 0.00014301 |  |  | 0.00158339 |  |
| Complement component C8 alpha chain | 0.00012568 | 0.00010333 |  | 0.00402891 | 0.00370467 |  | 0.00021684 | 0.00108941 | 0.0001625 |  |  |  |  |  |
| Complement component C9 | 0.0001244 | 0.00065429 |  | 0.01778702 | 0.01695714 | 0.00281797 | 0.01071057 | 0.00458278 | 0.01406005 | 0.00558165 | 0.002864 | 0.10236631 | 0.05261942 |  |
| Immunoglobulin lambda variable 6-57 | 0.00011443 |  |  | 0.00732499 |  |  |  | 0.00283598 |  |  |  |  |  |  |
| Keratin, type I cytoskeletal 10 | 9.5398E-05 | 8.3154E-05 |  |  |  | 0.04237514 | 0.03858652 |  |  | 0.02854689 |  | 0.00078686 |  | 0.13806099 |
| Adiponectin | 9.2214E-05 |  |  | 0.00098834 |  |  | 0.0008541 | 0.00108001 | 0.00097786 |  |  | 0.0001569 | 0.00015954 |  |
| Immunoglobulin kappa variable 2-40 | 9.1718E-05 |  |  |  |  |  |  | 0.00063754 |  |  |  |  |  |  |
| Fibronectin | 8.9451E-05 |  |  | 0.00076663 |  |  | 0.0020315 |  | 0.0035387 |  |  |  |  |  |
| Sulfhydryl oxidase 1 | 7.5458E-05 |  |  |  |  |  |  | 0.00116008 |  |  |  |  |  |  |
| HLA class I histocompatibility antigen, B-46 alpha chain | 6.1833E-05 | 0.00045782 |  |  |  |  | 6.7312E-05 |  | 6.8727E-05 |  |  | 0.00228052 |  |  |
| Protein MENT | 6.0606E-05 | 0.00089665 | 0.00043778 |  |  | 0.00441889 | 0.00561283 | 7.2997E-05 | 0.00521661 | 0.0012545 |  | 0.01534442 | 0.05606032 |  |
| Coagulation factor XIII B chain | 5.335E-05 | 7.2327E-05 |  | 0.00083449 |  |  |  | 0.01510902 |  |  |  |  |  |  |
| Tubulin alpha-1C chain | 5.0292E-05 |  |  |  |  |  |  |  |  |  |  |  |  | 0.02076774 |
| Hyaluronan-binding protein 2 | 4.0011E-05 |  |  | 0.00102304 |  |  | 0.00081941 | 0.00274877 | 0.00121106 |  |  | 0.00139411 |  |  |
| Heat shock 70 kDa protein 1-like | 3.9291E-05 |  |  |  |  |  |  |  |  |  |  |  |  |  |
| LBP | 3.4991E-05 |  |  |  |  |  | 9.5004E-05 |  |  |  |  | 0.00075119 | 0.00044951 |  |
| Coagulation factor VII | 1.336E-05 |  |  |  |  |  |  |  | 0.00012737 |  |  | 0.00082482 |  |  |
| Golgi-associated plant pathogenesis-related protein 1 | 1.1328E-05 |  |  |  |  |  | 0.00017959 |  |  |  |  | 0.00019314 |  |  |
| Coagulation factor X | 6.4825E-06 | 1.8873E-05 |  | 0.00034235 | 0.0003501 |  |  | 0.00029722 |  |  |  |  |  |  |
| Alpha-1-acid glycoprotein 1 |  |  | 0.00113117 | 0.30576603 | 0.21168601 |  |  |  |  | 0.28005026 | 0.02340618 |  |  |  |
| Aminopeptidase N |  | 0.00027421 |  |  |  | 0.00095476 | 0.00066498 |  | 0.03049915 | 0.00050886 |  | 0.01239082 | 0.00762067 |  |
| Attractin |  |  |  | 0.00365393 |  |  |  |  |  |  |  | 0.00021978 |  |  |
| Biotinidase |  |  |  | 0.00175902 | 0.00142227 |  |  |  |  |  |  |  |  |  |
| C4b-binding protein alpha chain |  |  | 0.00522398 | 0.05670781 |  | 0.00067771 |  |  |  |  |  |  | 0.00227083 |  |
| Carboxypeptidase B2 |  |  |  | 0.00096168 | 4.375E-05 |  |  |  |  |  |  |  |  |  |
| Carboxypeptidase N catalytic chain |  |  |  | 0.00089922 |  |  | 0.00019165 |  |  |  |  |  |  |  |
| Cartilage acidic protein 1 |  |  |  |  |  |  |  |  | 0.00127197 |  |  | 0.00227852 |  |  |
| CD5 antigen-like |  |  |  | 0.00199284 |  | 0.00024749 | 0.0026195 |  | 0.00581812 | 0.00030004 |  | 0.01625459 | 0.01811961 |  |
| CD44 antigen |  | 0 |  |  |  |  |  |  |  | 6.794E-05 |  |  |  |  |
| Coagulation factor IX |  |  |  |  | 0.00017842 |  |  |  |  |  |  |  |  |  |
| Coagulation factor XII |  |  |  | 0.00716768 | 0.00018035 |  |  |  |  |  |  |  |  |  |
| Complement C1q subcomponent subunit B |  |  |  | 0.00215886 |  |  |  |  |  |  |  |  |  |  |
| Complement C1q subcomponent subunit C |  |  |  | 0.00216886 |  |  |  | 0.00041725 |  |  |  |  |  |  |
| Complement C1r subcomponent-like protein |  | 0.00056499 |  |  | 0.00049312 |  |  |  | 7.8913E-05 |  |  | 0.00011443 |  |  |
| Complement component C8 beta chain |  |  |  | 0.00451325 | 0.00540416 |  |  | 8.5584E-05 |  |  |  |  |  |  |
| Complement component C8 gamma chain |  |  |  | 0.00102949 | 0.00282781 |  |  | 0.00022634 |  |  |  |  |  |  |
| Complement factor H-related protein 1 |  |  |  | 0.01195415 | 0.00308263 |  |  | 0.00207699 |  |  |  |  |  |  |
| Complement factor H-related protein 4 |  |  | 0.00054656 |  |  |  | 0.0013988 | 0.00107743 |  |  |  |  |  |  |
| Complement factor I |  |  |  | 0.01502325 | 0.00921186 |  |  |  |  |  |  |  |  |  |
| Corticosteroid-binding globulin |  |  |  | 0.0228332 | 0.06589948 |  |  |  |  | 0.00612927 | 0.00462691 |  |  |  |
| CUB and sushi domain-containing protein 2 |  |  |  |  |  |  |  |  |  | 0.007067 |  |  |  |  |
| Cystatin-C |  |  |  | 0.00022328 |  |  |  |  |  |  |  |  |  |  |
| Dipeptidyl peptidase 4 |  |  |  |  |  |  |  |  |  |  |  | 0.00615467 | 0.00177475 |  |
| EGF-containing fibulin-like extracellular matrix protein 1 |  |  |  | 0.00043838 | 0.00034023 |  |  |  |  |  |  | 6.8837E-05 |  |  |
| Endoplasmin |  |  |  |  |  |  |  | 0.00105895 |  |  |  |  |  |  |
| Fetuin-B |  |  |  | 0.00039191 | 0.00037287 |  |  |  |  |  |  |  |  |  |
| Ficolin-3 |  |  |  | 0.00128952 |  |  |  |  |  |  |  |  |  |  |
| Glutathione peroxidase 3 |  |  |  | 0.00125452 | 0.0031033 |  |  |  |  |  |  |  |  |  |
| Glycine N-methyltransferase |  |  |  |  |  |  |  |  |  | 0.18475244 | 0.11060183 |  |  |  |
| Hepatic triacylglycerol lipase |  |  |  |  |  |  |  |  |  |  |  | 0.00068584 | 0.00237495 |  |
| HLA class I histocompatibility antigen, A-24 alpha chain |  |  |  |  |  |  | 0.00143913 |  | 0.00032257 |  |  | 0.00051003 |  |  |
| HLA class I histocompatibility antigen, B-56 alpha chain |  | 0.00090512 |  |  |  |  |  |  |  |  |  | 0.0007642 |  |  |
| Immunoglobulin heavy constant mu |  |  | 0.00976478 | 0.09882981 |  | 0.00371284 |  | 0.00047921 | 0.00108553 |  |  |  | 0.03115565 |  |
| Immunoglobulin heavy variable 3-9 |  |  |  | 0.0014652 |  |  |  | 0.00129975 |  |  |  |  |  |  |
| Immunoglobulin heavy variable 3-33 |  |  |  | 0.00147655 |  |  | 0.0009104 |  |  |  |  |  |  |  |
| Immunoglobulin J chain |  |  |  | 0.00254989 |  |  |  |  |  |  |  |  | 0.01376899 |  |
| Immunoglobulin kappa variable 1-5 |  |  |  |  |  |  |  | 0.00178309 |  |  |  |  |  |  |
| Immunoglobulin kappa variable 1-17 |  |  |  | 0.00045341 |  |  |  | 0.00041829 |  |  |  |  |  |  |
| Immunoglobulin kappa variable 1D-39 |  |  |  |  |  |  |  | 0.001321 |  |  |  |  |  |  |
| Immunoglobulin kappa variable 3D-15 |  | 0.00020302 |  | 0.00140126 |  |  |  | 0.00312738 |  |  |  |  |  |  |
| Immunoglobulin kappa variable 4-1 |  |  |  | 0.00025369 |  |  |  | 0.00142039 |  |  |  |  |  |  |
| Immunoglobulin lambda variable 1-40 |  |  |  | 0.00079731 |  |  |  | 0.00087323 |  |  |  |  |  |  |
| Immunoglobulin lambda variable 1-44 |  |  |  | 0.00124271 |  |  |  | 0.00102006 |  |  |  |  |  |  |
| Immunoglobulin lambda variable 1-51 |  |  |  | 0.00030754 |  |  | 0.0018953 | 0.00015005 |  |  |  |  |  |  |
| Immunoglobulin lambda variable 3-21 |  | 8.8815E-05 |  | 0.04348291 | 0.00066757 |  | 0.01106543 | 0.05695432 | 0.00215161 | 0.00028225 |  |  | 0.00015972 |  |
| Inhibin beta C chain |  | 3.2926E-05 |  |  |  |  | 0.00010523 |  | 0.0001059 | 0.00276016 |  | 0.00088017 | 0.00022103 |  |
| Insulin-like growth factor-binding protein 3 |  |  |  |  |  |  |  | 0.00061748 |  |  |  |  |  |  |
| Integrin alpha-11 |  |  |  |  |  |  |  |  |  |  |  |  | 0.00757037 |  |
| Integrin alpha-M |  |  |  |  |  |  |  |  |  |  |  | 0.00028794 | 0.00220992 |  |
| Intercellular adhesion molecule 1 |  |  |  |  |  |  |  | 9.4326E-05 |  |  |  |  |  |  |
| Kallistatin |  |  |  | 0.00367495 | 0.00392309 |  |  |  |  |  |  |  |  |  |
| Keratin, type II cytoskeletal 2 epidermal |  |  |  |  |  | 0.00088145 |  |  |  | 0.00525177 |  |  |  | 0.17539691 |
| Leucine-rich alpha-2-glycoprotein |  |  | 0.00087335 | 0.01470461 | 0.0088595 |  |  |  |  | 0.0177012 | 0.00391916 |  |  |  |
| Mannan-binding lectin serine protease 1 |  |  |  |  |  |  | 0.01130001 | 0.00137262 | 0.00034367 |  |  |  | 0.00244521 |  |
| Mannan-binding lectin serine protease 2 |  |  |  | 4.6248E-05 |  |  | 9.3073E-05 | 5.8539E-05 | 0.00080063 |  |  | 0.00124582 |  |  |
| Mucosal addressin cell adhesion molecule 1 |  |  |  |  |  |  |  |  | 0.00072951 | 0.00056627 |  | 0.0039884 | 0.00538653 |  |
| Neural cell adhesion molecule 1 |  |  |  |  |  |  | 0.00071845 |  |  |  |  |  |  |  |
| Neurogenic locus notch homolog protein 3 |  |  |  |  |  | 0.01404703 |  |  |  |  |  |  | 0.00602907 |  |
| P-selectin |  |  |  |  |  |  |  |  |  | 4.376E-05 |  | 0.00078114 | 0.00196072 |  |
| Pigment epithelium-derived factor |  |  |  | 0.00980142 | 0.00210971 |  |  |  |  |  |  |  |  |  |
| Plasma kallikrein |  |  |  | 0.01729109 | 0.00012206 |  | 0.02127523 | 0.00401255 | 0.02019632 |  |  | 0.00053713 |  |  |
| Plasminogen |  |  |  | 0.46925484 | 0.05774113 |  |  |  |  | 0.00265721 |  |  | 0.00026983 |  |
| Plexin-B1 |  |  |  |  |  |  |  |  | 0.0059538 |  |  |  |  |  |
| Podocalyxin |  |  |  |  |  | 0.00118657 |  |  |  |  |  |  | 0.00131752 |  |
| Pregnancy zone protein |  |  |  | 0.01254748 |  |  |  |  |  |  |  |  |  |  |
| Properdin |  |  |  |  |  |  | 0.00066388 |  | 0.00140348 |  |  |  |  |  |
| Ras-interacting protein 1 |  |  |  |  |  | 0.01990637 |  |  |  |  |  |  |  |  |
| Ras-related protein Rab-13 |  |  |  |  |  |  |  |  |  | 7.99142932 | 7.76339127 |  |  |  |
| Retinol-binding protein 4 |  |  | 0.00238747 | 0.01967576 | 0.00032297 |  |  |  |  | 0.00075771 |  |  |  |  |
| Serum amyloid P-component |  |  |  | 0.0039022 |  |  |  |  |  |  |  |  |  |  |
| Sex hormone-binding globulin |  |  |  | 0.00183859 |  | 0.00287272 | 0.00450653 | 0.00032573 | 0.00495065 | 0.00081263 | 0.00066285 | 0.04122837 | 0.05523684 |  |
| Tenascin-X |  |  |  |  |  |  | 0.15373887 |  | 0.0031524 |  |  |  |  |  |
| Thyroxine-binding globulin |  |  |  | 0.00515876 | 0.0122742 |  |  |  |  | 0.00091627 | 0.00177705 |  |  |  |
| Toll-like receptor 8 |  |  |  |  |  |  |  |  |  |  |  | 0.00057168 |  |  |
| Transforming growth factor-beta-induced protein ig-h3 |  |  |  |  |  |  |  | 0.00033085 |  |  |  |  |  |  |
| Trypsin-1 |  |  |  |  |  | 0.0004825 |  | 0.00251651 |  | 0.000323 |  |  |  |  |
| Trypsin-3 |  | 0.00070704 | 0.00281292 |  |  |  |  | 0.00095922 | 0.00074374 |  | 8.6276E-05 |  |  | 0.04945087 |
| Vitamin K-dependent protein Z |  |  |  | 8.7464E-05 |  |  |  | 0.00018264 |  |  |  |  |  |  |
| von Willebrand factor |  |  |  |  |  |  |  |  | 0.00177262 |  |  | 0.01805443 | 0.00113283 |  |
| Zinc finger protein 235 |  |  |  |  |  |  |  |  | 0.48179577 |  | 0.00055625 |  |  |  |
| Zinc-alpha-2-glycoprotein |  |  |  | 0.10502497 | 0.02889363 |  |  |  |  | 0.04090353 | 0.01064327 |  |  |  |

**Supplemental Table S8**. Ribonucleic Acid (RNA) sequencing result of all isolated LPP fractions. Fractions analyzed include intermediate density lipoprotein (IDL); low density lipoprotein (LDL); high density lipoprotein large (HDLL); high density lipoprotein medium (HDLM); high density lipoprotein small (HDLS); albumin (Alb); dense LDL (DLDL); dense HDL-L (DHDLL); dense HDL-M (DHDLM); dense HDL-S (DHDLS). Fractions were sequenced four times (1-4). Small RNA (sRNA) sequences were categorized to long non-coding RNA (lncRNA), micro RNA (miRNA), miscellaneous RNA (misc RNA), mitochondrial transfer RNA (mt tRNA), ribosomal RNA (rRNA), small nucleolar RNA (snoRNA), small nuclear RNA (snRNA), transfer RNA, Y RNA (yRNA).

|  | FeatureReads | GenomeReads | MappedReads | TooShortReads | TotalReads | UnannotatedReads | lncRNA | miRNA | misc RNA | mt tRNA | rRNA | snoRNA | snRNA | tRNA | yRNA |
| --- | --- | --- | --- | --- | --- | --- | --- | --- | --- | --- | --- | --- | --- | --- | --- |
| DHDLL_1 | 8902 | 531 | 9433 | 80614 | 10844214 | 10754167 | 47 | 13 | 10 | NA | 8627 | 8 | 21 | 166.0 | 10 |
| DHDLL_2 | 8442 | 504 | 8946 | 75603 | 10625005 | 10540456 | 19 | 17 | 4 | NA | 8244 | NA | 14 | 133.0 | 11 |
| DHDLL_3 | 8971 | 510 | 9481 | 81285 | 10944194 | 10853428 | 40 | 14 | 10 | 1 | 8742 | 1 | 20 | 127.0 | 16 |
| DHDLL_4 | 8704 | 530 | 9234 | 78318 | 10781944 | 10694392 | 19 | 10 | 5 | NA | 8485 | 5 | 24 | 144.0 | 12 |
| DHDLM_1 | 2367 | 544 | 2911 | 75383 | 8202574 | 8124280 | 44 | 18 | 19 | 4 | 2101 | 9 | 29 | 131.0 | 12 |
| DHDLM_2 | 2175 | 496 | 2671 | 67330 | 7966308 | 7896307 | 24 | 15 | 13 | 4 | 1988 | 10 | 22 | 98.0 | 1 |
| DHDLM_3 | 2340 | 558 | 2898 | 72499 | 8177195 | 8101798 | 42 | 18 | 17 | 1 | 2102 | 8 | 24 | 125.0 | 3 |
| DHDLM_4 | 2248 | 511 | 2759 | 71186 | 8002501 | 7928556 | 46 | 21 | 14 | 3 | 2005 | 14 | 27 | 112.0 | 6 |
| DHDLS_1 | 5386 | 1098 | 6484 | 49538 | 3903245 | 3847223 | 86 | 48 | 39 | 7 | 4773 | 49 | 129 | 240.0 | 15 |
| DHDLS_2 | 5128 | 1079 | 6207 | 44414 | 3791522 | 3740901 | 55 | 49 | 43 | 3 | 4560 | 44 | 153 | 203.0 | 18 |
| DHDLS_3 | 5478 | 1081 | 6559 | 49355 | 3919551 | 3863637 | 82 | 69 | 34 | 9 | 4850 | 42 | 133 | 240.0 | 19 |
| DHDLS_4 | 5159 | 1118 | 6277 | 45920 | 3839533 | 3787336 | 76 | 44 | 33 | 9 | 4600 | 36 | 153 | 188.0 | 20 |
| TRLP_1 | 8881 | 3648 | 12529 | 64874 | 3325583 | 3248180 | 343 | 79 | 52 | 7 | 7625 | 37 | 229 | 485.0 | 24 |
| TRLP_2 | 7746 | 3556 | 11302 | 56278 | 3210085 | 3142505 | 309 | 98 | 53 | 15 | 6600 | 46 | 187 | 408.0 | 30 |
| S1TRLP_3 | 8930 | 3912 | 12842 | 64201 | 3344693 | 3267650 | 384 | 105 | 56 | 20 | 7597 | 35 | 209 | 501.0 | 23 |
| TRLP_4 | 8402 | 3511 | 11913 | 58684 | 3258153 | 3187556 | 323 | 95 | 51 | 9 | 7253 | 51 | 189 | 402.0 | 29 |
| DPP_1 | 28847 | 37421 | 66268 | 103296 | 4839948 | 4670384 | 3978 | 794 | 161 | 100 | 18898 | 126 | 1488 | 1137.0 | 2165 |
| DPP_2 | 26767 | 35876 | 62643 | 91671 | 4686005 | 4531691 | 4019 | 751 | 144 | 94 | 17355 | 138 | 1415 | 898.0 | 1953 |
| DPP_3 | 29758 | 38424 | 68182 | 103804 | 4869074 | 4697088 | 4239 | 825 | 160 | 106 | 19301 | 159 | 1583 | 1055.0 | 2330 |
| DPP_4 | 27936 | 36612 | 64548 | 96465 | 4759632 | 4598619 | 4005 | 758 | 167 | 107 | 18096 | 146 | 1479 | 1029.0 | 2149 |
| IDL_1 | 6384 | 1228 | 7612 | 56985 | 11131379 | 11066782 | 99 | 50 | 31 | 7 | 5918 | 22 | 83 | 157.0 | 17 |
| IDL_2 | 6074 | 1121 | 7195 | 50430 | 10823316 | 10765691 | 87 | 53 | 20 | 6 | 5689 | 19 | 67 | 117.0 | 16 |
| IDL_3 | 6460 | 1228 | 7688 | 54688 | 11026443 | 10964067 | 112 | 47 | 26 | 3 | 6011 | 18 | 81 | 138.0 | 24 |
| IDL_4 | 6237 | 1190 | 7427 | 53164 | 10808975 | 10748384 | 111 | 55 | 30 | 5 | 5795 | 21 | 85 | 111.0 | 24 |
| LDL_1 | 1863 | 318 | 2181 | 76658 | 12771435 | 12692596 | 66 | 47 | 20 | 1 | 1363 | 5 | 26 | 311.0 | 24 |
| LDL_2 | 1831 | 307 | 2138 | 69582 | 12521126 | 12449406 | 36 | 54 | 14 | 2 | 1401 | 7 | 21 | 280.0 | 16 |
| LDL_3 | 1918 | 338 | 2256 | 74785 | 12851881 | 12774840 | 78 | 50 | 18 | 1 | 1416 | 8 | 31 | 297.0 | 19 |
| LDL_4 | 1854 | 340 | 2194 | 71932 | 12681911 | 12607785 | 42 | 52 | 14 | 1 | 1414 | 8 | 21 | 278.0 | 24 |
| HDLL_1 | 1261 | 191 | 1452 | 110457 | 10492992 | 10381083 | 49 | 30 | 22 | 1 | 950 | 7 | 10 | 172.0 | 20 |
| HDLL_2 | 1137 | 209 | 1346 | 101943 | 10258703 | 10155414 | 18 | 21 | 17 | 1 | 898 | 7 | 18 | 132.0 | 25 |
| HDLL_3 | 1342 | 204 | 1546 | 110324 | 10583894 | 10472024 | 51 | 20 | 27 | NA | 1034 | 11 | 20 | 143.0 | 36 |
| HDLL_4 | 1208 | 216 | 1424 | 105663 | 10420360 | 10313273 | 21 | 16 | 15 | 1 | 949 | 3 | 17 | 166.0 | 20 |
| HDLM_1 | 1278 | 424 | 1702 | 199710 | 9515472 | 9314060 | 49 | 35 | 25 | NA | 895 | 13 | 29 | 210.0 | 22 |
| HDLM_2 | 1217 | 407 | 1624 | 184434 | 9315811 | 9129753 | 23 | 41 | 22 | NA | 916 | 7 | 18 | 174.0 | 16 |
| HDLM_3 | 1300 | 426 | 1726 | 199753 | 9570386 | 9368907 | 46 | 26 | 18 | 2 | 954 | 11 | 29 | 202.0 | 12 |
| HDLM_4 | 1277 | 417 | 1694 | 192350 | 9434533 | 9240489 | 25 | 32 | 13 | 1 | 928 | 14 | 23 | 220.0 | 21 |
| HDLS_1 | 9141 | 1505 | 10646 | 92145 | 8613712 | 8510921 | 91 | 28 | 34 | 2 | 8554 | 16 | 66 | 339.6 | 10 |
| HDLS_2 | 8604 | 1363 | 9967 | 83640 | 8416830 | 8323223 | 81 | 24 | 41 | 3 | 8031 | 21 | 67 | 324.0 | 12 |
| HDLS_3 | 9393 | 1481 | 10874 | 90719 | 8666621 | 8565028 | 116 | 19 | 39 | 5 | 8722 | 22 | 79 | 368.0 | 23 |
| HDLS_4 | 8686 | 1434 | 10120 | 87538 | 8558018 | 8460360 | 96 | 22 | 42 | 6 | 8098 | 28 | 68 | 303.0 | 23 |
| ALB_1 | 8621 | 1674 | 10295 | 137607 | 6833663 | 6685761 | 139 | 94 | 70 | 11 | 7561 | 56 | 110 | 506.0 | 74 |
| ALB_2 | 7689 | 1608 | 9297 | 122612 | 6620328 | 6488419 | 123 | 88 | 70 | 8 | 6762 | 58 | 122 | 397.0 | 61 |
| ALB_3 | 8693 | 1783 | 10476 | 137665 | 6880607 | 6732466 | 136 | 96 | 98 | 13 | 7556 | 70 | 114 | 531.0 | 79 |
| ALB_4 | 8181 | 1696 | 9877 | 128266 | 6725294 | 6587151 | 103 | 94 | 94 | 10 | 7182 | 40 | 104 | 482.0 | 72 |
| DLDL_1 | 495 | 163 | 658 | 81102 | 11645742 | 11563982 | 36 | 37 | 8 | 1 | 252 | 5 | 12 | 123.0 | 21 |
| DLDL_2 | 431 | 174 | 605 | 74238 | 11164520 | 11089677 | 22 | 15 | 14 | 1 | 224 | 9 | 7 | 110.0 | 29 |
| DLDL_3 | 497 | 206 | 703 | 80242 | 11729019 | 11648074 | 51 | 30 | 9 | NA | 249 | 6 | 16 | 109.0 | 27 |
| DLDL_4 | 438 | 189 | 627 | 76182 | 11364245 | 11287436 | 26 | 19 | 7 | NA | 233 | 4 | 14 | 114.0 | 21 |

**Supplemental Table S9.** Nuclear Magnetic Resonance (NMR) LipoProfile subclass terminologies along with their size ranges (nm) in particle diameter.

| Analyte | Size Range (nm) | Unit |
| --- | --- | --- |
| TRLP | 24-240 | nmol/L |
| Very Large TRLP | 90-240 | nmol/L |
| Large TRLP | 50-89 | nmol/L |
| Medium TRLP | 37-49 | nmol/L |
| Small TRLP | 30-36 | nmol/L |
| Very Small TRLP | 24-29 | nmol/L |
| cLDLP | 19-23 | nmol/L |
| Large cLDLP | 21.5-23 | nmol/L |
| Medium cLDLP | 20.5-21.4 | nmol/L |
| Small cLDLP | 19-20.4 | nmol/L |
| cHDLP | 7.4-13 | μmol/L |
| Large cHDLP | 9.6-13 | μmol/L |
| Medium cHDLP | 8.1-9.5 | μmol/L |
| Small cHDLP | 7.4-8.0 | μmol/L |

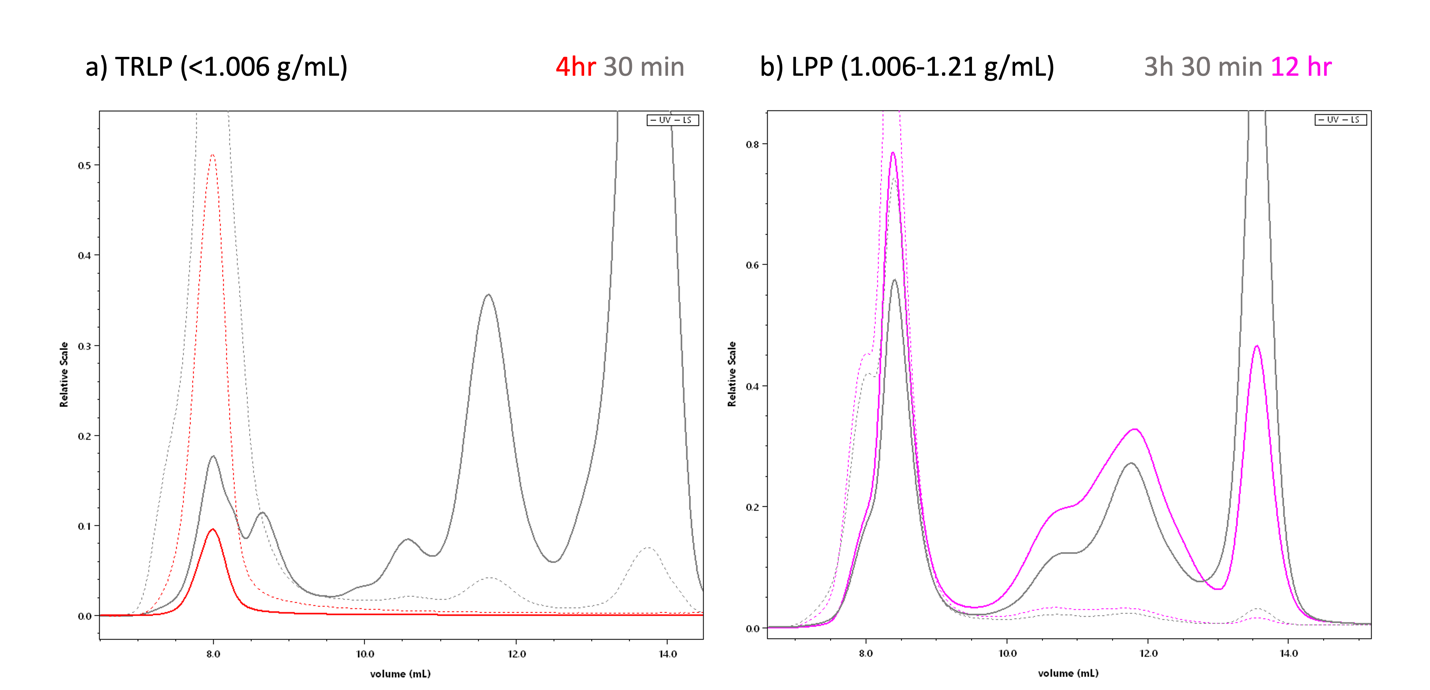

**Supplemental Figure S1.** Chromatograms of lipoprotein particles, where earlier elution corresponds to larger hydrodynamic radii and later elution to smaller radii. (a) TRLP fraction (<1.006 g/mL): The grey chromatogram represents particles spun at 110,000 RPM (656241.5 x *g)* for 30 min, while the red chromatogram represents particles spun for 4 hours under the same conditions. (b) Lipoprotein particles (1.006–1.21 g/mL): The grey chromatogram corresponds to particles spun at 58,000 RPM (182446x *g)* for 3h 30min, while the pink chromatogram represents those spun for 12 hours. All samples were processed using ultracentrifugation with a fixed-angle rotor (k-factor = 13) before injection into the size exclusion chromatography column adapted from Zheng et al. (2021).

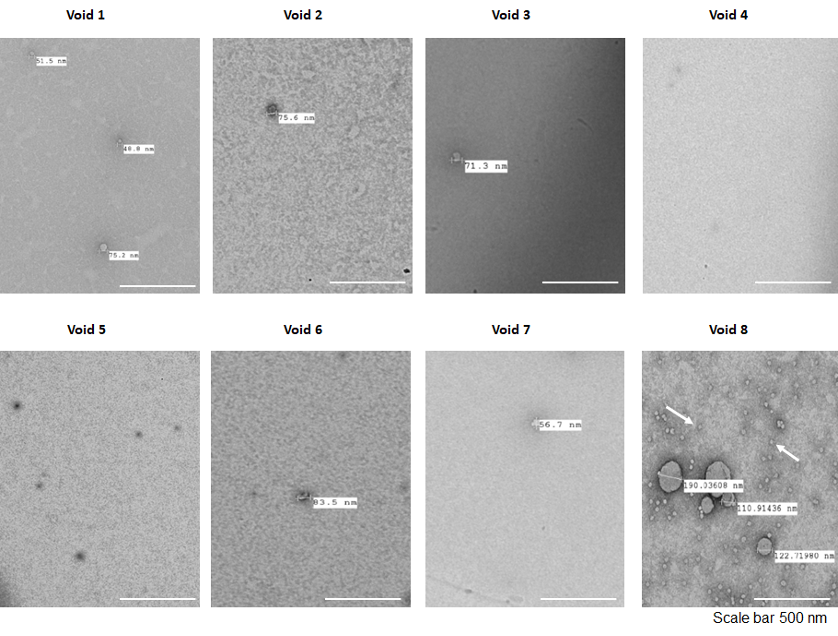

**Supplemental Figure S2.** Transmission electron micrograph (TEM) microscopy images of size exclusion chromatography void volume collected in 1 ml increment leading up to low density lipoprotein (LDL) peak during injection of lipoprotein between density ranges of 1.006-1.21 g/mL. Fraction Void 1 represents the collected fraction furthest away and Void 8 is the closest from the start of the LDL particle elution.
